# Functional Annotation of the Major Histocompatibility Complex Locus

**DOI:** 10.64898/2026.02.01.703124

**Authors:** Lexi R. Bounds, Alejandro Barrera, Maria ter Weele, Siyan Liu, Euphy Wu, Shengyu Li, Revathy Venukuttan, Ruhi Rai, Wancen Mu, Nahid Iglesias, Paola Giusti-Rodríguez, Timothy E. Reddy, Yun Li, Raluca Gordan, Andrew S. Allen, Michael I. Love, Patrick F. Sullivan, Gregory E. Crawford, Charles A. Gersbach

## Abstract

The human major histocompatibility complex (MHC) locus has the greatest density of disease-associations in the human genome, including links to over 100 polygenic disorders. Its complex haplotype structure, rich gene density, and high degree of linkage disequilibrium combine to make deciphering the gene regulatory logic of the MHC locus extremely challenging. Employing complementary high-throughput CRISPR interference (CRISPRi) and activation (CRISPRa) epigenetic screens coupled with single-cell transcriptome profiling across three distinct human cell types, we identified hundreds of new connections between *cis*-regulatory elements (CREs) and their target genes in this locus. These CRE-gene links are largely cell type-specific and act as enhancers. Additionally, some CREs have complex features, including harboring both active and repressive histone marks, lacking chromatin accessibility, targeting multiple genes, or acting as silencers. Computational methods fail to predict a majority of these CRE-gene connections. These findings emphasize the potential for functional perturbation experiments to dissect complex loci and reveal shared and cell type-specific regulatory mechanisms relevant to genomics of complex diseases. Collectively, this study provides a unique resource for understanding the complex regulatory landscape within the MHC locus and supports the need for creating new models that encompass CRE-gene interactions, cell type-specific gene expression, and disease genetics in the noncoding genome.

## Introduction

The vast majority of genetic variation linked to susceptibility to common diseases and other complex traits by genome-wide association studies (GWAS) maps to noncoding regions of the genome, implicating gene regulation rather than protein coding sequences as driving increased risk^1–5^. The human major histocompatibility complex (MHC) locus spans 3.5 Mb on chromosome 6 and contains the greatest number of eQTL SNPs and GWAS catalog significant associations with complex polygenic disorders (> 100 traits) compared to other genomic regions of similar size^2,4,6–11^. Despite its clear involvement in many disorders, the MHC locus is often excluded from functional genomics studies, such as massively parallel reporter assays^12,13^, fine-mapping^14–16^, and molecular trait QTL analyses^17–19^. This exclusion is largely due to analytical challenges that arise from the heterogeneous haplotype structure^20^ and near-perfect linkage disequilibrium (LD) across the MHC locus, which present major obstacles for identifying disease-contributing variants, understanding their mechanisms, and discerning affected genes. Therefore, a targeted effort to link putative noncoding regulatory elements to the genes they regulate within the MHC locus is a crucial step for determining how this region is involved in diverse diseases.

The epigenomic features of the genome have been extensively profiled, collectively identifying more than one million candidate *cis*-regulatory elements (cCREs) across hundreds of human cell types and tissues^21^. The development of CRISPR-based genome and epigenome editing tools has enabled direct perturbation of cCREs to determine the subset that are functionally controlling gene expression (hereafter referred to as CREs). For example, the fusion of a Krüppel associated box (KRAB) domain to the nuclease-inactive Cas9 (dCas9^KRAB^) allows targeted repression of promoters or enhancers to decrease gene expression^22–31^ by depositing H3K9me3, and reducing chromatin accessibility^24^. Conversely, an activating epigenome editor, comprising dCas9 fused to the core domain of the p300 acetyltransferase (dCas9^p300^), increases gene expression when targeted to promoters and distal CREs^32^ by depositing H3K27ac^32^.

By combining high-throughput, pooled screening approaches with CRISPR-based epigenetic activation or repression, it is possible to interrogate large numbers of cCREs in their endogenous context. Previous CRISPR screens, using saturation tiling or targeting of cCREs, have identified and linked CREs to specific genes^26,27,33–38^ or to selectable phenotypes such as cell proliferation^25,39^, response to environmental stimuli^40,41^, or drug treatment^42^. Most CREs from these studies exhibit activating biochemical features (e.g., H3K27ac) and reside in accessible chromatin in the relevant cell or tissue type^43^. These studies typically performed CRISPR screens in a single cell type using CRISPRi loss-of-function perturbations, and employed bulk readouts limited to a particular gene or phenotype. Recent work has coupled CRISPR screens with single-cell transcriptome profiling, allowing for the unbiased connection of CREs with specific genes rather than general phenotypes^34^. This strategy is particularly useful for characterizing disease-associated loci in which the causal genes are uncertain. By comparing screen results across diverse cell types, we can build comprehensive *cis*-regulatory maps that identify both cell type-specific and shared regulatory mechanisms. Moreover, by using both CRISPR interference (CRISPRi) and CRISPR activation (CRISPRa), we can test directionality of regulatory function and reveal the potential for inactive elements to gain function in other cell types or conditions. These studies can define classes of CREs, connect functional noncoding variants to their target genes, and reveal downstream affected phenotypes.

In this study, we focused on understanding the gene regulatory architecture of the challenging and highly complex MHC locus. We coupled noncoding CRISPRi and CRISPRa screens with targeted single-cell transcriptome profiling to build comprehensive *cis*-regulatory maps across diverse cell types. We defined distinctive genomic and epigenomic features of CREs, and identified 142 functionally confirmed CREs and 196 distinct CRE-gene connections within the MHC locus. We demonstrate that functional perturbation experiments identify many CRE-gene connections that are not predicted by currently available computational methods. Our screens in human induced pluripotent stem cells (iPSCs), iPSC-derived neural progenitor cells (NPCs), and K562 erythroid cells revealed shared and cell type-specific CRE-gene connections. We found that GWAS disease-associated variants were enriched in CRE-gene connections, highlighting the importance of characterizing functional cell type-specific regulatory maps. This work establishes a functional genomic resource for the MHC locus that links regulatory elements, noncoding DNA variants, and target genes across three diverse cell types, and represents a critical step toward understanding how this region contributes to many human diseases.

## Results

### Single-cell CRISPR epigenetic screens identify CRE-gene connections within the MHC locus in three diverse cell types

The MHC locus is one of the most complex and gene-dense regions of the human genome (**Supplementary Note 1, Supplementary Figures 1-3, Supplementary Tables 1-3**), creating challenges in understanding gene regulatory logic and the connection of noncoding regulatory elements to their target genes. We performed both inhibition (CRISPRi using dCas9^KRAB^) and activation (CRISPRa using dCas9^p300^) epigenetic editing screens, coupled with single-cell RNA-sequencing (scRNA-seq), to discover CRE-gene connections (defined as a CRE that regulates the expression of a given gene) in three diverse cell types: K562 cells, iPSCs, and NPCs (**Supplementary Note 2**). We designed a library of 12,625 single-guide RNAs (gRNAs) targeting a union set of 537 cCREs (∼20 gRNAs/cCRE) identified via chromatin accessibility in four diverse cell types (K562, iPSCs, NPCs, iPSC-derived Ngn2 neurons) across 3.5Mb of the MHC locus and 44 previously identified enhancers of the pluripotency gene, *POU5F1*^38^, which is in the locus (**Supplementary Figure 4**). We also included positive control gRNAs targeting promoters of lowly and highly expressed genes and negative control gRNAs that do not target sequences within the human genome (nontargeting gRNAs) (**Figure 1A, Supplementary Figure 5**). The library was synthesized in pooled oligo format and cloned into a modified CROP-seq vector^44^ for subsequent recovery of gRNA identities from single-cell transcriptome data.

**Figure 1.**
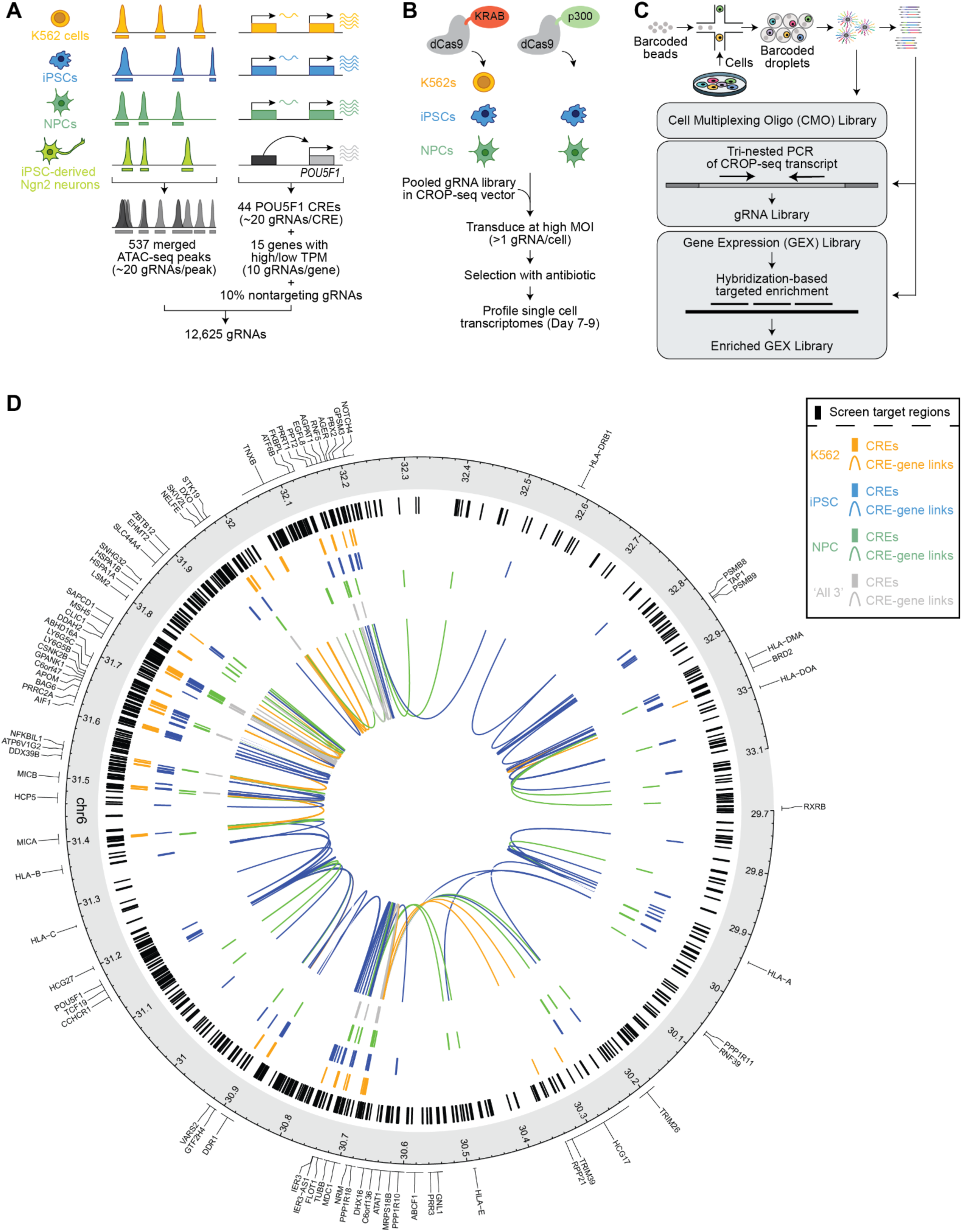
Single cell CRISPR/dCas9 screens identify CRE-gene connections in the MHC locus. **(A)** Overview of gRNA library design targeting a union set of accessible chromatin regions, lowly and highly expressed genes, and previously identified *POU5F1* enhancers^1^ (**Methods**). **(B)** Experimental workflow for single CRISPR/dCas9-based epigenome editing screens. K562 cells, iPSCs, and NPCs expressing dCas9^KRAB^ and iPSCs and NPCs expressing dCas9^p300^ were transduced with the gRNA library such that on average each cell received multiple gRNAs. Single cell transcriptomes were profiled seven to nine days post-transduction. **(C)** Overview of single cell library preparation. **(D)** Circos plot of the targeted cCREs (N=581) and significant CRE-gene connections (N=196). The outermost track indicates the genomic location (GRCh38 coordinates) and the location of each detected gene. The black lines indicate the locations of targeted cCREs in the library. The orange, blue, and green lines indicate CREs paired with a gene in K562 cells, iPSCs, and NPCs, respectively. The innermost track shows a curved line drawn from the CRE to the paired gene with orange, blue, and green shading indicating the connection was observed in K562 cells, iPSCs, and NPCs, respectively. Gray shading indicates the connection was observed in all three cell types.

K562 cells, iPSCs, and NPCs expressing dCas9^KRAB^, and iPSCs and NPCs expressing dCas9^p300^, were transduced with the gRNA library at high multiplicity of infection (MOI), such that on average each cell received multiple epigenetic perturbations in order to maximize library coverage (**Figure 1B, Supplementary Note 2**). Seven to nine days post-transduction, a total of ∼1.2 million high-quality single-cell transcriptomes were profiled using the 10X Genomics platform (**Methods**). To enhance the signal for genes of interest and reduce the required sequencing depth, we performed hybridization-based targeted enrichment of the gene expression libraries for genes within the MHC locus, achieving >79% enrichment of the target genes across all three cell lines (**Figure 1C, Supplementary Figure 6, Methods**). By leveraging both CRISPRi and CRISPRa modalities with targeted enrichment, we effectively characterized the regulation of genes across a wide range of expression levels, overcoming limitations due to data sparsity^45^. Across the five screens, we observed a median of 3-19 unique gRNAs per cell and recovered a median of 62-378 cells per gRNA (**Supplementary Figures 7-8**). We performed differential expression testing for all expressed genes in the MHC locus by comparing all cells in which a given gRNA was observed versus all cells in which one or more different gRNAs were observed (**Methods**).

To assess the quality of the screens, we compared the distribution of gRNA-gene pair p-values using quantile-quantile plots for promoter-targeting positive controls, non-targeting negative controls, and cCRE-targeting gRNAs. Across all five screens, the distributions of both positive control promoter-targeting gRNAs and cCRE-targeting gRNAs separated from the negative control non-targeting gRNAs, indicating that the these two groups led to significant changes in gene expression (**Supplementary Figure 9**). Accordingly, we examined gene expression changes induced by the three most significant gRNAs per cCRE-gene test and found that cCRE-targeting gRNAs consistently led to significantly greater changes than non-targeting negative control gRNAs across all five screens (**Supplementary Figure 10A-E**). Having established that the perturbations for individual gRNAs performed as expected, we calculated a cCRE-level effect size and significance by aggregating individual gRNA effects for each cCRE-gene pair using a robust rank algorithm approach^39^ that directly accounts for the observed effects of non-targeting gRNAs (**Methods**). We defined cCREs with significant effects (FDR <1e-6) on gene expression following perturbation as ‘CREs’ and ‘CRE-gene’ pairs (**Methods, Figure 1D, Supplementary Table 4**). Additionally, all CRE-gene connections within 500 bp of the transcription start site of the paired gene (1.56%-9.04% of all CRE-gene pairs depending on the cell type) were excluded from analysis (as in previous work^46^) since they represent proximal promoter elements that are not the focus of this study.

### Comprehensive identification and validation of CRE-gene connections within the MHC locus

Across all five screens, we connected 24% (142/581) of targeted cCREs to one or more genes, and identified a CRE for 54% (49/90) of genes detected within the MHC locus in these cells (**Figure 2A-B**). In iPSCs, K562s, and NPCs, we identified 96, 59, and 51 CREs, respectively (**Figure 2A**), corresponding to 130, 63, and 69 CRE-gene connections (**Figure 2C, Supplementary Table 4**). The proportion of CREs located within 100 kb, within 1Mb, or over 1Mb away, from the connected gene ranged from 59%-91%, 3%-21%, and 5%-23%, respectively, depending on the cell type (**Figure 2D**). While 73%-93% of CREs regulated a single gene, some CREs were connected to 2-5 genes in each cell type (**Figure 2E**). Similarly, most genes (50%-69%) were connected to 1-2 CREs, although a small number were connected to up to 15 CREs (**Figure 2F**).

**Figure 2.**
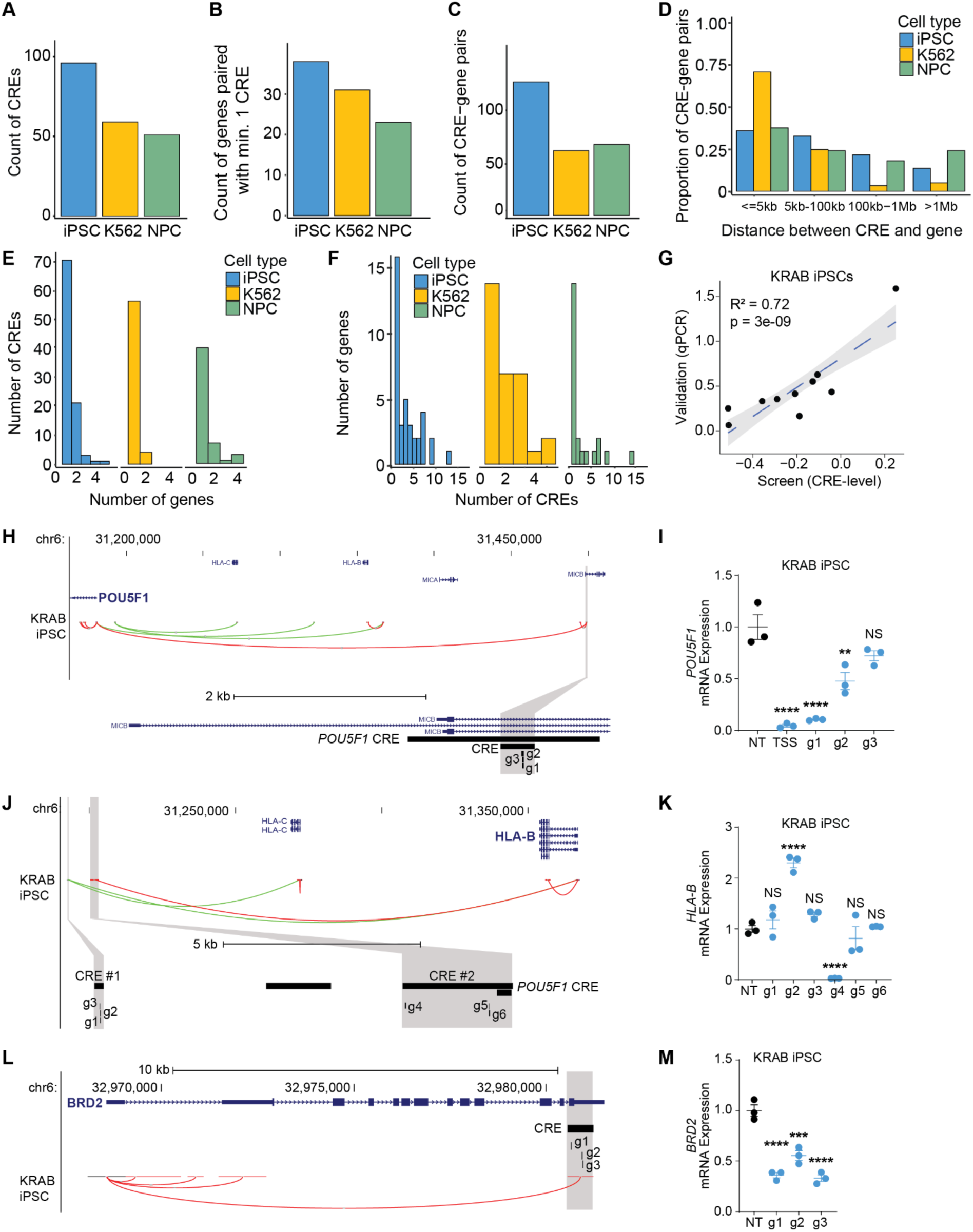
General characteristics of CREs and CRE-gene links. **(A)** Number of CREs identified in each cell type. **(B)** The number of genes connected to at least one CRE. **(C)** Number of CRE-gene pairs identified in each cell type. **(D)** Proportion of CRE-gene pairs that were located within 500bp-5kb, 5kb-100kb, 100kb-1Mb, and >1Mb. **(E)** Histogram of the number of genes connected to a given CRE. **(F)** Histogram of the number of CREs connected to a given gene. **(G)** Scatterplot of the mean change in gene expression measured via RT-qPCR in single gRNA validations versus the effect on gene expression observed in the scRNA-seq screen for the KRAB iPSCs. Pearson correlation and related p-value are denoted in the plot and the blue, dashed line indicates the linear trend. **(H, J, L)** Gene annotations and CRE-gene connections in the KRAB iPSC screen are shown. Red and green curved lines indicate a decrease or increase in gene expression upon perturbation, respectively. The grey shading indicates the CRE(s) and gRNAs (e.g., g1, g2, etc.) tested in validation experiments. **(I, K, M)** Bars represent mean +/- 1 SEM (N=3) and asterisks indicate adjusted p-value from One-way ANOVA followed by Tukey’s HSD post-hoc tests for NT versus each targeting gRNA as follows: ****p<0.0001, ***p<0.001, **p<0.01, *p<0.05. ‘NT’ denotes nontargeting gRNA and ‘TSS’ denotes gRNAs targeting the promoter of the paired gene. **(H)** Example of a long-range interaction between an intronic CRE and *POU5F1*. Gene annotations and CRE-gene connections in the KRAB iPSC screen are shown. The zoomed view below shows the overlap of the CRE with the *MICB* intron and the location of the individual gRNAs used in validation experiments. **(I)** Change in *POU5F1* expression measured via RT-qPCR following delivery of individual gRNAs targeting the CRE in **(H)** to KRAB iPSCs. **(J)** Example of two nearby CREs that have opposing effects on HLA-B expression upon perturbation. The zoomed view below shows the locations of individual gRNAs used in validation experiments. **(K)** Change in *HLA-B* expression measured via RT-qPCR following delivery of individual gRNAs targeting the CREs in **(J)** to KRAB iPSCs. **(L)** Example CRE that overlaps a 3’ UTR region of the regulated gene, *BRD2*. **(M)** Change in *BRD2* expression measured via RT-qPCR following delivery of individual gRNAs targeting the CREs in **(L)** to KRAB iPSCs.

Previous noncoding CRISPRi screens suggest that regulatory elements closer to target genes exert stronger effects than those farther away^34^. In our CRISPRi screens, perturbations of CREs <100 kb of the target gene predominantly led to decreases in gene expression. Interestingly, CRISPRi perturbations of CREs >100 kb mostly led to increased gene expression, perhaps due to trans effects, disruption of boundary elements of topologically associating domains that lead to inappropriate enhancer activation of the target gene, or potential statistical artifacts due to the sensitivity of probe-based enrichment (**Supplementary Figure 11A-C**). For CRISPRa screens, we detected increases in target gene expression at all distances from the CRE. (**Supplementary Figure 11D-E**).

To validate the screen results, we delivered individual gRNAs corresponding to ten CRE-gene connections (9 with decreased gene expression and 1 with increased gene expression in CRISPRi screens) to dCas9^KRAB^ iPSCs and measured the change in target gene expression using RT-qPCR (**Supplementary Tables 5-6**). These CREs overlap diverse genomic annotations, including intergenic, intronic, promoters, and 3’ untranslated regions (UTRs). Individual validation of all ten connections correlated with screen results (**Figure 2G**). For example, gRNAs targeting an intronic CRE in *MICB* led to 50%-90% reductions in *POU5F1* mRNA expression located 6 kb away (**Figure 2H-I**). This CRE is located within a 2 kb *POU5F1* enhancer previously identified in human embryonic stem cells^38^. We also observed that perturbing some promoters resulted in altered expression of other genes, consistent with recent models of promoters predicted to act as enhancers^47^. As a representative example, targeting the promoter of *TAP2* with dCas9^KRAB^ or dCas9^p300^ led to decreased or increased expression, respectively, of *PSMB8* located 5 kb away (**Supplementary Figure 12A**) and was independently validated by RT-qPCR (**Supplementary Figure 12B**). We also found an example of two intergenic CREs located within ∼7 kb of each other that were validated as having opposing regulatory effects on the same gene, resulting in either a 2.3-fold increase (gRNA-2 in ‘CRE1’) or a 97% decrease (gRNA-4 in ‘CRE2’) in *HLA-B* mRNA expression (**Figure 2J-K**). Lastly, perturbation of the 3’ UTR of the *BRD2* gene resulted in a ∼50% decrease in *BRD2* mRNA expression (**Figure 2L-M**).

### Activating and repressive perturbations offer complementary insights into CRE activity for lowly and highly expressed genes

We compared the CRISPRi and CRISPRa experiments and found that ∼90%-95% of CREs were unique to either CRISPRi or CRISPRa perturbations in iPSCs (**Figure 3A**) or NPCs (**Figure 3B**), while only 5%-10% of CREs were detected in both CRISPRi and CRISPRa in the same cell type. We found a similar amount of CRE-gene pairs that were either unique or shared in CRISPRi or CRISPRa in iPSCs (**Figure 3C**) or NPCs (**Figure 3D**). We hypothesized that this lack of overlap between CRISPRi and CRISPRa perturbations in the same cell type was due to lowly expressed genes being difficult to detectably repress further with CRISPRi, but more amenable to activation by CRISPRa. To test this, we examined genes linked to a CRE in either or both screens (**Figure 3E-F**). Indeed, genes connected to a CRE only in the CRISPRi screen had higher expression levels than those linked to a CRE exclusively in the CRISPRa screen in both iPSCs (**Figure 3G**) and NPCs (**Figure 3H**). In addition, genes connected to a CRE in both CRISPRi and CRISPRa screens had intermediate expression levels, indicating they could be perturbed in either direction. Given the opposing biochemical marks deposited by dCas9^KRAB^ and dCas9^p300^, we anticipated that CRE-gene pairs identified by both methods would have opposite effects on gene expression. Indeed, perturbation of CRE-gene pairs identified by both CRISPRi and CRISPRa led to opposite changes in gene expression with dCas9^KRAB^ and dCas9^p300^ **(Supplementary Figure 13A-B)**.

**Figure 3.**
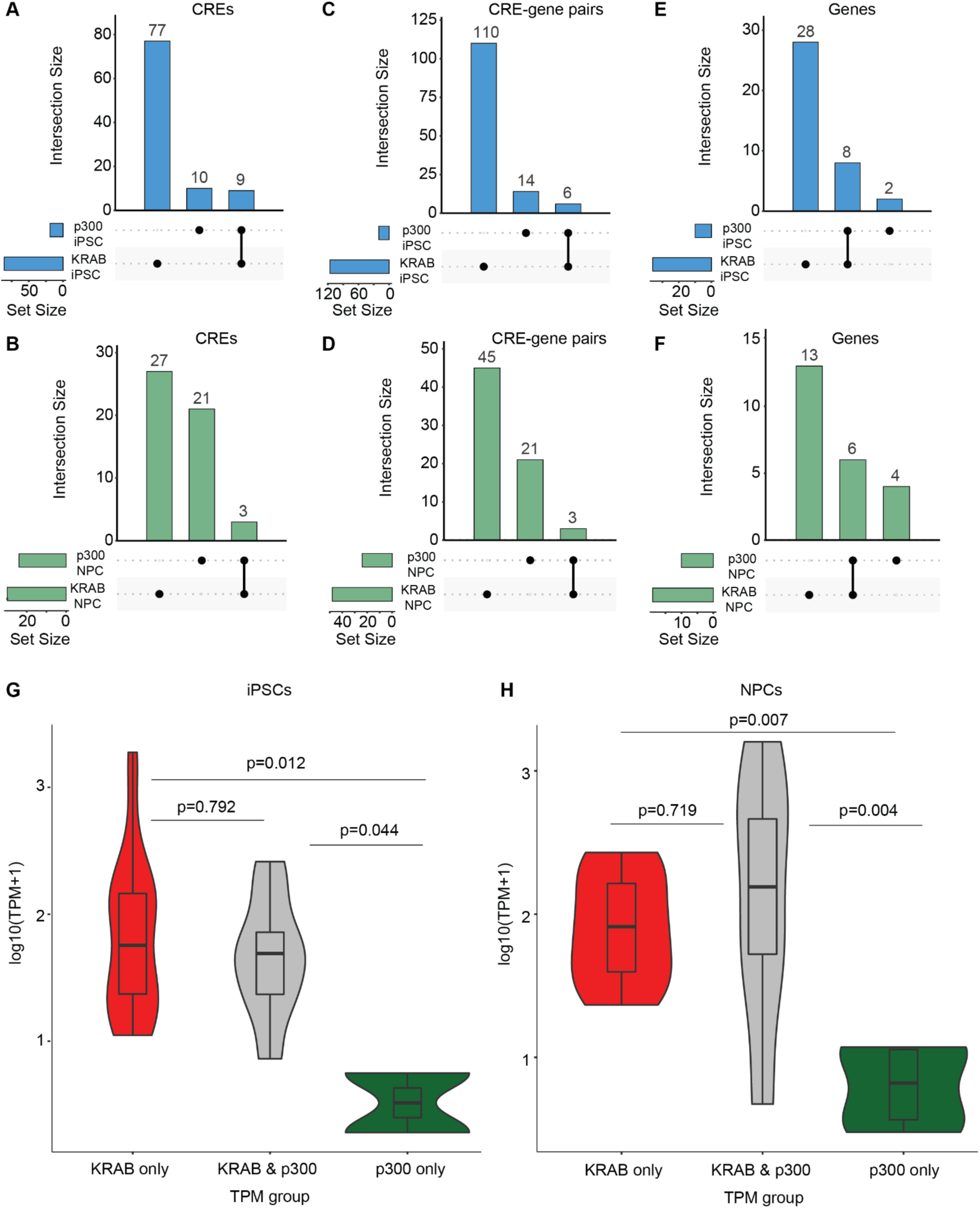
CRISPRi and CRISPRa identify mostly unique regulatory mechanisms. Upset plots of CREs, CRE-gene pairs, and genes connected to a CRE for CRISPRi and CRISPRa screens in iPSCs **(A,C,E)** and NPCs **(B,D,F)**. **(G-H)** Violin and boxplots comparing the basal expression level of genes connected to a CRE with only CRISPRi (red), with only CRISPRa (green), or with both perturbation modalities (grey) in **(G)** iPSCs **(H)** and NPCs. The p-values from Tukey’s post hoc test are shown above each comparison in the plot.

Since our gRNA library targeted a union set of ATAC-seq peaks across four cell types, in each experimental cell type we targeted a combination of open and closed chromatin regions. Therefore, we next compared the effect on gene expression for CRISPRi and CRISPRa perturbations of cCREs that do/do not overlap ATAC-seq peaks in a given cell type. For iPSCs and NPCs, perturbing CREs within accessible chromatin using dCas9^KRAB^ led to a greater median decrease in gene expression compared to targeting non-accessible chromatin (**Supplementary Figure 13C-E**). In contrast, perturbations with dCas9^p300^ led to similar changes in gene expression regardless of whether the chromatin was accessible or inaccessible (**Supplementary Figure 13F-G**). This indicates that epigenetic repression via dCas9^KRAB^ may primarily act on regulatory elements that are accessible, while epigenetic activation via dCas9^p300^ can similarly impact regulatory elements regardless of their chromatin accessibility status. This is consistent with previous work showing the ability of dCas9 to access heterochromatic regions^48^.

### Epigenomic and genomic features define subclasses of CREs

We next characterized the CREs for genomic and epigenomic features. Previous CRISPR screens of the noncoding genome suggest that regulatory elements are predominantly located in accessible chromatin regions^35,38,43,46^. Since our set of test cCREs encompassed the union of accessible chromatin regions across four cell types, we could evaluate if any of the CREs were inaccessible in a cell type in which they were experimentally linked to a gene. Consistent with previous studies, nearly all CREs in K562 cells (95%) were found in accessible chromatin (**Figure 4A**) but for iPSCs and NPCs, only 70% and 80% or 63% and 42% of CREs identified with CRISPRi or CRISPRa, respectively, overlapped with accessible regions, a difference potentially due to the large number of ATAC-seq peaks unique to K562 cells included in the gRNA library design (**Supplementary Figure 4**). There was an enrichment for open chromatin in CREs identified in K562 cells and NPCs with CRISPRi relative to all targeted cCREs, with iPSC CREs trending towards enrichment. Comparing CREs with ATAC-seq peaks across cell types revealed that a large fraction of CREs in iPSCs and K562s are exclusively accessible in that same cell type (**Figure 4B-D, Supplementary Figure 15A**). Overall, we observed that while most CREs overlap accessible chromatin regions, targeting inaccessible regions can also impact gene expression^48^. However, this could also be due to chromatin accessibility being below the threshold for peak calling.

**Figure 4.**
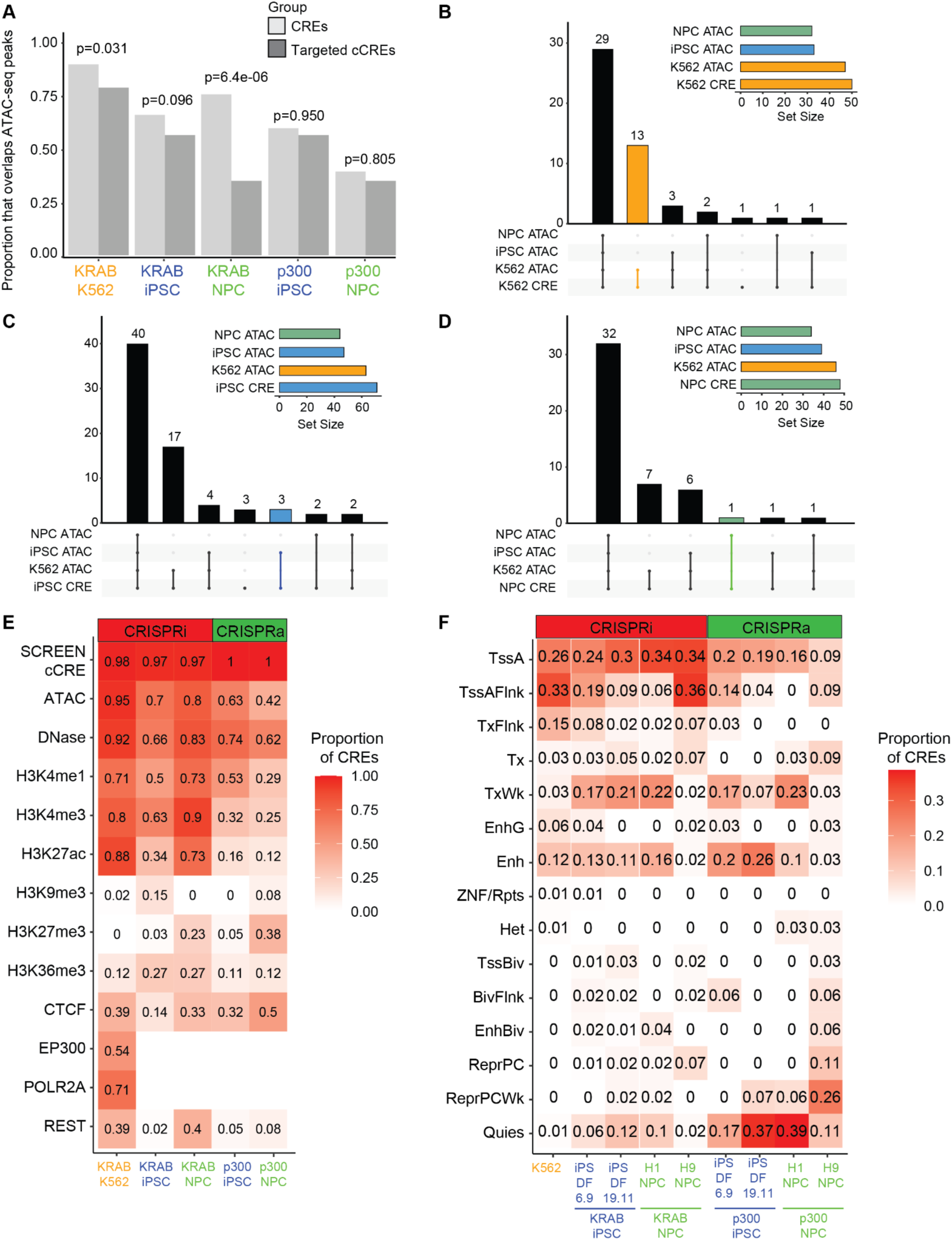
Epigenomic and genomic features define subclasses of CREs. **(A)** Proportion of cCREs included in the gRNA library design and the CREs in each cell type that also overlap an ATAC-seq peak in that cell type. P-value from the chi-squared test comparing the proportion that does or does not overlap an ATAC-seq peak is shown above each pair of bars. **(B-D)** Upset plots of CREs and ATAC-seq peaks in all three cell types and targeted cCREs in the library design. The total set sizes are shown in the upper right of each panel. The intersection size is shown in the main plot. Bars colored by orange, blue, and green, denote the intersections of cell type-specific CREs and cell type-specific ATAC-seq peaks. **(E)** Proportion of CREs in K562s, iPSCs, and NPCs, that overlap various genomic and epigenomic annotations in the corresponding cell types. **(F)** Proportion of CREs that overlap chromHMM chromatin state annotations in the same or similar cell types. ‘iPS DF 6.9’ and ‘iPS DF 19.11’ are iPSC lines and ‘H1 NPC’ and ‘H9 NPC’ are H1 ESC-derived NPCs and H9 ESC-derived NPCs, respectively. Y-axis labels are abbreviated as follows: TssA = Active TSS, TssBiv = Bivalent/poised TSS, BivFlnk = Flanking bivalent TSS/enhancer, EnhBiv = Bivalent enhancer, ReprPC = Repressed polycomb, ReprPCWk = Weak repressed polycomb, Quies = Quiescent/low, TssAFlnk = Flanking TSS, TxFlnk = Strong flanking transcription, Tx = Strong transcription, TxWk = Weak transcription, EnhG = Genic enhancer, Enh = Enhancer, ZNF/Rpts = ZNF genes and repeats, Het = Heterochromatin. **(E-F)** The proportion of CREs is noted in each cell.

In agreement with previous studies^43^, the majority of CREs overlapped with regions of accessible chromatin and/or regions marked by active histone marks (H3K4me1/3, H3K27ac), RNA Pol II, CTCF, and EP300 binding, consistent with these features being associated with enhancers^49^ (**Figure 4E**). Notably, CREs identified using CRISPRi have a greater overlap with H3K27ac peaks compared to CREs identified using CRISPRa, supporting that 1) CRISPRi works primarily on marked enhancers (i.e., inactive regions cannot be further silenced), and 2) CRISPRa can activate silent regions that have been identified as enhancers in other cell types (**Figure 4E, Supplementary Figure 14, Supplementary Figure 15B, Supplementary Table 7)**. K562 and NPC CREs were enriched for REST ChIP-seq peaks, supporting the role of REST in repressing neuronal genes in non-neuronal cell types and in NPCs before final differentiation into neurons^50,51^ (**Supplementary Figure 15B**). To determine the chromatin states that define the CREs in each cell type, we intersected the CREs with chromHMM annotations in the same or similar cell types (**Figure 4F, Supplementary Figure 15C, Supplementary Table 8**). Across all three cell types, CREs identified by CRISPRi were enriched for transcriptionally active regions (TSS and flanking regions) (**Supplementary Figure 15C**). In contrast, CREs identified by CRISPRa were enriched in quiescent regions (**Supplementary Figure 15C**). Collectively, these results support that accessible chromatin, H3K4me1/3, and H3K27ac, are key defining features of CREs, in accordance with previous work^52,53^. However, these findings also reveal the complex combinations of features that can mark CREs, the regulatory function of regions predicted to be inactive, cell type-specific differences in the regulatory landscape, and the distinct CRE features identified by different perturbation modalities.

### Identification of putative silencer regulatory elements

CRISPRi and CRISPRa perturbations that result in gene expression changes that are opposite of what is expected (e.g. CRISPRi resulting in increased gene expression, or CRISPRa resulting in decreased gene expression), are potentially indicative of silencer CRE function^26,54^. We identified a number of possible silencer CREs, defined as CREs only linked to increased or decreased expression with CRISPRi or CRISPRa, respectively, in K562 cells (N=2), iPSCs (N=14, **Figure 2J**, ‘CRE #1’), and NPCs (N=4) (**Supplementary Figure 16A**). All silencer CREs were unique to each cell type except for one silencer CRE shared between iPSCs and NPCs (**Supplementary Figure 16B**). Consistent with previous work showing that candidate silencer CREs are less accessible than active enhancers^55^, silencer CREs often lacked chromatin accessibility and activating histone marks (**Supplementary Figure 16C-D**, **Supplementary Table 9**). Accordingly, NPC silencer CREs were enriched for ‘Repressed polycomb’ (ReprPC) chromatin states and iPSC silencer CREs were enriched for ‘Weak transcription’ (TxWk) chromatin states (**Supplementary Figure 16E**, **Supplementary Table 10**). These results significantly expand the set of reported and functionally interrogated putative silencer CREs, which have not been studied to the same extent as enhancer CREs.

### Benchmarking functional perturbation results against various prediction strategies

Several methods have been developed to predict CRE-gene interactions, including the nearest gene, the Activity-By-Contact (ABC) model^46^, the ENCODE Enhancer-2-Gene (E2G) model^56^, the EnhancerAtlas 2.0^57^, and chromatin looping methods such as HiCAR^54^. We investigated whether CRE-gene connections identified through direct perturbation in this study would be predicted by such methods. We found that a simple ‘nearest gene’ approach predicted 52%-94% of all CRE-gene links within a given cell type (**Figure 5A**). However, CREs also bypassed many nearby genes to regulate more distant targets (**Figure 5B**). Interestingly, other methods designed to predict longer range interactions predicted significantly fewer CRE-gene pairs than we observed (**Figure 5A**). Combining all prediction methods with the nearest gene approach yielded only a modest improvement over the nearest gene approach alone (**Figure 5A**). Overall, the E2G model performed better than ABC, EnhancerAtlas, and Hi-CAR, and showed better performance for CRISPRi than CRISPRa and specifically for K562 cells, which could be due to model training on K562 enhancers^56^ (**Supplementary Figure 17**). We observed no correlation between the ABC Score and the significance of the CRE-gene links across all five experiments (**Supplementary Figure 18**). The distance between the CRE and its target gene was greater for CRE-gene pairs that were not predicted by any method than for the CRE-gene links that were predicted (**Figure 5C**). Additionally, these unpredicted CRE-gene pairs had a lesser absolute change in gene expression following CRISPRi perturbation, supporting that their lack of prediction may be due to more subtle regulatory effects (**Figure 5D**). Overall, these findings indicate that functional experimental perturbations identify CRE-gene connections in this locus that are largely missed by existing computational methods and chromatin looping assays. This observation could be specific to the MHC locus and its high gene density and complex regulatory logic and may not generalize to the more typical loci on which these models were developed. We also cannot rule out the possibility that these long range interactions are secondary downstream effects.

**Figure 5.**
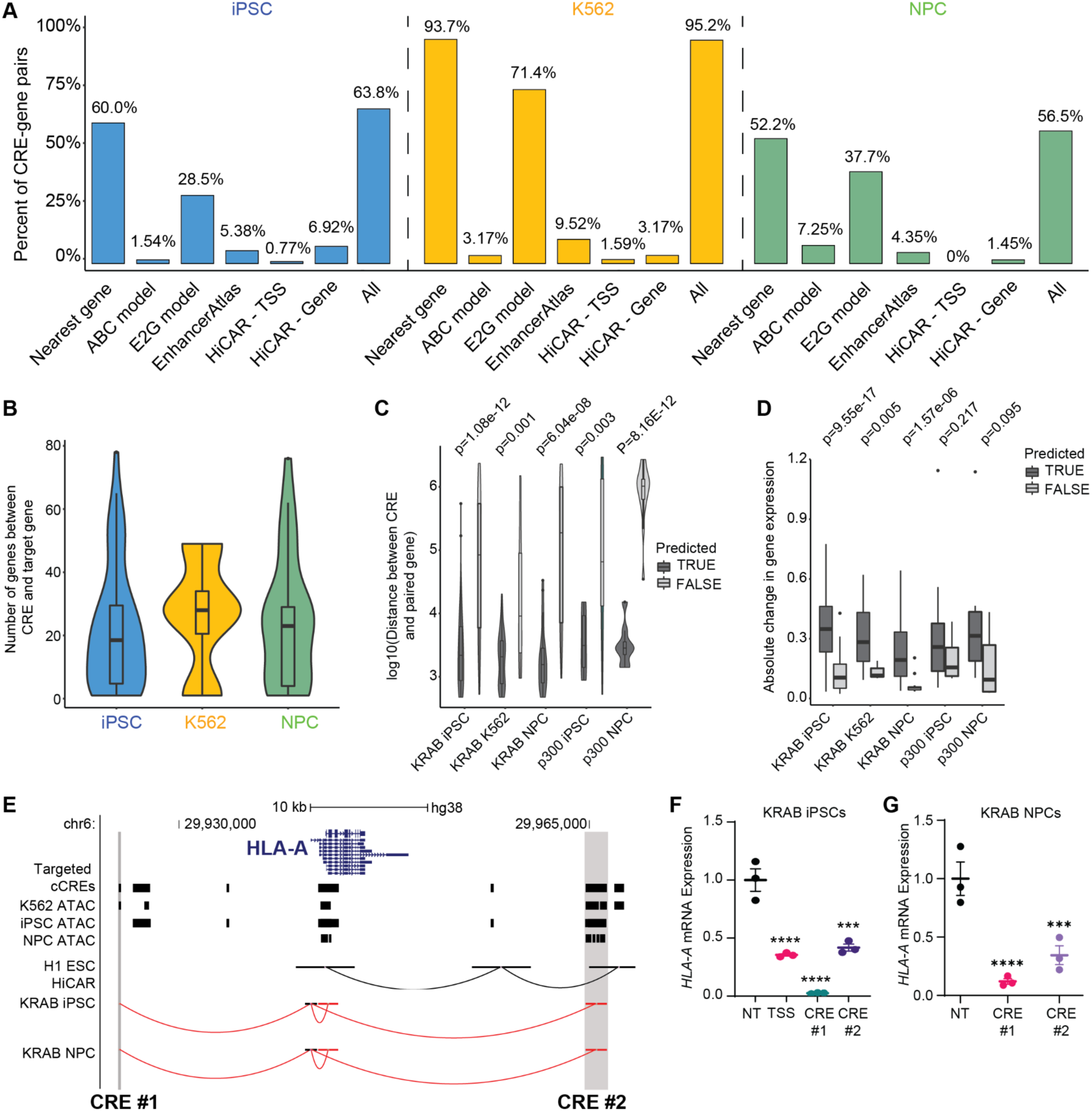
Benchmarking functional perturbation results with common regulatory prediction methods. **(A)** Proportion of CRE-gene pairs identified in the screen that were also predicted by various methods (left to right: nearest gene, ABC model, E2G model, EnhancerAtlas, HiCAR - TSS, and HiCAR - Gene). ‘HiCAR - TSS’ and ‘HiCAR - Gene’ indicate the chromatin loop is between the CRE and the TSS or the gene body of the paired gene. ‘All’ indicates the union set of CRE-gene pairs across all methods. **(B)** Number of genes between CREs and paired genes in each cell type for pairs with at least one gene between the CRE and paired gene. The y-axis is log10-transformed for visualization. **(C)** The log10-transformed distance between the CRE and paired gene for CRE-gene pairs predicted (dark grey) or not predicted (light grey) using all methods except for the ‘Nearest gene’ method. P-values from the Wilcoxon test using log10-transformed distance values are noted in the panel. **(D)** Absolute change in gene expression of paired genes for CRE-gene pairs predicted (dark grey) or not predicted (light grey) using all methods. P-values from the Wilcoxon test using log10-transformed distance values are noted in the panel. **(E)** Example CREs that regulate *HLA-A* expression in iPSCs and NPCs and have different genomic and epigenomic features. Gene annotation, gRNA library target regions, ATAC-seq peaks in K562 cells, WTC11 iPSCs, and WTC11 NPCs, chromatin contact loops in H1 ESCS measured via HiCAR^2^, and CRE-gene connections in the KRAB iPSC and KRAB NPC are shown from top to bottom, respectively. For CRE-gene connections, red and green curved lines indicate a decrease or increase in gene expression upon perturbation, respectively. The grey shading indicates the CREs tested in validation experiments. **(F-G)** Change in *HLA-A* expression measured via RT-qPCR following delivery of individual gRNAs targeting the CREs in **(D)** to **(F)** KRAB iPSCs and **(G)** KRAB NPCs. Bars represent mean +/- 1 SEM (N=3) and asterisks indicate adjusted p-value from One-way ANOVA followed by Tukey’s HSD post-hoc tests for NT vs each targeting gRNA as follows: ****p<0.0001, ***p<0.001, **p<0.01, *p<0.05.

As a representative example of these differences in prediction methods, we focused on two CREs near the *HLA-A* gene in iPSCs and NPCs that displayed differences in chromatin accessibility and 3D chromatin conformation (**Figure 5E**). CRE #1 overlaps an ATAC-seq peak only in K562 cells whereas CRE #2 is accessible in all three cell types. Chromatin looping as determined by HiCAR was only observed between CRE #2 and the *HLA-A* promoter (via an intermediate intergenic region) in a cell type similar to iPSCs (human embryonic stem cells) but not in NPCs. CRE #2 was predicted to regulate *HLA-A* expression by only two methods (nearest gene and E2G), whereas CRE #1 was connected to *HLA-A* only by the nearest gene method. In our CRISPRi screens, perturbation of either CRE decreased *HLA-A* expression in both iPSCs and NPCs. Using single gRNA validations, we confirmed that both CREs influenced *HLA-A* expression in KRAB iPSCs (**Figure 5F**) and KRAB NPCs (**Figure 5G**). Of the ten validated CRE-gene connections in KRAB iPSCs (**Figure 2G**), three were not predicted by any method, six were predicted by a single or by only two methods, and one was predicted by three methods (**Supplementary Table 11**). Together, this supports the need for functional perturbation experiments rather than relying solely on any of the currently established prediction methods.

### Shared CREs are marked by accessible chromatin and are located closer to promoters of target genes

We next investigated the 18 CREs shared across all three cell types (**Figure 6A-B**). These 18 CREs were involved in 44 total CRE-gene connections (**Figure 6C**). The effect sizes of these 18 shared CRE-gene connections were correlated and consistent in direction within perturbation modalities across cell types (**Figure 6D**). Over 77% of the shared CREs overlapped accessible chromatin regions in each cell type (**Figure 6E**) and shared CREs were located closer to the promoters of their target genes (**Figure 6F**). Notably, the connected genes did not show higher expression levels (**Figure 6G)** compared to links identified in only one or two cell types. An illustrative example of a shared CRE-gene connection is an intronic region of *CLIC1* which is highly expressed in all three cell types (>200 TPM). Perturbation of this region led to decreased *CLIC1* mRNA expression in all CRISPRi experiments (**Figure 6H**). This region also overlaps an ATAC-seq peak in iPSCs, K562 cells, and Ngn2-induced iPSC-derived neurons and displays conservation across 241 mammals^58^ (**Figure 6H**). Validation experiments with individual gRNAs targeting this region replicated the expression changes observed in the single-cell screens across all cell types (**Figure 6I**). Perturbing the *CLIC1* CRE with dCas9^KRAB^ resulted in a >90% decrease in *CLIC1* mRNA expression, demonstrating the potency of this intronic CRE.

**Figure 6.**
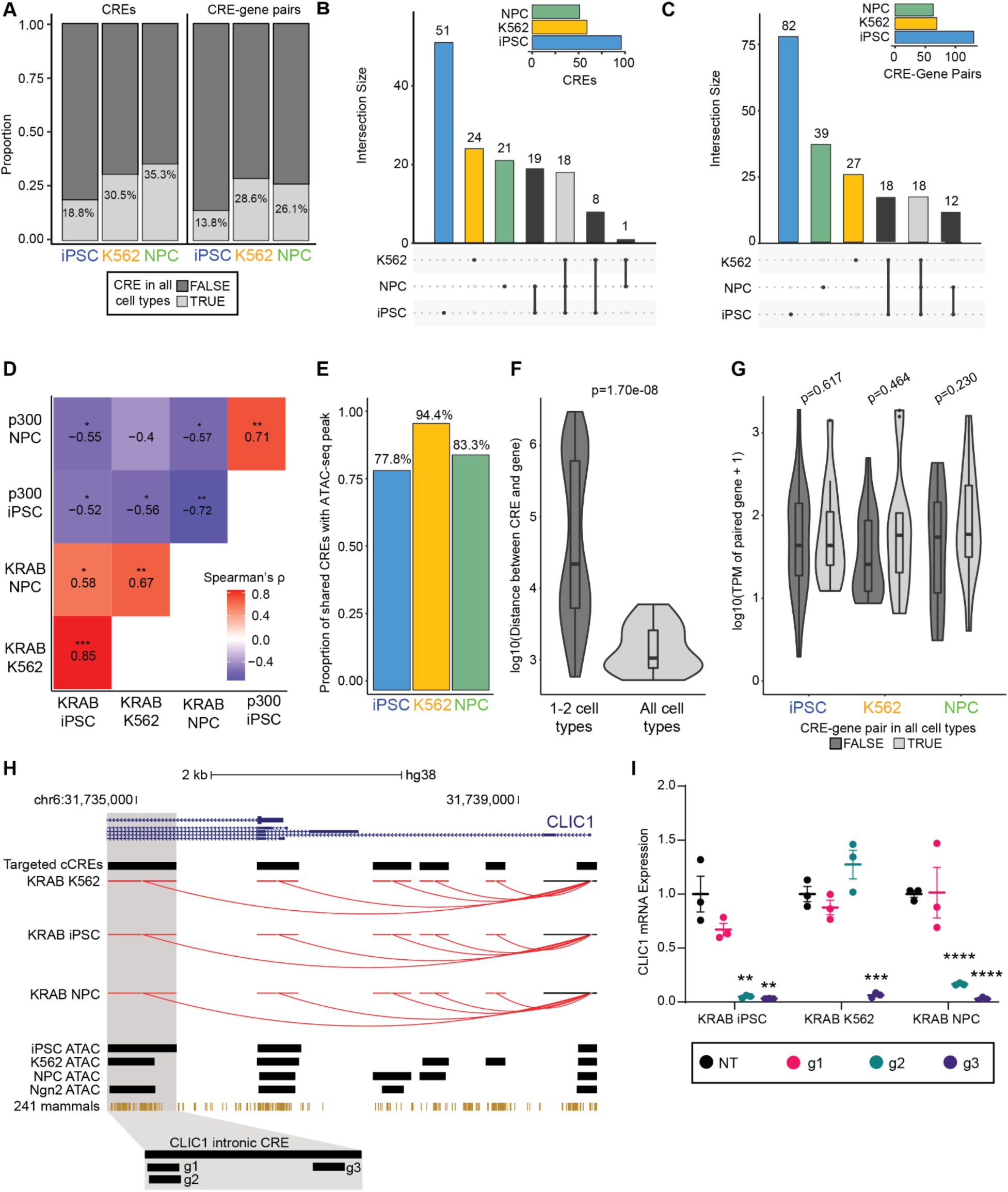
Shared CREs are marked by accessible chromatin and are located closer to target genes. **(A)** Proportion of CREs (left) and CRE-gene pairs (right) that were identified in all three cell types (light grey, ‘TRUE’) or in only one or two cell types (dark grey, ‘FALSE’). **(B)** Upset plot of CREs identified in the three cell types. Total number of CREs shown in the upper right plot. The intersection size of each set is shown in the main plot. Light grey fill indicates CREs identified in all three cell types. **(C)** Upset plot of CRE-gene pairs identified in the three cell types. Total number of CRE-gene pairs is shown in the upper right plot. The intersection size of each set is shown in the main plot. Light grey fill indicates CRE-gene pairs identified in all three cell types. **(D)** Heatmap of the Spearman’s correlation of effect sizes (Methods) for the shared CRE-gene pairs (N=17, light grey fill in **C**) for each pairwise comparison across the five experiments. Spearman correlation value is noted in each cell with the significance denoted as: *p<0.05, **p<0.01, ***p<0.001. **(E)** The proportion of shared CREs (N=20, light grey fill in **B**) that overlap ATAC-seq peaks in each cell type. **(F)** Violin and boxplots comparing the distance between CREs and connected genes for CRE-gene pairs only detected in one or two cell types (dark grey, ‘1-2 cell types’) or detected in all three cell types (light grey, ‘All cell types’). P-value from Wilcoxon test is noted in the plot. **(G)** Violin and boxplots comparing the basal expression level of genes connected to a CRE in only one or two cell types (dark grey, ‘FALSE’) or in all three cell types (light grey, ‘TRUE’) (Methods). The p-value from the Wilcoxon test for each cell type is noted in the plot. **(H)** Example of a CRE-gene pair identified in all three cell types and in all five experiments. The gene annotation and gRNA library target regions are shown, followed by the CRE-gene connections in the KRAB K562, KRAB iPSC, KRAB NPC, p300 iPSC, and p300 NPC screens, with red- and green-colored arcs indicated a decrease or increase in *CLIC1* mRNA expression, respectively. ATAC-seq peaks for all cell types included in the gRNA library design, conserved bases across 241 mammals^3^, and the gRNA locations used in individual gRNA validation experiments are shown below. **(I)** Change in *CLIC1* mRNA expression measured via RT-qPCR following delivery of individual gRNAs targeting the CRE in **(H)** to all KRAB-expressing cell lines (top) and p300-expressing cell lines (bottom). Bars represent mean +/- 1 SEM (N=3). ‘NT’ denotes nontargeting gRNA. Significant comparisons from t-test (p<0.05) comparing NT and an individual targeting gRNA is denoted as follows: *p<0.05, **p<0.01, ***p<0.001, ****p<0.0001.

### CRE-gene connections are largely cell type-specific

Given the distinct lineages of iPSCs, NPCs, and K562 cells, we tested whether the CRE-gene links are specific to individual cell types. Of the 142 total CREs, 68% (96/142) were significant in only one cell type (**Figure 7A**), with cell type-specific CREs comprising 36%-53% of all CREs within each individual cell type (**Figure 7B**). CRE-gene connections showed greater cell type-specificity, with 76% (148/196) of all connections significant in only one cell type (**Figure 7C**), and 42%-63% of CRE-gene connections in a given cell type identified exclusively in that cell type (**Figure 7D**). A representative example of cell type specificity was observed in an intergenic region 25 kb upstream from *IER3*, where seven CREs (6 with CRISPRi and 1 with CRISPRa) regulated *IER3* mRNA expression exclusively in iPSCs, despite *IER3* being expressed in all three cell types (41.7 TPM iPSC vs 7.4 TPM K562 and 29.9 TPM NPC) (**Figure 7E**). Validation experiments in iPSCs confirmed that perturbing two of these CREs with dCas9^KRAB^ led to a 50%-90% decrease in *IER3* expression, akin to direct promoter perturbation (**Figure 7F**).

**Figure 7.**
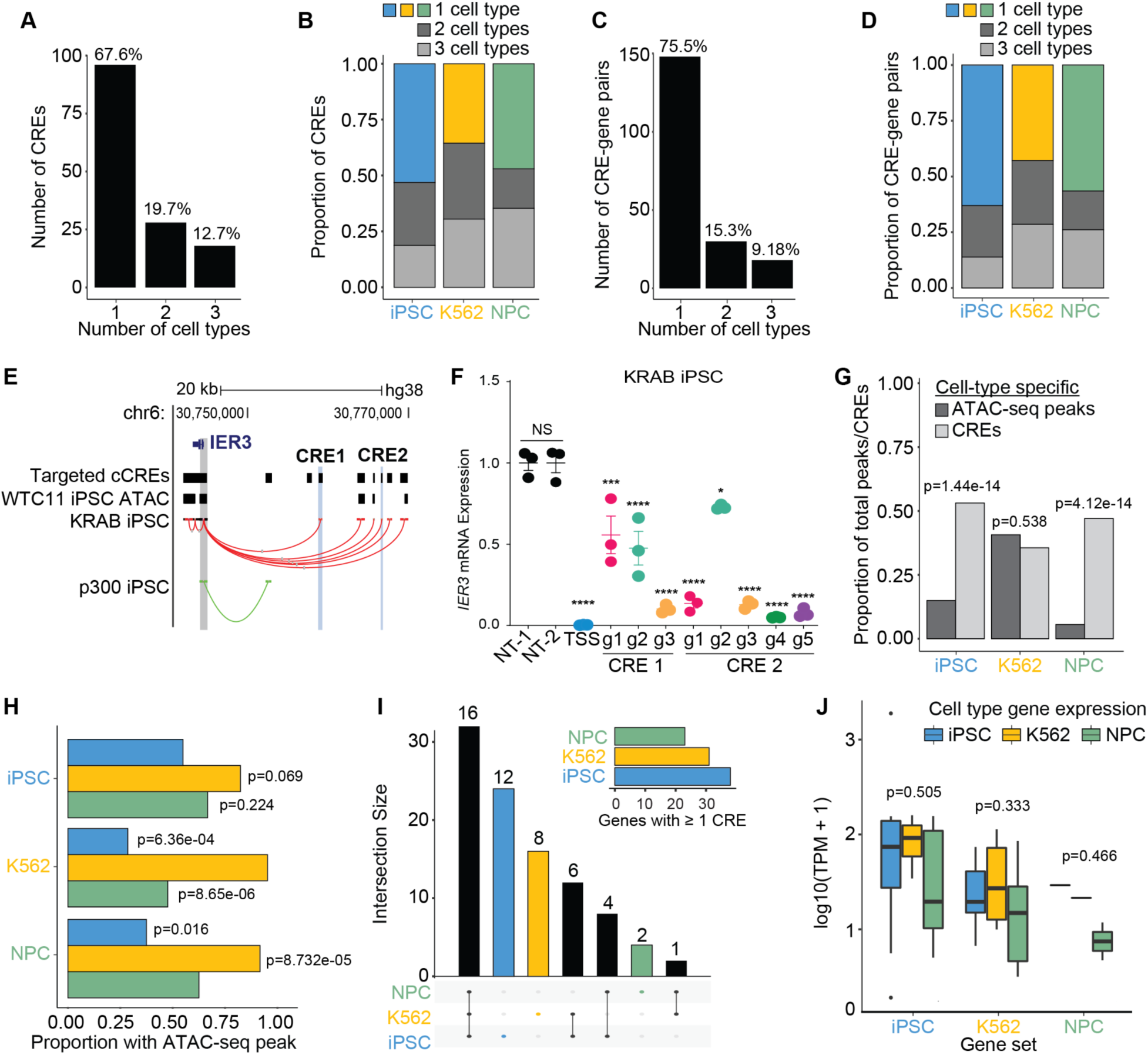
CRE-gene connections are largely cell-type specific. **(A)** Proportion of CREs in each cell type that were only detected in that cell type (cell type-specific). **(B)** Number of CRE-gene pairs identified in 1, 2, or 3 cell types. Percentage indicates the proportion of the union set of CRE-gene pairs. **(C)** Proportion of CRE-gene pairs in each cell type that were only detected in that cell type. **(D)** Number of CREs identified in 1, 2, or 3 cell types. Percentage indicates the proportion of the union set of CREs. **(E)** Example cell-type specific CRE-gene connections between intergenic CREs and *IER3* only identified in the iPSCs. The gene annotation, targeted cCREs, ATAC-seq peaks in WTC11 iPSCs are shown, followed by the CRE-gene connections in the KRAB iPSC and p300 iPSC screens, with red- and green-colored arcs indicating a decrease or increase in *IER3* mRNA expression. The light grey shading indicates a region near the TSS of *IER3* and the two blue shaded regions indicate the CREs tested in validation experiments in **(F)**. **(F)** Change in *IER3* mRNA expression measured via RT-qPCR following delivery of individual gRNAs targeting the CREs in **(E)** to KRAB iPSCs. Bars represent mean +/- 1 SEM (N=3). ‘NT’ denotes nontargeting gRNA and ‘TSS’ denotes gRNA targeting TSS of *IER3*. Results from Tukey’s post hoc tests between ‘NT-1’ and each targeting gRNA are denoted as follows: *p<0.05, **p<0.01, ***p<0.001, ****p<0.0001. **(G)** Proportion of cell type-specific ATAC-seq peaks used in library design (dark grey) and proportion of CREs identified only in that cell type (light grey). P-value from chi-squared test noted above each pair of bars. **(H)** Proportion of cell type-specific CREs that overlap ATAC-seq peaks in each cell type. P-value from chi-squared test comparing proportion of the cell-type specific CREs that overlap an ATAC-seq peak in the same cell line versus the cell line corresponding to the fill color. **(I)** Upset plot of the genes connected to at least one CRE. The total number of genes connected to a CRE is shown in the upper right and the intersection size of each set is shown in the main plot. **(J)** Boxplot comparing the expression of genes connected to a CRE in only one cell type across all three cell types. P-values from One-way ANOVA tests are noted above each group of boxplots.

Correlations between the effect sizes for all cCRE-gene tests (**Supplementary Figure 19A**) and between cell type-specific CRE-gene pairs (**Supplementary Figure 19B**) across the five screens were lower than those for shared CRE-gene pairs (**Figure 6D**). However unperturbed basal gene expression levels in the MHC locus in single cell RNA-seq (**Supplementary Figure 20A**) and in bulk RNA-seq data (**Supplementary Figure 20B**) were highly correlated across all three cell types, which supports that shared global expression patterns in the MHC locus do not imply shared cis-regulatory mechanisms. This highlights the need for functional perturbation experiments in diverse cell types to accurately map gene regulatory logic.

Since the cCREs targeted by the gRNA library constituted a union set of accessible chromatin regions across cell types, we compared the number of cell type-specific CREs to the number of accessible chromatin regions in the library design. We observed a significantly higher proportion of CREs to ATAC-seq peaks identified in iPSCs and in NPCs, relative to K562 cells (**Figure 7G**). In addition, NPC-specific CREs overlapped more with K562 ATAC-seq peaks than with NPC ATAC-seq peaks, with iPSCs trending towards a similar result (**Figure 7H**). This pattern suggests that chromatin accessibility alone may not be a sufficient predictor of cell type-specific gene regulatory activity.

Examining genes connected to CREs across cell types, iPSCs demonstrated the highest cell type-specificity (46%), followed by K562 cells (35%), and NPCs (9%) (**Figure 7I**). To rule out potential biases from basal gene expression differences and thus detection limitations, we compared the expression levels of genes connected to a CRE in only one cell type across all three cell types. There was no significant difference in basal gene expression, indicating that the presence or absence of a CRE-gene connection was not due to technical limitations of single-cell transcriptome profiling (**Figure 7J**).

We next performed gene set enrichment analysis to explore the functional pathways associated with genes linked to CREs (**Supplementary Table 12**). We focused on genes that were expressed and whose CREs were accessible in at least one other cell type to identify CRE-gene connections that were specific to cell types, rather than simply connections for genes that were only expressed in one cell type. While the MHC region is strongly associated with immune system-related pathways, genes connected to CREs were enriched in diverse pathways across the cell types, including responses to multiple stressors (e.g., heat and virus), as well as metabolic, reproductive, and developmental processes (**Supplementary Figure 21-22**). Collectively, these results highlight the cell-type specific nature of *cis*-regulatory interactions and the varied biological processes they regulate between different cell types within the same genomic region.

### Cell Type-Specific Disease Associations Revealed through Functional Perturbations in the MHC Region

Since GWAS have implicated the MHC region in >100 disorders, we investigated whether functional perturbations could reveal cell type-specific disease associations. We calculated the enrichment of genome-wide significant SNP associations (for 14 diverse traits) located in CREs. We evaluated different significance thresholds for both SNPs and CREs relative to a permutation-derived null distribution that maintained the original LD structure (**Methods**, **Supplementary Table 13**). Among the 14 traits analyzed, we found significant enrichment or depletion (multiple comparison corrected P-value < 0.05) for GWAS SNPs in CREs for schizophrenia, bipolar disorder, educational attainment, neuroticism, complete blood count (CBC), hematocrit, CBC neutrophil, CKD, and type 2 diabetes (**Figure 8, Supplementary Figure 23**). Notably, the enrichment of GWAS SNP associations became more significant as the CRE-gene connection and GWAS SNP for schizophrenia gained significance; this pattern was not observed for the 13 other traits. Additionally, the NPC CREs showed specific enrichment for SCZ SNPs as compared to SNPs from multiple traits being enriched in iPSC and K562 CREs. These results are consistent with the strong association of the MHC with schizophrenia^59,60^ and with the neurological basis of schizophrenia, suggesting that CRE-gene links in NPCs can be leveraged to narrow the search space for functional non-coding variants associated with psychiatric disorders. These detected enrichments, despite extensive LD, indicate that CRE-gene connections are enriched for disease-associated variants and highlight the importance of conducting such functional perturbation experiments across diverse, biologically relevant cell types to better understand disease mechanisms.

**Figure 8.**
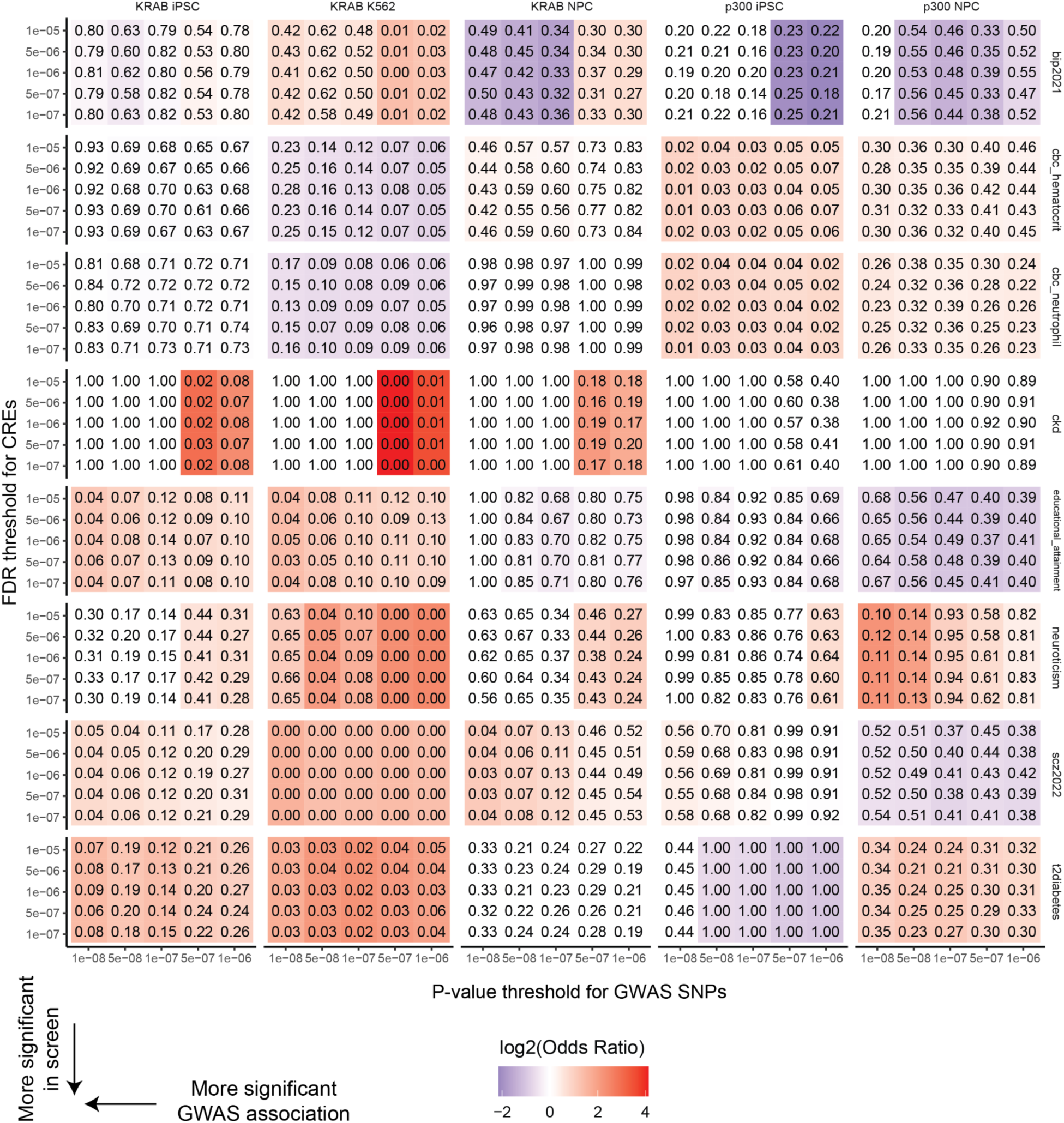
Disease relevance of regulatory-element gene connections. Heatmap of overlap of CREs with GWAS SNPs with shading indicating log2(Odds ratio) and p-values from Chi-squared test noted in each cell (**Methods**) for traits with significant enrichment in at least one cell type (p<0.05; Bipolar Disorder (Bip2021), Complete Blood Count (CBC) Hematocrit, CBC Neutrophil, chronic kidney disease (CKD), educational attainment, neuroticism, Schizophrenia (scz2022), and Type 2 Diabetes (t2diabetes) respectively). The x-axis indicates the significance threshold for GWAS SNPs and the y-axis indicates the significance threshold for the CREs.

## Discussion

Our study represents a significant advance in understanding the complex regulatory dynamics governing the MHC locus. Through the application of CRISPRi and CRISPRa perturbations to 581 cCREs in three distinct human cell types, we present the first comprehensive resource of functionally validated regulatory interactions in the MHC locus. This study marks one of the largest sets of noncoding CRISPR screens to date using single-cell RNA-seq readouts (12,625 perturbations in ∼1.2 million cells). In three cell types, we found that most distal CRE-gene connections are cell-type specific. The combination of CRISPRi and CRISPRa allowed identification of CREs and evaluation of a far broader range of gene expression levels than possible with either method alone. Our findings illuminate the complex landscape of CRE regulation, in which a single CRE can regulate multiple genes, some genes are regulated by several CREs, and CRE-gene interactions extend across short and long genomic distances. Moreover, we identified CREs in diverse genomic regions (e.g., intronic, intergenic, and 3’ UTRs), and revealed the versatility of promoters functioning as CREs for multiple genes.

By characterizing the epigenomic features defining CREs across the three cell types, we observed an enrichment of activating histone marks, consistent with previous studies associating these marks with active enhancer signatures^26,43,61^. In addition, our analysis revealed unique combinations of genomic and epigenetic features specific to each cell type, as in previous studies^21^. We also identified poised CREs in all cell types, marked by the combination of accessible chromatin and repressive histone modifications, as well as CREs overlapping active histone marks and REST ChIP-seq peaks in K562 cells and in NPCs. These findings support the crucial role of genes in the MHC locus in regulating pluripotency and differentiation^49,62,63^ and inform their mechanism of regulation.

Contrary to findings from previous noncoding CRISPR screens^43^, our results challenge the conventional notion that chromatin accessibility is a defining feature for all functional CREs. Specifically, we showed that perturbations of CREs located outside of open chromatin regions can also significantly alter gene expression. Furthermore, we demonstrated that quiescent chromatin regions could exhibit regulatory activity, suggesting that the large portions of the MHC locus predicted to have minimal regulatory potential in a particular cell type can still elicit significant changes in gene expression when perturbed. Importantly, our results also challenge the assumption that the absence of active histone marks, TF binding, or chromatin accessibility, in a given genomic region implies a lack of potential *cis*-regulatory function. This reveals a far more complex regulatory landscape within the MHC locus than previously known, extending beyond the simpler definitions of distal enhancer-promoter interactions. Future work is required to determine the relationship of these CRE-gene connections identified via synthetic perturbation, which reveals the potential for activity, to a native physiologic function in specific experimental systems or biological contexts.

Interestingly, we found that over 40% of CREs in iPSCs and NPCs impact a more distal target gene than the nearest gene, and few CRE-gene pairs showed chromatin interactions detectable by HiCAR across all cell types. When assessing the performance of multiple computational methods for predicting enhancer-gene interactions, which incorporate genomic and epigenomic data, we found that only 52%-94% of CRE-gene pairs were accurately predicted in any given cell type. This highlights the importance of functional perturbation experiments to reliably map CRE-gene interactions, underscores the complexity of gene regulation, and cautions against sole reliance on current computational predictions. Additionally, the E2G model performed best for K562 cells. Since the model is trained using K562 enhancer-gene links, the lower performance in iPSCs and NPCs demonstrates the need for cell-type specific training data. It is important to note, however, that these prediction methods may be limited by the thresholds set by the models or peak- and loop-calling methods, or the unique challenges posed by features such as gene density and regulatory complexity of the MHC locus.

We identified a number of CRE-gene connections that regulated genes in the opposite direction than expected, indicating the potential presence of silencers. These silencers may work through different mechanisms, including: directly repressing target genes, indirectly silencing through enhancer perturbation of nearby genes that inhibit the expression of the target gene, or affecting boundary elements (e.g. CTCF binding, TAD boundaries), leading to abnormal regulation by nearby enhancers in the opposite direction. Future work will be needed to delineate these or other possible mechanisms.

By examining the same set of cCREs across three distinct cell types, we identified shared CREs with regulatory activity across all three cell types and regulating the same gene. Notably, shared CREs were often located closer to transcription start sites and may be interfering with nearby promoters via spread of epigenetic marks created by dCas9^KRAB^ and dCas9^p300^. The vast majority of CREs we identified were cell type-specific, with 68% of CREs and over 76% of CRE-gene pairs uniquely identified in a single-cell type. Interestingly, iPSCs displayed the highest degree of cell type specificity and the greatest number of CRE-gene connections. This likely reflects the dynamic chromatin landscape inherent in pluripotent cell types, which progressively becomes more constrained during differentiation and lineage specification^64,65^. While the MHC locus is conventionally associated with immune response, our findings revealed that CREs also regulate genes linked to various non-immune biological processes in a cell type-specific manner.

Finally, we leveraged the identified CRE-gene links to inform the cell-type relevance for GWAS traits. A consistent trend emerged, with increasing significance at more stringent thresholds, reinforcing the hypothesis that the co-localization of GWAS-significant SNPs with CREs identified here has biological relevance. Specifically, we observed a notable enrichment of SCZ-associated SNPs within significant CREs, suggesting a potential mechanistic link between the *cis*-regulatory landscape of the MHC locus and the genetic underpinnings of SCZ. Strengthening the association of the MHC with SCZ, we identified CREs for several synaptic-and neurodevelopment-related genes (*TUBB*, *VARS2*, *CSNK2B*, *ABHD16A*) and subsequently validated a CRE that regulates *HLA-B*, which has been associated with side effects of antipsychotic medication (e.g., clozapine-induced agranulocytosis/granulocytopenia^66^). Additionally, we showed the expression of genes that are important targets for engineering hypoimmunogenic iPSCs^67^ and for cell transplantation^68^ (*HLA-A*, *HLA-B*, *HLA-C*, *HLA-DMA*) is regulated via multiple CREs. These results emphasize the potential of functional perturbation experiments to elucidate CREs relevant to specific diseases or traits.

Given that the MHC locus is the most complex region in the human genome, identifying variants that alter gene expression and ultimately lead to downstream phenotypes is challenging. This study takes a crucial step in nominating functional variants by mapping the cis-regulatory landscape, identifying CREs, and delineating their target genes. Collectively, this work provides a resource for expanding the definitions of CREs, improving computational methods for predicting CRE-gene interactions, and ultimately identifying variant-gene-phenotype relationships across the more than 100 disorders associated with the MHC locus.

## Notes

https://data.igvf.org

### Competing Interest Statement

C.A.G. is a co-founder of Tune Therapeutics, Sollus Therapeutics, and Locus Biosciences and is an advisor to Tune Therapeutics, Sollus Therapeutics, Pappas Capital, and Sarepta Therapeutics. T.S.K. is a co-founder of Tune Therapeutics. M.t.W., T.S.K., S.S.A., N.I., T.E.R., G.E.C., and C.A.G. are inventors on patents or patent applications related to CRISPR epigenome editing and screening technologies. PFS was a consultant and shareholder for Neumora Therapeutics. L.R.B. is an employee of Xaira Therapeutics (all work performed prior).

### Funding

- National Institutes of Health: HG011123, MH125236, HG012053, HG011967, HG012003
- U.S. National Science Foundation: EFMA-1830957, NSF-GRFP DGE - 2139754
- Open Philanthropy Project

