## Supplementary Notes and Methods for "Functional Annotation of the Major Histocompatibility Complex Locus"

**Table of Contents**


**Table of Contents** 1
**Supplementary Notes** 3
Supplementary Note 1. Rationale for selecting the MHC locus. 3
Supplementary Note 2. Cell line selection. 5
**Supplementary Tables** 6
**Methods** 7
Individual gRNA cloning 7
Lentivirus production 8
Cell line generation and maintenance 8
iPSC culture 8
WTC11 iPSC dCas9-effector cell lines 8
NPC culture 9
K562 culture 10
HEK293T culture 11
Bulk RNA-seq 11
Bulk RNA-seq analysis 11
Bulk ATAC-seq 11
Bulk ATAC-seq analysis 11
gRNA library design 12
gRNA library cloning 12
Lentivirus titering 14
iPSCs 14
NPCs 14
K562s 14
Single cell CRISPR screens 15
iPSCs 15
NPCs 15
K562s 15
scRNA-seq library prep 15
gRNA library PCR 16
Targeted enrichment 16
Probe design 16
Hybridization of single cell gene expression libraries 17
Sequencing 17
Individual gRNA validations 17

iPSCs 17
NPCs 17
K562s 18
HiCAR validations 18
Single cell screen analysis pipeline 19
Data processing 19
gRNA assignment 19
Differential expression testing 19
EP-length calculation 20
Data visualization 20
Evaluation of targeted enrichment panel 2
0
Quantile-quantile plots 21
Comparison of effect sizes by type of gRNA 21
Single cell screen effect size versus individual gRNA validations in KRAB iPSCs 21
Correlation of CRISPRi and CRISPRa screens 21
Correlation between change in gene expression and CRE-gene pair distance. 21
Feature enrichment analysis 21
Genomic and epigenomic annotations 21
chromHMM annotations 22
ENCODE SCREEN cCREs 22
Comparison to other prediction methods 22
Nearest gene 22
ABC model 22
E2G model 23
EnhancerAtlas 23
HiCAR 23
Definition of ‘shared’ and ‘cell-type’ specific CREs and CRE-gene connections 23
Correlation of effect sizes for CRE-gene links 24
Correlation of basal gene expression 24
Comparison of effect sizes for all cCRE-gene tests 24
Comparison of proportion of CREs and library that overlap ATAC-seq peaks 24
Gene ontology enrichment analysis 24
Enrichment of GWAS SNPs in CREs 25
HiCAR analysis 25
**Data availability** **25**
**Code Availability** **25**
**Materials availability** **25**
**Author Contributions** **26**
**Funding** **26**
**Acknowledgements** **26**

**Conflicts of Interest** **26**
**References** **27**

### Supplementary Notes

#### Supplementary Note 1. Rationale for selecting the MHC locus.

In our experiments, we targeted cCREs in a 3.5Mb region (chr6:29.700-33.200Mb, hg38) central to the human major histocompatibility complex (MHC). We selected the MHC due to its enormous importance to human health and disease. However, its high gene density, sequence diversity, and linkage disequilibrium (LD) greatly complicate fine-mapping efforts.

Genomic features. We compared the genomic features of the 3.5Mb MHC region to 655 autosomal regions of the same size (**Supplementary Figure 1.1**). These regions were consecutive genomic segments of the same size. Each bin was annotated with the numbers of: GC bases; constrained bases across 240 mammals (phyloP ≥ 2.270) and in 43 primates (phastCons ≥ 0.961)[1], unique protein-coding sequence (CDS) base pairs, transcription start sites (TSS), 5'UTR base pairs, 3'UTR base pairs, and intronic base pairs from GENCODE gene/transcript models[2], regulatory features from ENCODE3[3] including candidate cis-regulatory elements of unique promoter-like sequence (PLS) and enhancer-like sequence (ELS) base pairs along with DNase hypersensitive sites (DHS, 243 cell lines / tissues), and transcription factor binding site footprints (TFBS). For human genetic variation, biallelic SNPs are from TOPMed[4] (v8, N=140,306), contrasting common SNPs (allele frequency, AF ≥ 0.005, 2.6% of all SNPs) and ultra-rare SNPs (allele count, AC == 1, 46.2% of all SNPs) and including LD scores in the European ancestry subsample. Unique eQTL SNPs (any tissue) are from GTEx v8[5]. Unique genome-wide significant SNPs are from the NHGRI/EBI GWAS Catalog (10/2021)[6]. The schizophrenia results are from the latest analysis from the Psychiatric Genomics Consortium (PGC scz2022)[7]. MHC rank scoring 1=most, rank 656=least, and percentile scoring: 100=most, 0=least.

The rank and percentile of the MHC in genomic context is provided in **Supplementary Table 1**. The MHC has relatively high GC content and evolutionary constraint. It has the greatest frequency of coding bases and is at the 99$^{th}$ percentile for TSS and 5'UTRs. The MHC is dense with regulatory features, particularly promoter-like sequence and TF binding sites. Most notably, the MHC region has exceptionally high LD in terms of LD scores, long-range LD[8] , and LD blocks[9]. In addition, the MHC ranks first for GWAS findings, eQTL SNPs, and common SNPs.

Gene content. The MHC contains 300 annotated genes. These include: 138 protein-coding genes, 78 pseudogene biotypes, 50 long intergenic non-coding RNAs (lncRNA), 9 microRNAs (miRNA), and 25 non-coding genes of various types (e.g., snoRNA or snRNA). Most notable among the protein-coding genes are the MHC Class 1 cell surface receptors (all classical and many of the non-classical) that are expressed in all nucleated cells as well as in platelets along with the antigen-presenting MHC Class 2 cell surface proteins. This region contains many other notable genes including *POU5F1* (Oct-4, a pluripotency marker and one of the Yamanaka reprogramming factors).

Given these functions, 33 MHC genes have an OMIM annotation[10], and these are dominated by diseases characterized by dysfunction of the immune system (e.g., malaria susceptibility and autoimmune conditions like multiple sclerosis and rheumatoid arthritis). There are an additional

33 MHC genes expressed at high and relatively constant levels across many tissues (i.e., “housekeeping” genes)[11].

Gene set analysis. We conducted gene set analyses of protein-coding genes in the MHC against a background of 19,022 autosomal protein-coding genes (hypergeometric tests, Bonferroni-corrected P-value to < 0.005). 82 gene sets were significant which clustered into three groups: positive regulation of immune response, adaptive immune response, and antigen processing and presentation.

Human genomic findings. There are 33 MHC genes with an OMIM entry (**Supplementary Table 13**). However, common genetic variation detectable using GWAS is notably enhanced in the MHC region. This 3.5 Mb region is about 0.1% of the genome but accounts for 18.2% of all unique SNP-trait associations (=503/2763). The subset of the 503 traits with ≥ 10 unique SNP-trait associations in the MHC region is listed in **Supplementary Table 3**. The numbers of GWAS associations, OMIM entries, and eQTL-SNPs by chromosome, are shown in **Supplementary Figure 1**. Chromosome 6 is a major outlier for GWAS associations, a notable outlier for eQTL SNPs, but not for OMIM entries. Chromosome 6 is relatively large (170.8 Mb) and the MHC locus comprises a small fraction (3.5 Mb). The gene density of chromosome 6, GWAS associations, and eQTL-SNPs per 100 kb, are shown in **Supplementary Figure 2**. Most of chr6 is relatively homogeneous for these features with the MHC region (blue bar) being markedly discrepant. LD scores on chromosome 6 for four ancestry groups are shown in **Supplementary Figure 3**. All global groups show sharply greater LD scores in the MHC region.

**Supplementary Note 2. Cell line selection.**

To assay cis-regulatory mechanisms across a diverse set of cell types, we selected the following cell lines to include in the assays: i) the human leukemia cell line, K562; ii) human induced pluripotent stem cells (iPSCs); and iii) iPSC-derived neural progenitor cells (NPCs). The K562 cell line represents a non-neuronal control cell type, has been extensively characterized by the Roadmap and ENCODE Consortiums, and has successfully been used for CRISPR screens by our group and others. We elected to perform only CRISPRi perturbations in K562 cells because this cell type contributed the greatest number of target cCREs to the library design. The WTC11 iPSCs are an additional non-neuronal cell line that can be differentiated towards a neural lineage and are important for characterizing regulatory mechanisms within the MHC region which contains the pluripotency-related gene, *POU5F1*. NPCs represent a neural lineage, are amenable to large scale experiments, and the cell populations are more homogenous than further differentiated neuronal cultures.

**Supplementary Tables**

SupplementaryTables
Supplementary Table 1. Characteristics of the MHC relative to other genomic regions.
Supplementary Table 2. Disease relevance of genes in the MHC locus.
Supplementary Table 3. Traits associated with the MHC locus using GWA studies.
Supplementary Table 4. CRE-gene pair differential gene expression results.
Supplementary Table 5. CRE-gene pairs selected for individual gRNA validation.
Supplementary Table 6. Individual gRNA validation results.
Supplementary Table 7. Fisher's exact test results for genomic and epigenomic features for all CREs.
Supplementary Table 8. Fisher's exact test results for chromHMM annotations for all CREs.
Supplementary Table 9. Fisher's exact test results for genomic and epigenomic features for repressor CREs.
Supplementary Table 10. Fisher's exact test results for chromHMM annotations for repressor CREs.
Supplementary Table 11. Benchmarking prediction methods with validated CRE-gene pairs.
Supplementary Table 12. Gene ontology analysis results.
Supplementary Table 13. GWAS SNP enrichment test results.
Supplementary Table 14. gRNA and PCR primer sequences.
Supplementary Table 15. Taqman probes used in RT-qPCR.
Supplementary Table 16. Publicly available and new datasets used in this study.

### Methods

#### Individual gRNA cloning

All gRNAs for individual gRNA experiments were cloned into a U6-driven gRNA vector with Blasticidin-resistance and dsRed fluorophore (Addgene #83919). The gRNA vector was digested with Esp3I (NEB #R0734L) for one hour at 37C then gel purified using the Zymoclean Gel DNA Recovery Kit (Zymo Research #D4007). Protospacer sequences are listed in **Supplementary Table 14**. Additional sequence for cloning was appended as previously described[12] and oligos were synthesized by IDT. The oligos were resuspended with water to 100uM. For each gRNA, 2 uL of the sense and antisense oligos were combined with 16 uL of DPBS (ThermoFisher #14190144). Next, the oligos were phosphorylated and annealed a follows:

| Reagent | Volume |
|---|---|
| Mixed oligos (1000 ng) | 5 uL |
| T4 DNA Ligase Buffer (NEB #B0202S) | 2 uL |
| PNK (NEB #M0201S) | 1 uL |
| Water | 12 uL |

This reaction was incubated at 37C for 30 minutes, at 65C for 20 minutes, and then ramped down at 5C per minute from 95C to 25C. Next, the phosphorylated and annealed oligos were diluted 1:50 with water and then ligated to the digested gRNA backbone as follows:

| Reagent | Volume |
|---|---|
| 1:50 dilution of insert | 1.5 uL |
| T4 DNA Ligase Buffer (NEB #B0202S) | 1 uL |
| T4 DNA Ligase (NEB #M0202L) | 1 uL |
| Water | To 10 uL total volume |

This reaction was incubated for 16 hours at 16C. Then 4 uL was transformed into 50uL of STBL3 cells. Individual colonies were grown overnight in 5 mL of LB broth and plasmids were purified using the QIAprep Spin Miniprep Kit (Qiagen #27106). The sequence of each gRNA was confirmed via Sanger sequencing (Genewiz).

#### Lentivirus production

For all lentivirus production, HEK293Ts of passage numbers 5-15 were used. For dCas9-effector constructs and gRNA libraries, HEK293T cells were seeded at 4e6 cells per 10cm dish. For individual gRNA validations, HEK293T cells were seeded at 800K cells per well of a 6-well tissue culture plate. 24-hours later, HEK293T cells were transfected using Lipofectamine 3000 with psPAX2 (Addgene #12260), pMD2.G (Addgene #12259), and the dCas9-effector plasmid (Addgene #83890, #83889). Media was changed 16 hours post-transfection and lentivirus was harvested 24 hours after the media change. The lentivirus was filtered using 0.45uM filters (MilliporeSigma #SLHVR33RS) and then concentrated with Lenti-X according to the manufacturer's protocol. For dCas9-effector constructs and individual gRNA validations, the lentivirus was resuspended in DPBS to a 20X final concentration. For gRNA libraries, the lentivirus was resuspended to a 50X final concentration. The lentivirus was then aliquoted into single-use tubes and stored at -80C.

**Cell line generation and maintenance**

All cell lines were cultured at 37C and 5% CO2.

<u>iPSC culture</u>

WTC11 iPSCs were obtained from the Coriell Institute for Biomedical Research (GM25256) and were propagated in mTeSR1 (StemCell Technologies #85850) on Matrigel (Corning #354230)-coated tissue culture plates and passaged as colonies using Versene (ThermoFisher #15040066). Cells were frozen in freezing solution comprised of 60% mTeSR1 media, 10% DMSO, and 30% KnockOut™ Serum Replacement (ThermoFisher #10828028), with 10uM Y-27632 (Dihydrochloride) (StemCell Technologies (#72302).

<u>WTC11 iPSC dCas9-effector cell lines</u>

*Monoclonal cell line generation*

To generate the dCas9-effector WTC11 iPSC lines, WTC11 iPSCs were passaged with Accutase (VWR #490007-741) and seeded at 200k cells per well of a 6-well Matrigel-coated plate with 10uM Y-27632. 24 hours after seeding, cells were transduced with 100uL of either the dCas9-KRAB or dCas9-p300 lentivirus. 24 hours post-transduction, a media change was performed. 24 hours after the initial media change, cells were passaged using Accutase and seeded at 125k cells per well of a 6-well Matrigel-coated plate with 10uM Y-27632. The dCas9-KRAB and dCas9-p300 bulk transduced populations were then treated with 50ug/mL hygromycin B (ThermoFisher #10687010) or 1 ug/mL puromycin (ThermoFisher #J67236.XF) for two weeks. The monoclonal cell lines were generated following the CloneR system according to the manufacturer's protocol (StemCell Technologies #05888). Seven days after plating in the 10cm-dish format, individual colonies were manually picked using a P200 pipette and plated into individual wells of a 96-well plate coated with Matrigel. Media was changed daily until the wells became confluent. Once confluent, monoclonal cell lines were expanded into 48-well plates and then 24-well plates.

*Characterization*

gDNA (genomic DNA) was extracted using the DNeasy Blood & Tissue Kit (Qiagen #69504) according to the manufacturer's protocol and PCRs were performed to confirm presence of dCas9-effector constructs using primer sequences in **Supplementary Table 14** as previously described[12]. PCR products were run on a 1% agarose gel for 45 minutes at 100V to confirm product size. After confirming presence of construct, each monoclonal cell line was transduced with an individual gRNA targeting the promoter the POU5F1[13] and targeting the promoter of IL1RN[14] for dCas9-KRAB and dCas9-p300, respectively, or transduced with a nontargeting gRNA (N=3 each). 24 hours post-transduction, the media was changed. Cells were then selected with 2.5ug/mL Blasticidin S HCl (ThermoFisher #A1113903) for 3-5 days. The cells were then harvested using Accutase and mRNA was purified using the Qiagen RNeasy kit according to the manufacturer's protocol. 250 ng mRNA was used as input for cDNA amplification using the Invitrogen™ SuperScript™ VILO™ cDNA Synthesis Kit (ThermoFisher #11754050). For RT-qPCR, each reaction contained 2 uL cDNA, 6 uL H20, 1 uL Taqman probe for 18S, 1 uL Taqman probe for gene of interest, and 10 uL PerfeCTa FastMix II (Quantabio #95118). Delta delta Ct analysis was performed in Microsoft Excel. Graphpad Prism was utilized to conduct one-way ANOVA tests followed by Tukey's HSD for post-hoc testing. Significance is reported in figures as follows: *p-value < 0.05, **p-value < 0.01, ***p-value < 0.001. Taqman probe information is provided in **Supplementary Table 15**. Normal karyotypes of the selected clones for each dCas9-effector were confirmed by Cell Line Genetics, Inc.

<u>NPC culture</u>

WTC11 NPCs were cultured in Neural Basal Medium (NBM), consisting of DMEM/F-12 containing 1% Penicillin-Streptomycin (10,000 units/mL) (ThermoFisher #15140122), 2% B27 Supplement w/o Vitamin A (50X) (ThermoFisher #12587010), and 1% N-2 Supplement (100X) (ThermoFisher #12587010), supplemented with 20 ug/mL Animal-Free Recombinant Human EGF (Peprotech #AF-100-15) and 20 ug/mL Animal-Free Recombinant Human FGF-basic (Peprotech #AF-100-18B) (NEM).

*NPC differentiation*

WTC11 iPSCs were differentiated into NPCs using embryoid-body (EB) formation and small molecules as previously described with minor modifications[15,16]. WTC11 iPSCs were seeded in low-adhesion 6-well culture plates (Corning #3471) in NBM containing 10 uM SB431542 (StemCell Technologies #72234) and 0.1 uM LDN193189 (StemCell Technologies #72147) (NIM) with 10 uM Y-27632. 24 hours later (Day 1), 2 mL of media was removed and replaced with 2 mL NIM. On Days 2-4, 3 mL of media was removed and replaced with 3 mL NIM. On Day 5, EBs were transferred onto Matrigel-coated 6-well plates containing 1 mL NIM per well. 24 hours later (Day 6), all media was removed and replaced with 3 mL NIM. On Days 7-12, 3 mL NIM was changed daily. On Day 13, neural rosettes were harvested using STEMdiff™ Neural Rosette Selection Reagent according to the manufacturer's protocol (StemCell Technologies #05832) and plated onto Matrigel-coated plates containing NEM ('Passage 0'). NPCs were propagated for five passages before characterization as described below. All experiments were performed with NPCs of passage number 7-15.

*Characterization*

gDNA was extracted using the DNeasy Blood & Tissue Kit (Qiagen #69504) according to the manufacturer's protocol and PCRs were performed to confirm presence of dCas9-effector constructs using primer sequences in **Supplementary Table 14** as previously described[12]. PCR products were run on a 1% agarose gel for 45 minutes at 100V to confirm product size. After confirming presence of construct, each NPC dCas9-effector cell line was transduced with an individual gRNA targeting the promoter of CHD8 and targeting the promoter of IL1RN for dCas9-KRAB and dCas9-p300, respectively, or transduced with a nontargeting gRNA (N=3 each). 24 hours post-transduction, the media was changed. Cells were then selected with 2.5ug/mL Blasticidin S HCl (ThermoFisher #A1113903) for 3-5 days. The cells were then harvested using Accutase and mRNA was purified using the Qiagen RNeasy kit according to the manufacturer's protocol. 250 ng mRNA was used as input for cDNA amplification using the Invitrogen™ SuperScript™ VILO™ cDNA Synthesis Kit (ThermoFisher #11754050). For RT-qPCR, each reaction contained 2 uL cDNA, 6 uL H20, 1 uL Taqman probe for 18S, 1 uL Taqman probe for gene of interest, and 10 uL PerfeCTa FastMix II (Quantabio #95118). Delta delta Ct analysis was performed in Microsoft Excel. Graphpad Prism was utilized to conduct one-way ANOVA tests followed by Tukey's HSD for post-hoc testing. Significance is reported in figures as follows: *p-value < 0.05, **p-value < 0.01, ***p-value < 0.001. Taqman probe information is provided in **Supplementary Table 15**.

<u>K562 culture</u>

K562 cells were obtained from the American Tissue Collection Center (ATCC) via the Duke University Cancer Center Facilities. K562 cells were cultured in RPMI1640 medium supplemented with 10% FBS and 1% penicillin-streptomycin. The polyclonal dCas9-KRAB cell line was generated as previously described[17]. K562 cells were transduced with dCas9-KRAB (Addgene #83890) lentivirus with polybrene at a concentration of 8 ug/mL. At two days post-transduction, cells were selected for 10 days with 600 ug/mL hygromycin B (ThermoFisher #10687010). Following selection, polyclonal cells were stained to detect expression of dCas9-KRAB protein.

*Characterization*

The dCas9-KRAB K562 cells were transduced with an individual gRNA targeting the HS2 enhancer region[18] or transduced with a nontargeting gRNA (N=3 each) via spinfection. 25,000 cells were resuspended in 0.5 mL of media containing 25 uL of lentivirus and 8 ug/mL polybrene, then centrifuged for 30 minutes at 25C. The volume was transferred into one well of a 24-well plate. 24 hours post-transduction, the cells were centrifuged for 5 minutes at 300g, resuspended in fresh media, and plated into a 6-well plate. Cells were then selected with 2.5ug/mL Blasticidin S HCl (ThermoFisher #A1113903) for 3 days. The cells were harvested and mRNA was purified using the Qiagen RNeasy kit according to the manufacturer's protocol. 250 ng mRNA was used as input for cDNA amplification using the Invitrogen™ SuperScript™ VILO™ cDNA Synthesis Kit (ThermoFisher #11754050). For RT-qPCR, each reaction contained 2 uL cDNA, 6 uL H20, 1 uL 10 uM FWD primer, 1 uL 10uM RVS primer, and 10 uL PerfeCTa SYBR Green FastMix (Quantbio #95072). Delta delta Ct analysis was performed in Microsoft Excel. Graphpad Prism was utilized to conduct one-way ANOVA tests followed by Tukey's HSD for post-hoc testing. Significance is reported in figures as follows: *p-value < 0.05, **p-value < 0.01, ***p-value < 0.001. Taqman probe information is provided in **Supplementary Table 15**.

HEK293T culture

HEK293T cells were obtained from the American Tissue Collection Center (ATCC) via the Duke University Cancer Center Facilities and were cultured in DMEM(1X) (Thermo #) with 10% FBS (Sigma #) and 1% Penicillin-Streptomycin and passaged 1:10 every 3-4 days.

**Bulk RNA-seq**

Total RNA was purified from WTC11 iPSCs and NPCs using the Qiagen RNeasy kit according to the manufacturer's protocol. RNA quality was verified to have RIN score > 8 for all samples using High Sensitivity RNA ScreenTape Analysis (Agilent #5067). RNA-sequencing libraries were prepared by Genewiz and sequenced on an Illumina HiSeq with 2x150bp configuration. RNA-sequencing was performed in duplicate for each cell type.

**Bulk RNA-seq analysis**

The pipeline for processing and analysis of the bulk RNA-seq data can be found here: https://github.com/Duke-GCB/GGR-cwl/blob/master/v1.0/RNA-seq_pipeline/pipeline-pe-unstranded-with-sjdb.cwl. Briefly, quality control of FASTQ files was performed using FASTQC[19]. Adapter reads were then trimmed using Trimmomatic[20] and mapped using the STAR aligner[21]. Finally, quantification is performed using RSEM[22]. For comparisons of basal gene expression levels, a mean transcripts per million (TPM) value was calculated using two biological replicates.

**Bulk ATAC-seq**

The Omni-ATAC seq protocol was used to generate ATAC-sequencing libraries[23]. 100,000 NPCs were harvested using Accutase and resuspended in 1 mL of cold ATAC-seq resuspension buffer and then the remainder of the protocol was followed. Following pre-amplification, 3 additional cycles of PCR were performed prior to final amplification and cleanup. Library concentrations and sizes were obtained using the KAPA Library Quantification Kit (Roche #7960255001) and High Sensitivity DNA ScreenTape Analysis (Agilent #5067). ATAC-seq was performed for two biological replicates. The final libraries were pooled to 3 nM and sequenced on a HiSeq 4000 with 1x50bp configuration.

**Bulk ATAC-seq analysis**

The pipeline for processing and analysis of the bulk ATAC-seq data can be found here: https://github.com/Duke-GCB/GGR-cwl/blob/master/v1.0/ATAC-seq_pipeline/pipeline-se-blacklist-removal.cwl. Briefly, quality control of FASTQ files was performed using FASTQC[19]. Adapter reads were then trimmed using Trimmomatic. Reads were then aligned to the reference genome using Bowtie[24]. Duplicate reads were removed using Picard MarkDuplicates and ENCODE hg38 blacklist reads were removed using bedtools2[25]. Peak calling was performed using MACS2[26] with narrowPeak settings. Sequencing-depth normalized ATAC bigWig files were generated using deeptools[27] bamCoverage.

**gRNA library design**

ATAC-seq peaks from K562 cells, WTC11 iPSCs, WTC11-derived NPCs, and WTC11-derived Ngn2 neurons (**Supplementary Table 16**) were intersected with a 3.5Mb region of the MHC locus using (chr6:29700000-33200000) using `bedtools intersect`. All four outputs and the screen region were then intersected using bedtools multiinter and overlapping regions were merged using `bedtools merge -d -1`. Additionally, we obtained coordinates for previously identified POU5F1 enhancers[28] and converted the coordinates from hg19 to hg38 coordinate builds using the UCSC liftOver utility. In total, there were 544 merged ATAC-seq peaks and 44 *POU5F1* enhancers for a total of 588 target regions.

We then obtained all potential SpCas9 gRNAs in the 3.5Mb region of the MHC locus from the CRISPOR database[29] and intersected the gRNAs with the 588 target regions using `bedtools intersect` requiring complete overlap of gRNAs with the target regions. Duplicated gRNA sequences were removed and the remaining gRNAs were filtered for a specificity score greater than 20, 20bp protospacer length, and no polyT ('TTTT') or polyG ('GGGGG') sequences, resulting in 537 of the 544 merged ATAC-seq peak with at least one gRNA (N=581 final target regions). We then took the top 20 gRNAs ranked by specificity per target region totaling to 11,300 gRNAs. Next, we added gRNAs targeting the promoters of 15 total genes that are highly or lowly expressed in at least one cell type (10 gRNAs/genes, 150 total gRNAs): *CLIC1*, *CSNK2B*, *DDX39B*, *POU5F1*, *HBE1*, *HBG1*, *HBG2*, *MAP2*, *NANOG*, *NESTIN*, *SOX2*, *ASCL1*, *IL1RN*, *MYOD1*, *NEUROG2*. For *ASCL1*[30], *IL1RN*[14], *MYOD1*[14], and *NEUROG2*[30], we included previously published gRNAs. For the remaining gRNAs per gene, we input each gene into the MIT CRISPick design tool with the CRISPRi and SpyoCas9 parameters and selected the ten (or remaining) highest ranked gRNAs per gene. For negative controls, we generated 1,273 nontargeting, gRNA sequences that have similar sequence composition to the targeting gRNAs. The positive and negative control gRNAs were similarly filtered for 20bp protospacer lengths and no polyT ('TTTT') or polyG ('GGGGG') sequences. The final library design contained 12,723 gRNAs (see 'gRNA library cloning' for final gRNA count after cloning).

Adapter sequences were then added to each protospacer sequence as follows:

```
ATATATCTTGTGGAAAGGACGAAACACCG[20bp gRNA]GTTTAAGAGCTATGCTGGAAACAGCATAG
```

The oligos were then synthesized by Twist Bioscience in a pooled format.

**gRNA library cloning**

CROP-seq-opti (Addgene #106280) was digested with MluI-HF (NEB #R3198S) and BsiWI-HIF (NEB #R3553S) for 1 hour at 37C and then run on a 1% agarose gel for 60 minutes at 100V. The upper band was purified using the Zymoclean Gel DNA Recovery Kit (Zymo Research #D4007). The Blasticidin resistance gene (Bsr) was amplified using Bsr-fragment-FWD and Bsr-fragment-RVS primers (**Supplementary Table 14**) from pLV-U6-gRNA-UbC-eGFP-P2A-Bsr (Addgene #83925), run on a 1% agarose gel for 30 minutes at 100V and purified using the Zymoclean Gel DNA Recovery Kit (Zymo Research #D4007). The Bsr fragment was inserted into the digested CROP-seq-opti backbone sing Gibson Assembly® Master Mix (NEB #E2611L) with a 1:3 ratio of insert to backbone. The reaction was incubated for 1 hour at 50C, purified using AMPure XP SPRI reagent in 2:1 ratio of reagent to reaction (Beckman Coulter #A63881), and transformed into STBL3 cells as described above. The final plasmid (pLRB100) sequence was confirmed via Sanger sequencing (Genewiz) and via whole plasmid sequencing (Massachusetts General Hospital DNA Core).

The pool of oligos were resuspended with water to 1 ng/uL. The oligos were then PCR-amplified in the following reaction and cycling parameters:

| Reagent | Volume |
|---|---|
| Q5 2X Master Mix (# | 12.5 uL |
| 10 uM pLRB266 | 1.25 uL |
| 10 uM pLRB267 | 1.25 uL |
| gRNA library (1 ng) | 1 uL |
| Water | 9 uL |

98C for 30 seconds, 98C for 10 seconds, 63C for 30 seconds, 72C for 15 seconds (10 total cycles), 72C for 15 seconds, then hold at 4C.

Following amplification, the PCR reactions were run on a 1% agarose gel for 45 minutes at 100V. The ~140bp band was extracted and purified using the Zymoclean Gel DNA Recovery Kit (Zymo Research #D4007). pLRB100 was digested with Esp3I as described above. A Gibson assembly reaction was then performed by combining the insert and backbone in a 1:3 ratio using Gibson Assembly® Master Mix (NEB #E2611L) and incubated for 1 hour at 50C. The reaction was then purified using AMPure XP SPRI reagent in 2:1 ratio of reagent to reaction (Beckman Coulter #A63881). The purified reaction was then transformed into 50 uL Endura electrocompetent cells (Lucigen #60242-1) according to the manufacturer's protocol. The plasmid was purified using the QIAGEN Plasmid Plus Midi Kit (Qiagen #12945). To verify the gRNA library was successfully cloned, the plasmid was PCR amplified as follows:

| Reagent | Volume |
|---|---|
| 5X Q5 Reaction Buffer (NEB #M0491L) | 5 uL |
| 10 mM dNTPs (ThermoFisher #R1122) | 0.5 uL |
| 10 uM pLRB278 | 1.25 uL |
| 10 uM pLRB279 | 1.25 uL |
| Plasmid (10 ng) | 1 uL |
| Q5 HiFi DNA Polymerase (NEB #M0491L) | 0.5 uL |
| Water | 15.5 uL |

98C for 30 seconds, 98C for 10 seconds, 63C for 30 seconds, 72C for 15 seconds (15 total cycles), 72C for 2 minutes, then hold at 4C.

Following PCR amplification, the reaction was purified using AMPure XP SPRI reagent in a 0.65:1 ratio followed by a 0.35:1 ratio of reagent to reaction. The purified product was then diluted to 4 nM and sequenced on an Illumina MiSeq with 2x50bp configuration and custom sequencing primers (pLRB280, pLRB281; **Supplementary Table 14**). Upon sequencing, we confirmed the presence of 12,625 gRNAs in the plasmid pool with at least one gRNA perturbation for every targeted region (N=581) and all positive and negative controls.

**Lentivirus titering**

The estimated MOI of the gRNA library lentivirus in each cell line was empirically determined using the lentiMPRA protocol[31]. RT-qPCR was performed with primers designed to amplify the gRNA construct, WPRE element, and a region on chromosome 15 (**Supplementary Table 14**). Each reaction contained 5.25 uL of gDNA (4 ng/uL), 2.75 uL H20, 1 uL 10 uM FWD primer, 1 uL 10uM RVS primer, and 10 uL PerfeCTa SYBR Green FastMix (Quantbio #95072). The MOI was calculated as previously described[31]. The virus amount with the greatest calculated MOI that did not lead to significant morphology changes and/or cell death in each cell line was selected and this value was scaled to determine final virus volume used in screen transductions.

<u>iPSCs</u>

dCas9-KRAB and dCas9-p300 iPSCs were seeded at 40K cells per well in Matrigel-coated 24-well plates. 24 hours after seeding, lentivirus was added to the cells (0 uL, 1 uL, 2 uL, 4 uL, 8 uL, 16 uL, 32 uL, 64 uL) with three replicates per condition. 24 hours after transduction, the media was replaced. Cells were harvested using Accutase two days post-transduction and gDNA was extracted as described above.

<u>NPCs</u>

dCas9-KRAB and dCas9-p300 NPCs were seeded at 45K cells per well in Matrigel-coated 24-well plates. 24 hours after seeding, lentivirus was added to the cells (0 uL, 1 uL, 2 uL, 4 uL, 8 uL, 16 uL, 32 uL, 64 uL) with three replicates per condition. 24 hours after transduction, the media was replaced. Cells were harvested using Accutase two days post-transduction and gDNA was extracted as described above.

<u>K562s</u>

dCas9-KRAB K562s were seeded at 25K cells per well in a 24-well plate containing media with 8 ug/mL polybrene and lentivirus was added to the cells (0 uL, 1 uL, 2 uL, 4 uL, 8 uL, 16 uL, 32 uL, 64 uL) with three replicates per condition. 24 hours after transduction, the media was replaced. Cells were harvested two days post-transduction and gDNA was extracted as described above.

### Single cell CRISPR screens

#### iPSCs

dCas9-KRAB iPSCs were seeded at 1.9e6 cells in a Matrigel-coated 10 cm dish. 24 hours after seeding, 382 uL of gRNA library lentivirus was added to the cells. 24 hours post-transduction, the media was changed. Starting 48 hours post-transduction, cells were selected with 2.5 ug/mL Blasticidin S HCl (ThermoFisher #A1113903) and passaged once. Cells were harvested nine days post-transduction with Accutase and then used as input for single cell profiling, as described below.

dCas9-p300 iPSCs were seeded at 1.27e6 cells in a Matrigel-coated 10 cm dish. 24 hours after seeding, 302 uL of gRNA library lentivirus was added to the cells. 24 hours post-transduction, the media was changed. Starting 48 hours post-transduction, cells were selected with 2.5 ug/mL Blasticidin S HCl (ThermoFisher #A1113903) and passaged once. Cells were harvested nine days post-transduction with Accutase and then used as input for single cell profiling, as described below.

#### NPCs

dCas9-KRAB and dCas9-p300 NPCs were seeded at 2.5e6 cells in a Matrigel-coated 10 cm dish. 24 hours after seeding, 356 uL of gRNA library lentivirus was added to the cells. 24 hours post-transduction, media was changed. Starting 48 hours post-transduction, cells were selected with 2.5 ug/mL Blasticidin S HCl (ThermoFisher #A1113903) and passaged once. Cells were harvested seven days post-transduction with Accutase and then used as input for single cell profiling, as described below.

### K562s

dCas9-KRAB K562s were seeded at 1.5e6 cells in a 15 cm culture dish with 30 mL media containing 8 ug/mL polybrene. 480 uL of the gRNA library lentivirus was added to the cells. 24 hours post-transduction, cells were centrifuged at 300g for 5 minutes, and transferred into a new 15 cm dish containing 10 ug/mL Blasticidin S HCl (ThermoFisher #A1113903). Selection continued for six days and the cells were passaged once. Cells were harvested seven days post-transduction and then used as input for single cell profiling, as described below.

### scRNA-seq library prep

For the CRISPRi screens we used the 10X Genomics' 3' CellPlex Assay (10X Genomics #PN-1000261) to overload each lane of the 10X Genomics' microfluidic chip following the manufacturer's protocol (CG000391, Rev B). Briefly, each population of cells was split into eight equal pools and labeled with a unique CMO prior to re-pooling and diluting to 1500 cells/uL. We then proceeded with the Chromium Next GEM Single Cell 3' v3.1 gene expression assay according to the manufacturer's protocol (CG000388, Rev A) with approximately 40K cells loaded onto each lane of the Next GEM Chip G. Library concentrations and sizes were obtained using the KAPA Library Quantification Kit (Roche #7960255001) and High Sensitivity DNA ScreenTape Analysis (Agilent #5067).

For the CRISPRa screens, we used the 10X Genomics' Chromium Next GEM Single Cell 3' v3.1 HT Dual Index Reagents. Each cell population was diluted to 1000 cells/uL and the Next GEM Chip M was loaded to recover approximately 20K cells per lane. The remainder of the library preparation was according to the manufacturer's protocol (CG000416, Rev A). Library concentrations and sizes were obtained using the KAPA Library Quantification Kit (Roche #7960255001) and High Sensitivity DNA ScreenTape Analysis (Agilent #5067).

**gRNA library PCR**

To recover the gRNA protospacer sequences in each cell, we performed a tri-nested PCR of the cDNA as previously described[32] using primers listed in **Supplementary Table 14**. For the CRISPRi screens, 4 uL of cDNA from each sublibrary was input into a 50 uL reaction with KAPA HiFi and PCR primers prLRB470 and prLRB471. For the CRISPRa screens, 4 uL of cDNA from each sublibrary (8 uL total) was input into a 50 uL reaction with KAPA HiFi and PCR primers prLRB470 and prLRB471. The reaction was amplified and removed prior to saturation, then purified using a 1:1 ratio of AMPure XP DNA beads and eluted in 25 uL H20. 1 uL of the purified sample was input into reaction 2 using PCR primers prLRB472 and prLRB473. The reaction was amplified and removed prior to saturation, then purified as described above. 1 uL of the purified sample was input into reaction 3 using PCR primers prLRB473 and prLRB289-295,298-314, amplifying each sample with a unique i7 sequencing index. The reaction was amplified and removed prior to saturation, then purified using a 1:1 ratio of AMPure XP DNA beads (Beckman Coulter #A63881), and eluted in 25 uL Buffer EB (Qiagen #19086). Library concentrations and sizes were obtained using the KAPA Library Quantification Kit (Roche #7960255001) and High Sensitivity DNA ScreenTape Analysis (Agilent #5067).

**Targeted enrichment**

Probe design

We obtained the Gencode v34 Comprehensive Gene Annotation using the UCSC Table Browser utility. For every transcript within the screen region and for transcripts of the positive control genes, we selected the 300bp region upstream of the transcription end site and input these target regions into the Agilent SureDesign software with the following design parameters: Species: H. sapiens; Build Version: UCSC hg38, GRCh38, December 2013; Category: SureSelect DNA; Hybridization: 90 Minutes; Boosting: Optimized Performance - 90 Minutes; Tiling Density: 2X. The final design included 380 target regions covering 85.531kbp. The final probe design contained 2565 probes covering 84.871 kbp (93.71% coverage).

Hybridization of single cell gene expression libraries

To enrich the gene expression libraries for genes of interest, we used the SureSelect XT HS2 Target Enrichment (Agilent #5191-6900) according to the manufacturer's protocol (Protocol version VA1, September 2020). 200 ng of each library was used as input to each hybridization reaction. Library concentrations and sizes were obtained using the KAPA Library Quantification Kit (Roche #7960255001) and High Sensitivity DNA ScreenTape Analysis (Agilent #5067).

**Sequencing**

For CRISPRi screens, CMO, gRNA, and gene expression libraries for each cell line were pooled and sequenced on an Illumina NovaSeq 6000 S4 flow cell with 2x100bp configuration. For CRISPRa screens, gRNA and gene expression libraries for each cell line were pooled and sequenced on an Illumina NovaSeq 6000 S4 flow cell with 2x100bp configuration. Unenriched gene expression libraries were pooled and sequenced on an Illumina NovaSeq 6000 S4 flow cell with 2x100bp configuration. We also pooled and sequenced a subset of the unenriched gene expression libraries (N=3 per CRISPRi screen) on an Illumina NovaSeq 6000 S4 flow cell with 2x100bp configuration to evaluate the targeted enrichment panel performance.

**Individual gRNA validations**

<u>iPSCs</u>

All experiments were performed with 3-4 biological replicates per condition. Cells were seeded onto Matrigel-coated 24-well plates at 35,000 cells per well. 24 hours later, 25 uL of 20X lentivirus was added per well. 24 hours post-transduction, lentivirus was removed and media was changed. 24 hours later (48 hours post-transduction), media was changed and selection for transduced cells was started using 2.5 ug/mL Blasticidin S HCl (ThermoFisher #A1113903). Selection was continued until cells were harvested. All conditions were passaged once prior to final harvest. The cells were harvested and mRNA was purified using the Qiagen RNeasy kit according to the manufacturer's protocol. 50 ng mRNA was used as input for cDNA amplification using the Invitrogen™ SuperScript™ VILO™ cDNA Synthesis Kit (ThermoFisher #11754050). For RT-qPCR, each reaction contained 2 uL cDNA, 6 uL H20, 1 uL Taqman probe for 18S, 1 uL Taqman probe for gene of interest, and 10 uL PerfeCTa FastMix II (Quantabio #95118). Delta delta Ct analysis was performed in Microsoft Excel. Graphpad Prism was utilized to conduct one-way ANOVA tests followed by Tukey's HSD for post-hoc testing. Significance is reported in figures as follows: *p-value < 0.05, **p-value < 0.01, ***p-value < 0.001. Taqman probe information is provided in **Supplementary Table 15**.

<u>NPCs</u>

All experiments were performed with 3-4 biological replicates per condition. Cells were seeded onto Matrigel-coated 24-well plates at 50,000 cells per well. 24 hours later, 25 uL of 20X lentivirus was added per well. 24 hours post-transduction, lentivirus was removed and media was changed. 24 hours later (48 hours post-transduction), media was changed and selection for transduced cells was started using 2.5 ug/mL Blasticidin S HCl (ThermoFisher #A1113903). Selection was continued until cells were harvested. All conditions were passaged once prior to final harvest. The cells were harvested and mRNA was purified using the Qiagen RNeasy kit according to the manufacturer's protocol. 50 ng mRNA was used as input for cDNA amplification using the

Invitrogen™ SuperScript™ VILO™ cDNA Synthesis Kit (ThermoFisher #11754050). For RT-qPCR, each reaction contained 2 uL cDNA, 6 uL H20, 1 uL Taqman probe for 18S, 1 uL Taqman probe for gene of interest, and 10 uL PerfeCTa FastMix II (Quantabio #95118). Delta delta Ct analysis was performed in Microsoft Excel. Graphpad Prism was utilized to conduct one-way ANOVA tests followed by Tukey's HSD for post-hoc testing. Significance is reported in figures as follows: *p-value < 0.05, **p-value < 0.01, ***p-value < 0.001. Taqman probe information is provided in **Supplementary Table 15**.

K562s

All experiments were performed with 3-4 biological replicates per condition. 25,000 cells were resuspended in 0.5 mL of media containing 25 uL of lentivirus and 8 ug/mL polybrene, then centrifuged for 30 minutes at 25C. The volume was transferred into one well of a 24-well plate. 24 hours post-transduction, the cells were centrifuged for 5 minutes at 300g, resuspended in fresh media, and plated into a 6-well plate. Cells were then selected with 10 ug/mL Blasticidin S HCl (ThermoFisher #A1113903) until final harvest. mRNA was purified using the Qiagen RNeasy kit according to the manufacturer's protocol. 50 ng mRNA was used as input for cDNA amplification using the Invitrogen™ SuperScript™ VILO™ cDNA Synthesis Kit (ThermoFisher #11754050). For RT-qPCR, each reaction contained 2 uL cDNA, 6 uL H20, 1 uL Taqman probe for 18S, 1 uL Taqman probe for gene of interest, and 10 uL Quantabio PerfeCTa FastMix II. Delta delta Ct analysis was performed in Microsoft Excel. Graphpad Prism was utilized to conduct one-way ANOVA tests followed by Tukey's HSD for post-hoc testing. Significance is reported in figures as follows: *p-value < 0.05, **p-value < 0.01, ***p-value < 0.001. Taqman probe information is provided in **Supplementary Table 15**.

**Single cell screen analysis pipeline**

Data processing

All data processing steps were performed using CellRanger v6.0.1 and the human reference genome ('refdata-gex-GRCh38-2020-A') was downloaded from 10X Genomics' software downloads webpage. FASTQ files for each flow cell lane and sequencing run were generated from .bcl files using the cellranger `mkFASTQ` pipeline. The corresponding FASTQ files for each sample were then merged. For the CRISPRi screens, the merged FASTQ files for the expression libraries were then processed using the cellranger `multi` pipeline and aggregated using the cellranger `aggr` command. For the CRISPRa screens, the merged FASTQ files for the expression libraries were processed using the cellranger `count` pipeline with the number of expected cells specified (`--expect-cells = 20000`) and aggregated using the cellranger `aggr` command. The merged FASTQ files for the gRNA libraries were processed using the cellranger `count` pipeline with the number of expected cells specified (`--expect-cells = 30000` for CRISPRi and `--expect-cells = 20000` for CRISPRa). The processed reads were then aligned to a custom bowtie index containing all protospacer sequences included in the pooled gRNA library and the UMI counts corresponding to each gRNA-cell pair were obtained.

gRNA assignment

Guide assignment is determined with a mixture model comprising two distributions. The Poisson distribution at 0 UMI models the cells in which the gRNA is not actually expressed by that cell, while the negative binomial models the cells in which the gRNAs should be assigned. A normalized cell-specific parameter allows for confounding technical factors that affect individual cells such as sequencing depth and batch effects. These cell-specific values contain information about relative cell gRNA UMI library size and are used to normalize the mean values of the two distributions. To increase the speed and efficiency of the model, the model analyzes non-zero gRNA counts and conditions on the input gRNA count being larger than zero. The priors applied to latent variables in the mixture model are generated using ground truth understanding of the ambient and signal gRNAs. The model produces a probability value for each gRNA-cell pair, which is the probability of the pair in the Poisson distribution over the negative binomial distribution. A gRNA is assigned to a cell when the probability value for the gRNA-cell pair is greater than 0.5.

Differential expression testing

The gene expression and gRNA UMI count data was imported into Seurat v3.1. We defined thresholds and filtered for quality cells separately for each experiment based on mitochondrial read percentage, total UMIs, and total genes detected, per cell, as follows:

*Enriched: KRAB K562:* `nFeature_RNA > 20 & nCount_RNA < 3000 & percent.mt < 20`

*Enriched: KRAB iPSC, KRAB NPC, p300 NPC:* `nFeature_RNA > 20 & nCount_RNA < 1500 & percent.mt < 20`

*Enriched: P300 iPSC:* `nFeature_RNA > 20 & nCount_RNA < 1500 & nCount_RNA > 220 & percent.mt < 20`

*Unenriched KRAB K562, KRAB iPSC, and KRAB NPC:* `nCount_RNA > 5000 & percent.mt < 20`

Differential expression analysis was performed in order to identify target genes of each gRNA. For each individual gRNA we compared the transcriptomes of cells where the gRNA was recovered and other perturbed cells. A minimum of 3 cells expressing a gene was required for each gRNA-gene test. Genes detected in less than half of the non-targeting perturbations were independently filtered out in each screen. SCTransform[33] was used to normalize transcript UMI counts. A negative binomial generalized linear model was fitted over the SCTransform adjusted counts to identify significantly differentially expressed genes between the two groups. Beta coefficients for the group variable were used as measurement of effect size.

The target gRNA-gene associated p-values (target p-values) were leveraged to identify significant regulatory regions as follows. First, a non-uniform distribution of p-values was observed in the non-targeting controls, which was linearly interpolated and used to transform the target p-values. Next, for each region, a robust rank aggregation (RRA) method was employed to test whether or not the set of target p-values was significantly different from the expected set, comparing each to a set of 10 million null simulations. Finally, the p-values produced by the RRA method were adjusted for multiple hypothesis testing using the Benjamini-Hochberg procedure. The code to run this method is publicly available[34] and can be found here: https://github.com/Gersbachlab-Bioinformatics/FRACTEL.

#### EP-length calculation

Gene coordinates were obtained from the Ensembl Human Gene v104 reference file. The distance between the CRE and the paired gene was calculated as follows: 1) the CRE midpoint (CRE_mid) was defined as (CRE_start + CRE_end)/2, the gene start coordinate (gene_start) was defined as the start coordinate for genes on the '+' plus strand, and end coordinate for genes on the '-' strand, and 'ep_length' was calculated as CRE_mid - gene_start.

#### Data visualization

All upset plots were generated using the `UpSetR` R package. The circos plot in Figure 1 was generated using the `circos` R package. All other plots were generated using the `ggplot2` R package or using Graphpad Prism.

#### Evaluation of targeted enrichment panel

For the enriched and unenriched datasets, we calculated the mean UMI counts for each detected gene within and not within the MHC locus. We then summed the counts across all genes within the MHC locus and across all genes and calculated the percent of UMIs for genes within the MHC locus relative to all UMIs. The fold-enrichment was then defined as the ratio of the enriched percentage to the unenriched percentage.

#### Quantile-quantile plots

A large number of theoretical quantiles were extracted from a uniform distribution (expected) and the distribution of observed gRNA-gene p-values (observed). Each quantile was compared and plotted across both groups.

#### Comparison of effect sizes by type of gRNA

For the CRISPRi screens, we compared the individual gRNA effects on CLIC1 mRNA expression for the three most significant gRNAs between gRNA types (targeting, TSS-targeting, nontargeting) by performing a One-way ANOVA test with the `aov` function in R followed by posthoc testing using the `TukeyHSD` function in R. For all screens, we compared the individual gRNA effects for all detected genes in a given screen between targeting and nontargeting gRNAs using the `t.test` function in R.

#### Single cell screen effect size versus individual gRNA validations in KRAB iPSCs

For the 10 CRE-gene links in KRAB iPSCs, the Pearson correlation and related p-value between the mean change in mRNA expression measured via RT-qPCR (DDCt) of three biological replicates versus the gene expression change observed in the single cell screen were calculated using the `stat_cor` function from the `ggpubr` R package.

#### Correlation of CRISPRi and CRISPRa screens

The Spearman correlation and related p-value of the change in gene expression for CRE-gene connections identified in both CRISPRi and CRISPRa screens within the iPSCs or NPCs was calculated using the `stat_cor` function from the `ggpubr` R package.

#### Correlation between change in gene expression and CRE-gene pair distance.

The Spearman correlation and related p-value of the change in gene expression of a given CRE-gene connection and the log10-transformed distance between the CRE and gene (`log10(abs(ep_length))`) for CRE-gene pairs within a cell type was calculated using the `stat_cor` function from the `ggpubr` R package.

#### Feature enrichment analysis

Datasets used for all related analyses are provided in **Supplementary Table 16**.

Genomic and epigenomic annotations

For iPSCs, peak calls for each annotation in WTC11 iPSCs, iPS DF 6.9, and iPS DF 19.11, were merged to generate a union peak set. For NPCs, peak calls for each annotation in WTC11 NPCs, H1-derived NPCs, and H9-derived NPCs, were merged to generate a union peak set. The targeted cCREs were intersected with the annotations in each cell type using `bedtools intersect`. A Fisher's exact test was then performed using the `fisher.test` function in R comparing the number of CREs and targeted cCREs that overlap or do not overlap the feature.

chromHMM annotations

For each cell type, we intersected the regions with annotated chromatin states with all targeted cCREs using `bedtools intersect`. A Fisher's exact test was then performed using the `fisher.test` function in R comparing the number of CREs and targeted cCREs that overlap or do not overlap each chromatin state annotation.

#### Comparison to other prediction methods

Nearest gene

For each CRE-gene pair, we calculated the number of genes "skipped" by the element to regulate the gene as follows. First, we defined the start and end coordinates for a given CRE and the start and end coordinates, and strand, for the paired gene. Next, we counted the number of genes detected in the gene expression dataset for which the entire gene body was contained within the region between the CRE and the connected gene and repeated this for all CRE-gene connections.

ABC model

The Activity-by-Contact (ABC) model[35] v0.2 pipeline was used to predict CRE-gene connections in iPSC, K562 and NPC cell lines as follows:

1. *Defining candidate elements:* This step is used to define a set of putative enhancer elements for which ABC Scores will be computed. For the purpose of our analysis, we used an already defined set of enhancer elements (all targeted cCREs), instead of the pre-defined `makeCandidateRegions.py` script.
2. *Quantifying enhancer activity:* Enhancer activity was quantified using the `run.neighborhoods.py` in the candidate enhancer regions from the previous step. The inputs used were H3K27ac ChIP-seq bam files, ATAC-seq bam files and a list of differential expressed genes for each cell type. The output files created are `EnhancerList.txt` and `GeneList.txt`, where the former quantifies enhancer activity and the latter counts reads in gene bodies and promoter regions.
3. *Computing ABC score:* The `predict.py script` from the ABC model software package was used to generate the ABC scores using the `EnhancerList.txt` and `GeneList.txt` from the above step and HiCAR data associated with each cell type with all other default parameters.

For each cell type, the ABC predicted connections were filtered for an ABC Score > 0.02 (default threshold) and then were intersected with the CRE-gene pairs in each cell type using `bedtools intersect`.

E2G model

ENCODE E2G[36] is a newly developed supervised classifier model that predicts enhancer-gene regulatory interactions in a given cell type. Using this model, an encyclopedia of enhancer-gene regulatory interactions in the human genome was built as part of ENCODE4 distal-regulatory WG efforts. We obtained the predicted enhancer-gene regulatory interactions for K562 cells, WTC11 iPSCs, and neural progenitor cells, and then intersected them with the CRE-gene pairs in each cell type using `bedtools intersect`.

EnhancerAtlas

For K562s, we used predicted region-gene pairs in K562 cells. For iPSCs, we used predicted region-gene pairs in H1 and H9 ESCs. For NPCs, we used predicted region-gene pairs in ESC-derived neuronal cells. We intersected all targeted cCREs with the predicted regions in each cell type using `bedtools intersect`. We then calculated the number of CRE-gene pairs for which the CRE overlaps the predicted region and the differentially expressed gene was the same gene as in the predicted region-gene pair.

HiCAR

For each cell type, anchors of all chromatin loops were separately intersected with all targeted cCREs using `bedtools intersect`. We then intersected the anchors with a 1kb region centered on the TSS of each gene and intersected the anchors with the entire gene body using `bedtools intersect`. We then calculated the number of CRE-gene pairs for which the CRE overlaps one anchor and the second anchor overlaps either the +/- 1kb region around the TSS or the gene body of the differentially expressed gene.

**Definition of 'shared' and 'cell-type' specific CREs and CRE-gene connections**

'Shared' CREs and CRE-gene connections were defined as those with significant effects (FDR < 0.01) on gene expression in all three cell types. 'Cell-type' specific CREs and CRE-gene connections were defined as those with significant effects (FDR < 0.01) on gene expression in only one cell type.

**Correlation of effect sizes for CRE-gene links**

The Spearman correlation of effect sizes, defined as (1-FDR)*(CRE effect), for shared CRE-gene pairs, cell-type specific CRE-gene pairs, and all cCRE-gene differential expression tests, between each screen were calculated using the `rcorr` function from the Hmisc R package.

**Correlation of basal gene expression**

Bulk RNA-seq TPM values for all detected genes in the screens in all cell types and for all detected genes in the bulk datasets in all cell types were increased by 1 and then log10-transformed. The Spearman correlation of the transformed values was then calculated using the `rcorr` function from the Hmisc R package.

**Comparison of effect sizes for all cCRE-gene tests**

Related to Figure 6, the effect sizes for all cCRE-gene differential expression tests observed in all screens were calculated as (1-FDR)*(CRE effect). A heatmap with the effect sizes was generated using the `pheatmap` function from the `pheatmap` R package with scaling by column (cCRE-gene test) and clustering by rows and by columns.

**Comparison of proportion of CREs and library that overlap ATAC-seq peaks**

The proportions of CREs that overlapped an ATAC-seq peak in a given cell type were compared to the proportion of ATAC-seq peaks in the same cell type included in the original library design using the `prop.test` function in R with the parameters `alternative = "two.sided"` and `correct = TRUE`. This process was then repeated, comparing the overlap of cell-type specific CREs with cell-type specific ATAC-seq peaks within and across cell types.

#### Gene ontology enrichment analysis

In the cell type-specific gene ontology analysis, our approach for identifying differentially expressed (DE) genes is as follows: a CRE link is classified as cell type-specific if FDR falls below the significance threshold for one and only one of the cell types. Additionally, for the gene connected with the CRE, it must be considered expressed in that cell type, while the same gene in at least one of the other cell types must also simultaneously demonstrate both expression (with a mean TPM greater than 1) and accessibility, defined as overlap with ATAC-seq peak(s). A binary indicator was applied, where "1" indicates that the target region overlaps with a peak in the respective cell type's ATAC data. Additionally, we applied a filter to the background genes, selecting only those with a mean TPM greater than 1. For example, consider AIF1 as a DE gene for NPC. This conclusion was made when at least one CRE link involving AIF1 has a -log10(FDR) greater than 1 (FDR < 0.1) for NPC, while both -log(FDR) values for iPSC and K562 are less than 1 for the same CRE link. Furthermore, AIF1 must have a mean TPM exceeding 1 and ATAC peak value of 1 in either K562 or iPSC, or in both. For our gene ontology analysis, we utilized goseq[37] in R/4.1.2. RNA-seq and ATAC-seq data sources are provided in **Supplementary Table 16**.

#### Enrichment of GWAS SNPs in CREs

In this analysis, we investigated the enrichment of significant genome-wide association study (GWAS) SNPs within CREs in the MHC region and their association with gene regulation. We established significance in CRE-gene connections with an FDR threshold of ≤0.001. The primary focus was to ascertain whether there was an overrepresentation of significant GWAS SNPs, defined by a p-value threshold of ≤5e-08, within significant CREs (defined as CREs with at least one CRE-gene connection at q-value ≤0.001). To address the confounding effects of high linkage disequilibrium (LD) among SNPs in the MHC region, we conducted a permutation-based chi-square test. We maintained the original LD structure of SNPs while randomly permuting CRE FDR values to obtain a null distribution of enrichment scores. By comparing the observed enrichment to the permutation-derived null distribution, we evaluated the probability of observing a similar enrichment of significant variants by chance. We then explored a range of FDR (0.001, 0.005, 0.01) and p-value (5e-08, 1e-07, 5e-07, 1e-6) thresholds to assess the robustness of the enrichment signal.

#### Data availability

All datasets used in this study are provided in **Supplementary Table 16**. All datasets generated in this study will be made publicly available and deposited to the IGVF Data Portal.

### Code Availability

All code used in this study will be made available in a public repository prior to publication.

### Materials availability

Addgene catalog numbers for plasmids used in this study are noted in the methods. Primer sequences and gRNA protospacer sequences are reported in **Supplementary Table 14**. All materials used in this study will be made available upon request.

### Author Contributions

Conceptualization: LRB, CAG, GEC, PFS
Methodology: LRB, CAG, GEC, PFS, TR, YL, RG, AA, MIL, PGR
Performed experiments: LRB
Data analysis: LRB, AB, MtW, SL, EW, SL, RV, RR
Visualization: LRB, NI
Funding acquisition: CAG, GEC, PFS
Project administration: LRB, CAG, GEC, PFS
Supervision: CAG, GEC
Writing and revisions: LRB, AB, CAG, GEC, PFS, PGR, RG, MIL, NI

### Funding

The work presented here was supported by National Institutes of Health grants HG011123 (CAG, GEC, TER, and ASA), MH125236 (CAG, GEC, and PFS), HG012053 (CAG, GEC, and TER), HG011967-03 (ASA), HG012003 (MIL), NSF EFMA-1830957 (CAG), and Open Philanthropy (CAG). L.R.B. was supported by the NSF-GRFP (NSF-GRFP DGE - 2139754).

### Acknowledgements

We would like to thank Devi Swain Lenz and the Duke Sequencing Core for excellent assistance with data production, and Dan Somers and the High-throughput Applied Research Data Analysis Cluster (HARDAC) for computing resources. We also thank members of the Allen, Crawford, Gersbach, Majoros, and Reddy Labs at Duke University, and members of the Love, Li, and Sullivan Labs at the University of North Carolina, Chapel Hill, who have provided feedback throughout this project.

### Conflicts of Interest

C.A.G. is an inventor on patents and patent applications related to genome engineering and CRISPR screens, and is a co-founder and advisor to Tune Therapeutics and Sollus Therapeutics, an advisor to Sarepta Therapeutics and Pappas Capital, and a co-founder of Locus Biosciences. L.R.B. is an employee of Xaira Therapeutics (all work performed prior).
