## Supplementary Figures 1-23 for "Functional Annotation of the Major Histocompatibility Complex Locus"

**Supplementary Figure 1. Density of genomic variants in chromosome 6 versus other regions.**

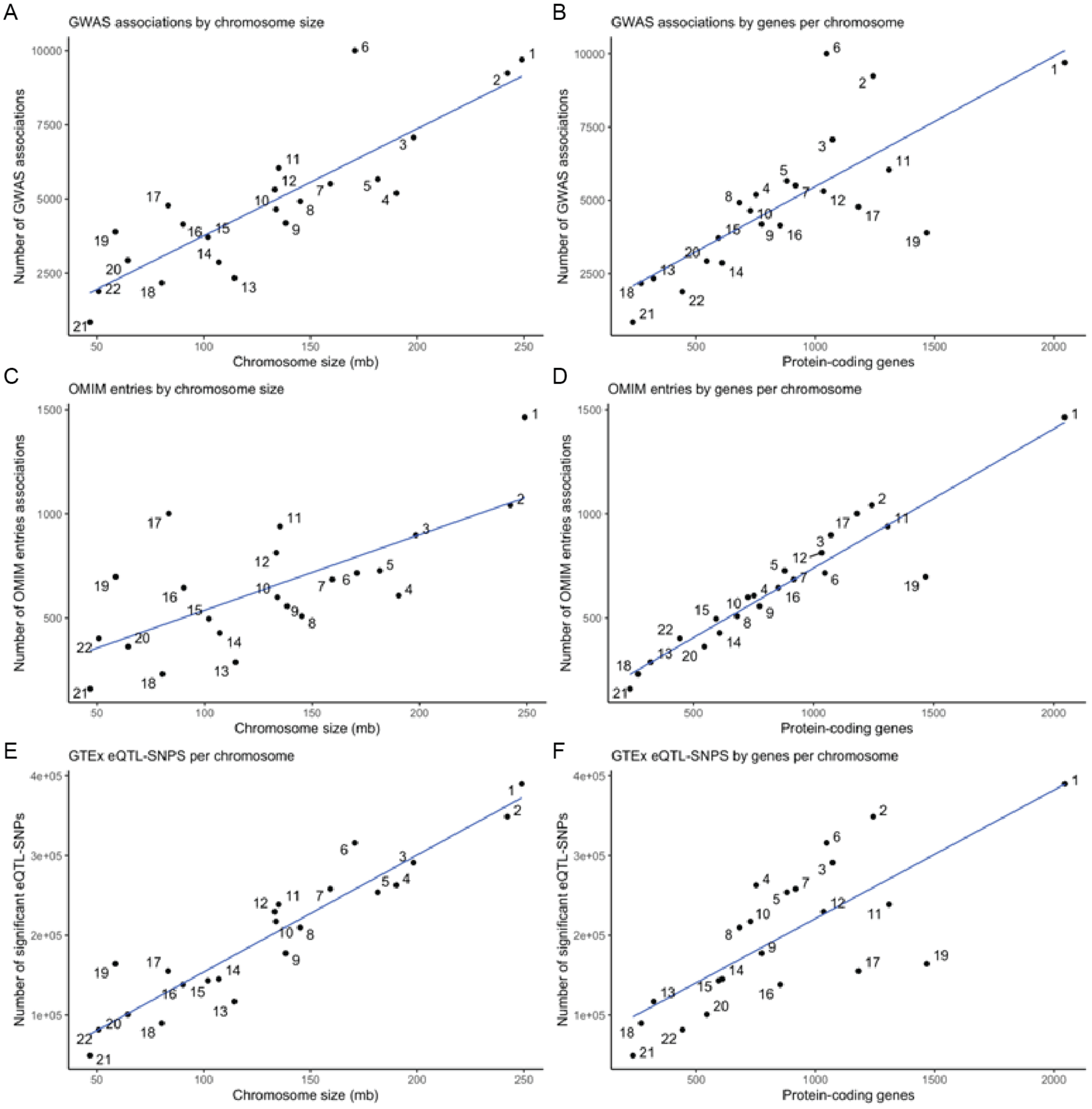


**Supplementary Figure 1. Density of genomic variants in chromosome 6 versus other regions.** Comparison of the chromosome size (Mb) versus the number of **(A)** GWAS associations (NHLBI/EBI GWAS catalog, 10/2021) per chromosome, **(B)** GWAS associations (NHLBI/EBI GWAS catalog, 10/2021) by protein-coding gene per chromosome, **(C)** OMIM entries (10/2021) per chromosome, **(D)** OMIM entries (10/2021) by protein-coding gene per chromosome, **(E)** GTEx (v8) eQTL-SNPs by chromosome, and **(F)** GTEx (v8) eQTL-SNPs by protein-coding gene per chromosome. Each point is a chromosome as labeled. Chromosome 6, which harbors the MHC locus, is a marked outlier for common disease GWAS studies, an outlier for eQTL-SNPs, but not for rare disease OMIM entries.

**Supplementary Figure 2. Gene density, GWAS hit density, and eQTL SNP density on chromosome 6.**

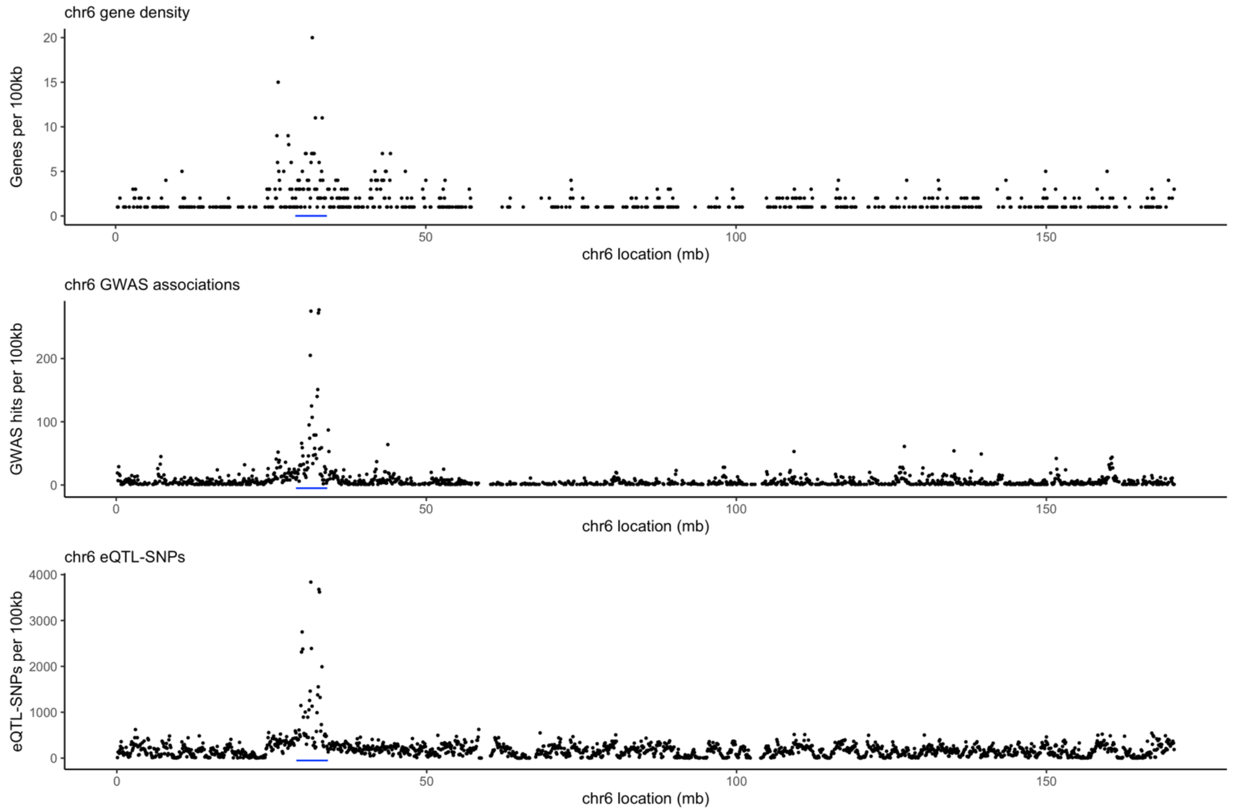


**Supplementary Figure 2. Gene density, GWAS association density, and eQTL SNP density are highly enriched in the MHC locus on chromosome 6.** The MHC region is indicated by the blue bars at the bottom of each graph. The graphs show gene density (top), unique GWAS associated SNPs (middle), and eQTL-SNPs in any GTEx tissue (bottom) (and are also the y-axes). The x-axis is the chromosome 6 location in 100 kb bins.

**Supplementary Figure 3. LD scores on chromosome 6 for different ancestries.**

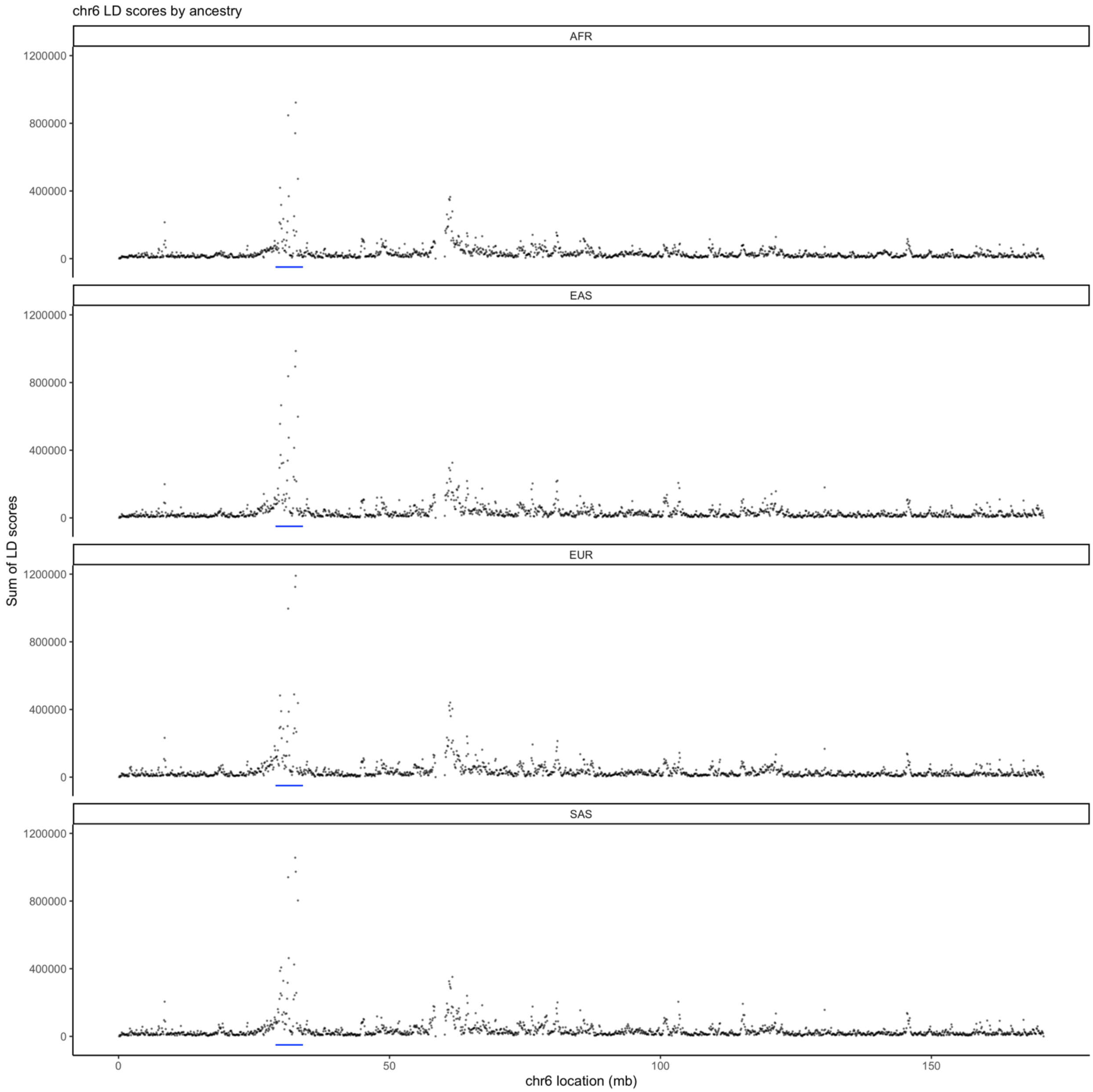


**Supplementary Figure 3. LD scores on chromosome 6 for different ancestries.** The graphs show LD scores on chromosome 6. The x-axis is the chromosome 6 position in Mb and the y-axis is the sum of LD scores per 100 kb bin. The four graphs show LD scores in TOPMed ancestry groups: AFR=African, EAS=East Asian, EUR=European, and SAS=South Asian. The MHC region is indicated by the blue bars at the bottom of each graph. The MHC region has markedly greater LD scores in all ancestries. The smaller peak around 60 Mb brackets the chromosome 6 centromere and results from peri-centromeric suppression of recombination.

**Supplementary Figure 4. Upset plot of targeted cCREs and accessible chromatin regions.**

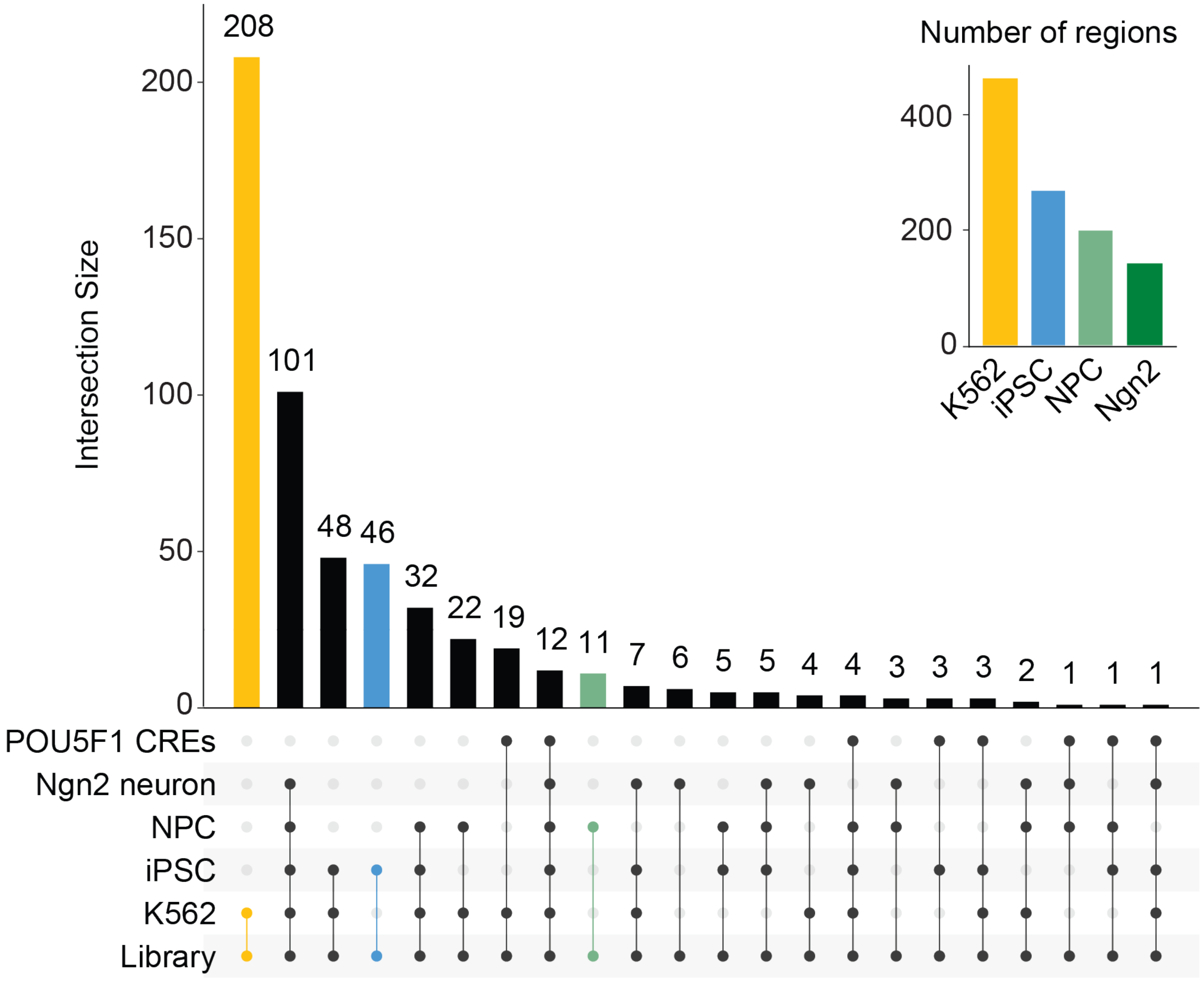


**Supplementary Figure 4. Upset plot of targeted cCREs and accessible chromatin regions.** Upset plot of the ATAC-seq peaks, previously identified *POU5F1* enhancers[1], and the cCREs targeted in the gRNA library design. The total set size of ATAC-seq peaks in each cell type is shown in the upper right and the intersection size for each comparison is shown in the main plot.

**Supplementary Figure 5. dCas9-effector cell line characterization.**

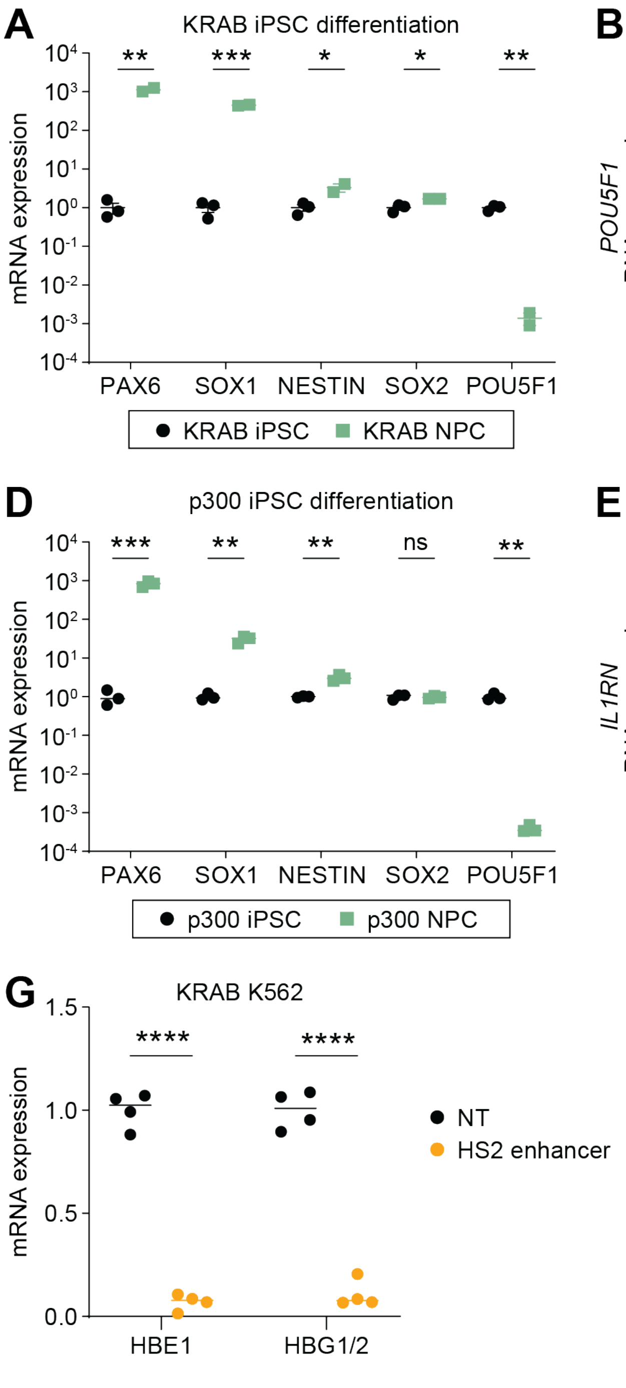


**Supplementary Figure 5. dCas9-effector cell line characterization. (A)** Expression of neural progenitor cell marker genes in dCas9$^{KRAB}$ iPSCs (black) and dCas9$^{KRAB}$ NPCs (green) (N=3) that do not contain any gRNAs. **(B)** *POU5F1* pluripotency gene mRNA expression in dCas9$^{KRAB}$ iPSCs after delivering nontargeting ('NT') gRNA or gRNA targeting the *POU5F1* promoter[4] ('*POU5F1* TSS') (N=3). **(C)** *CHD8* mRNA expression in dCas9$^{KRAB}$ NPCs after delivering nontargeting ('NT') gRNA or gRNA targeting the *CHD8* promoter ('*CHD8* TSS') (N=3). **(D)** Expression of neural progenitor cell marker genes in dCas9$^{p300}$ iPSCs (black) and dCas9$^{p300}$ NPCs (green) (N=3) that do not contain any gRNAs. **(E)** *IL1RN* mRNA expression in dCas9$^{p300}$ iPSCs after delivering nontargeting ('NT') gRNA or gRNA targeting the *IL1RN* promoter[5] ('*IL1RN* TSS') (N=3). **(F)** *IL1RN* mRNA expression after delivering nontargeting ('NT') gRNA or gRNA targeting the *IL1RN* promoter ('*IL1RN* TSS') to dCas9$^{p300}$ NPCs (N=3). **(G)** *HBE1* mRNA expression and *HBG1/2* mRNA expression in dCas9$^{KRAB}$ K562 cells after delivering nontargeting ('NT') gRNA or gRNA targeting the HS2 enhancer[6] (N=3). (**A-G**) Significance values from t-test as follows: *p<0.05, **p<0.01, ***p<0.001, ****p<0.0001.

**Supplementary Figure 6. Targeted enrichment of genes in the 3.5 Mb region of the MHC locus.**

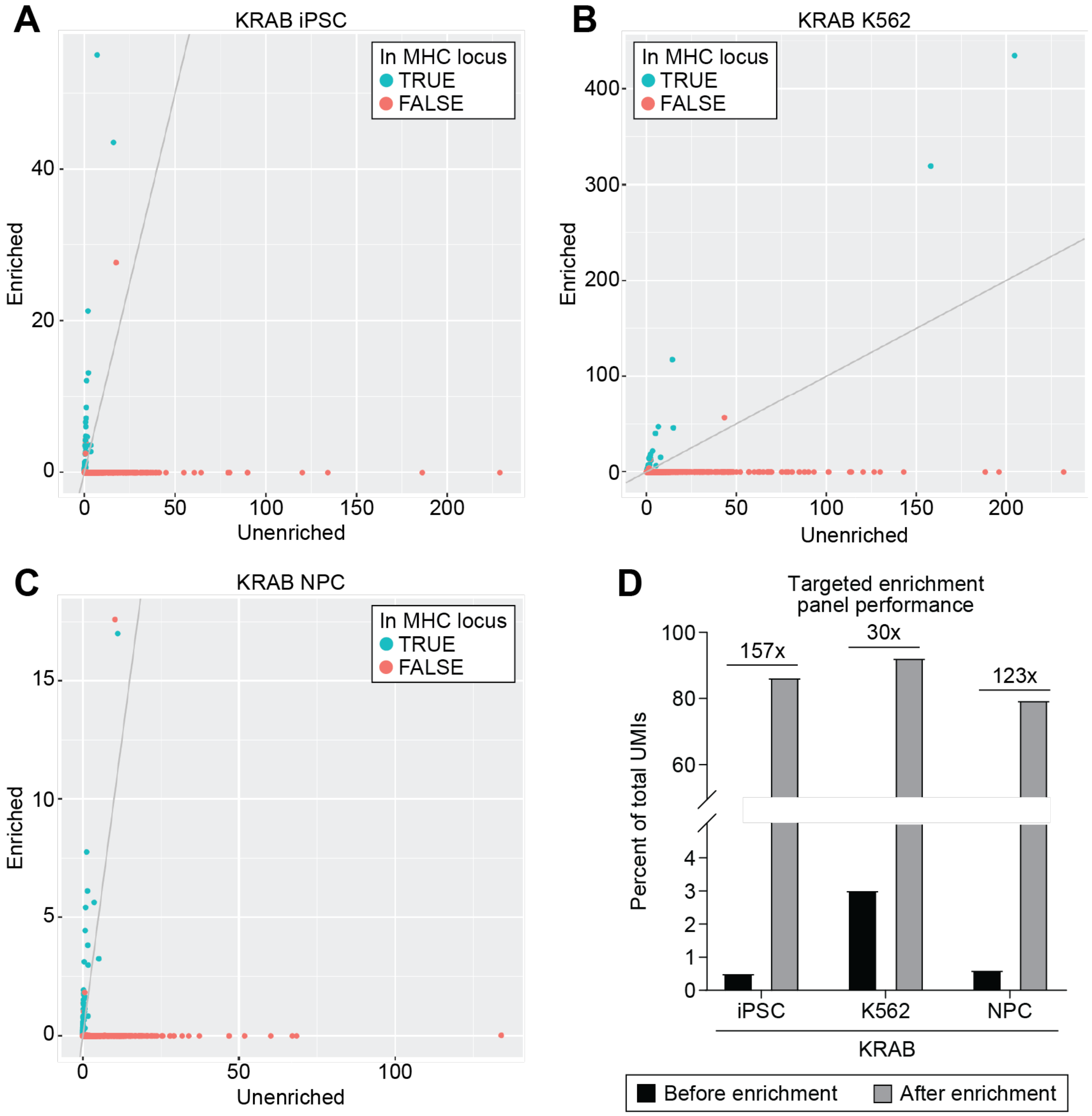


**Supplementary Figure 6. Targeted enrichment of genes in the 3.5 Mb region of the MHC locus.** Scatter plots comparing the mean raw UMI gene counts for genes within (blue) or not within (red) the targeted 3.5 Mb region of the MHC locus before enrichment ('Unenriched') and after enrichment ('Enriched') in the **(A)** dCas9$^{KRAB}$ iPSC, **(B)** dCas9$^{KRAB}$ K562, and **(C)** dCas9$^{KRAB}$ NPC experiments. Grey diagonal lines indicate equal counts. Mean value was calculated from four individual gene expression libraries per experiment. **(D)** Percent of total UMIs for genes within the MHC locus before enrichment (black) and after enrichment (grey). The percentage is calculated from four individual gene expression libraries per experiment, with fold enrichment noted above each pair of bars.

**Supplementary Figure 7. MOI and coverage histograms for CRISPRi screens.**

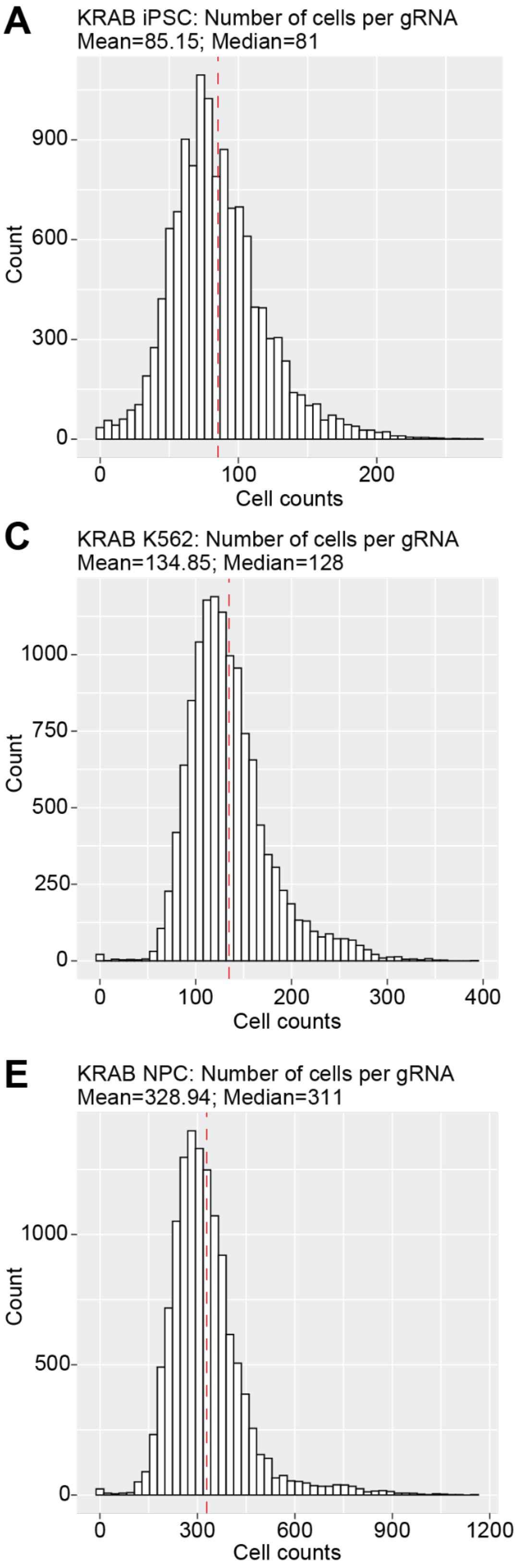

**Supplementary Figure 7. MOI and coverage histograms for CRISPRi screens.** Number of cells in which each gRNA was observed (‘coverage’) and the number of gRNAs observed per cell (‘MOI’) for **(A-B)** dCas9$^{KRAB}$ iPSCs, **(C-D)** dCas9$^{KRAB}$ K562s, and **(E-F)** dCas9$^{KRAB}$ NPCs. Mean and median values are noted above each histogram and the red dashed line indicates the mean value.

**Supplementary Figure 8. MOI and coverage histograms for CRISPRa screens.**

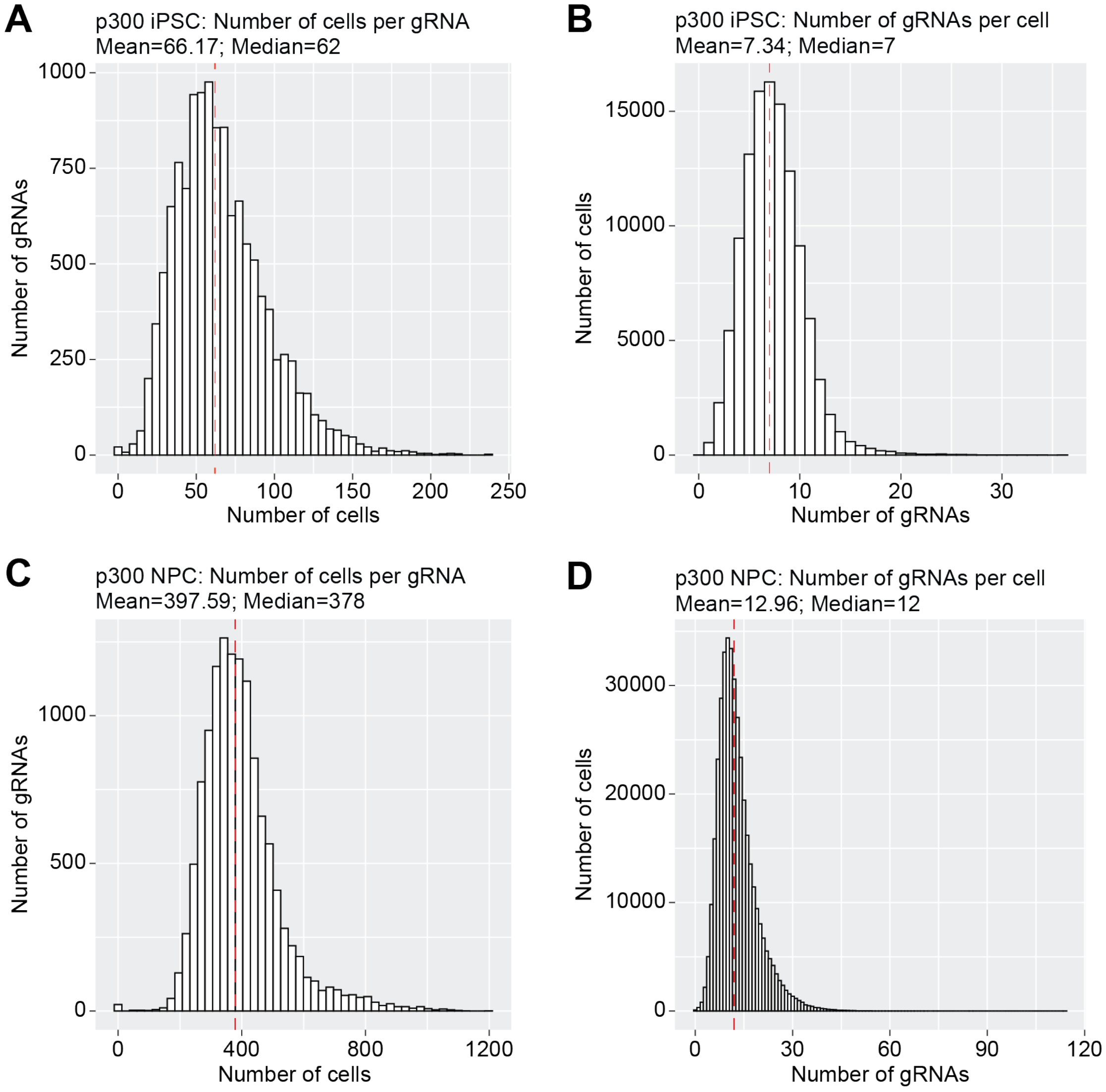


**Supplementary Figure 8. MOI and coverage histograms for CRISPRa screens.** Number of cells in which each gRNA was observed ('coverage') and the number of gRNAs observed per cell ('MOI') for (**A-B**) dCas9$^{p300}$ iPSCs and (**C-D**) dCas9$^{p300}$ NPCs. Mean and median values are noted above each histogram and the red dashed line indicates the mean value.

**Supplementary Figure 9. Quantile-quantile plots of gRNA perturbations.**

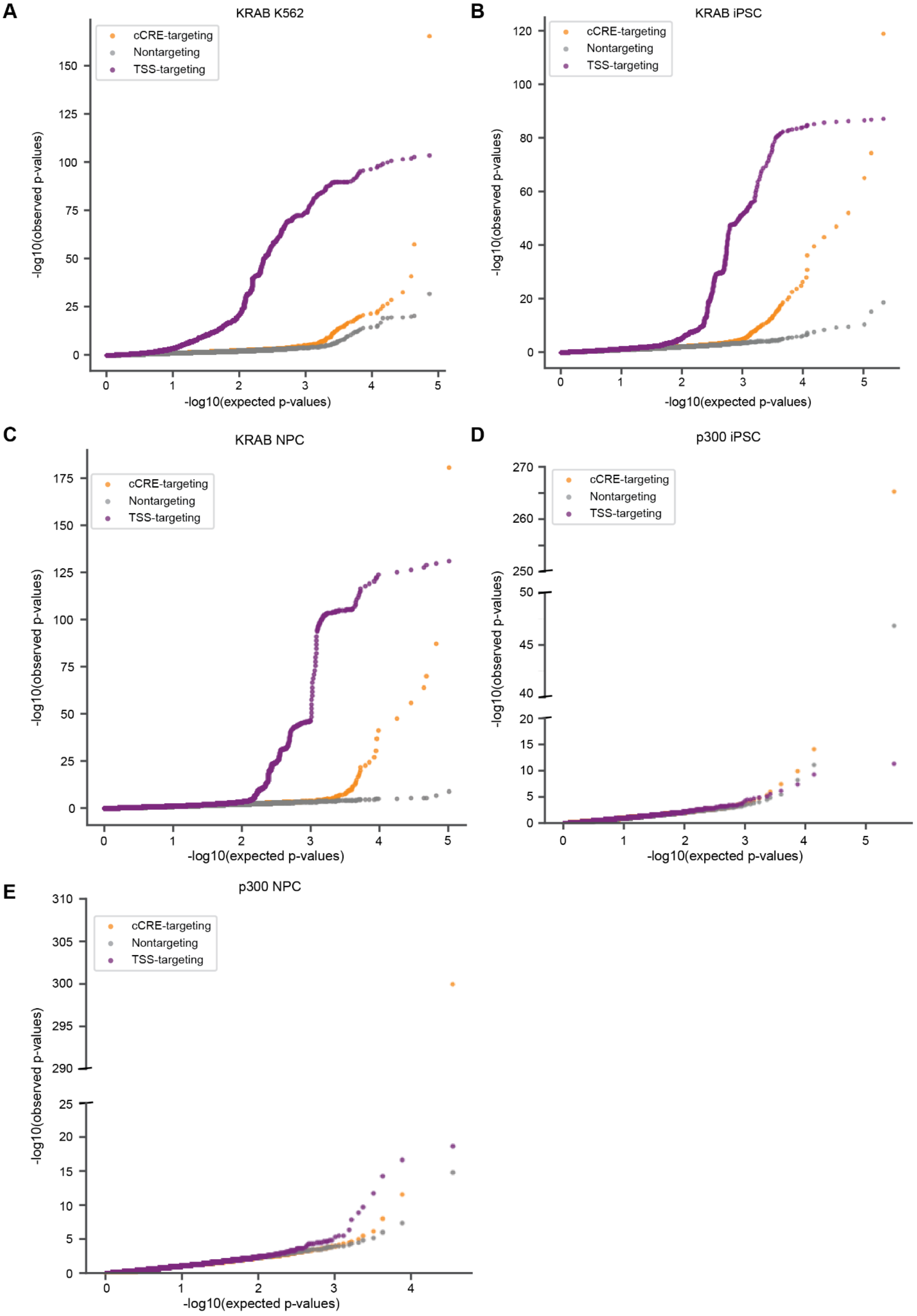

**Supplementary Figure 9. Quantile-quantile plots of gRNA perturbations.** Quantile-quantile plots comparing the -log10-transformed observed p-values and -log10-transformed expected p-values of gRNA perturbations in **(A)** dCas9$^{KRAB}$ K562 cells, **(B)** dCas9$^{KRAB}$ iPSCs, **(C)** dCas9$^{KRAB}$ NPCs, **(D)** dCas9$^{p300}$ iPSCs, and **(E)** dCas9$^{p300}$ NPCs. **(A-E)** TSS-targeting positive controls ('TSS-targeting'), cCRE-targeting, and nontargeting gRNAs are colored in purple, orange, and grey, respectively.

**Supplementary Figure 10. Comparison of control and cCRE-targeting gRNA perturbation effects on gene expression.**

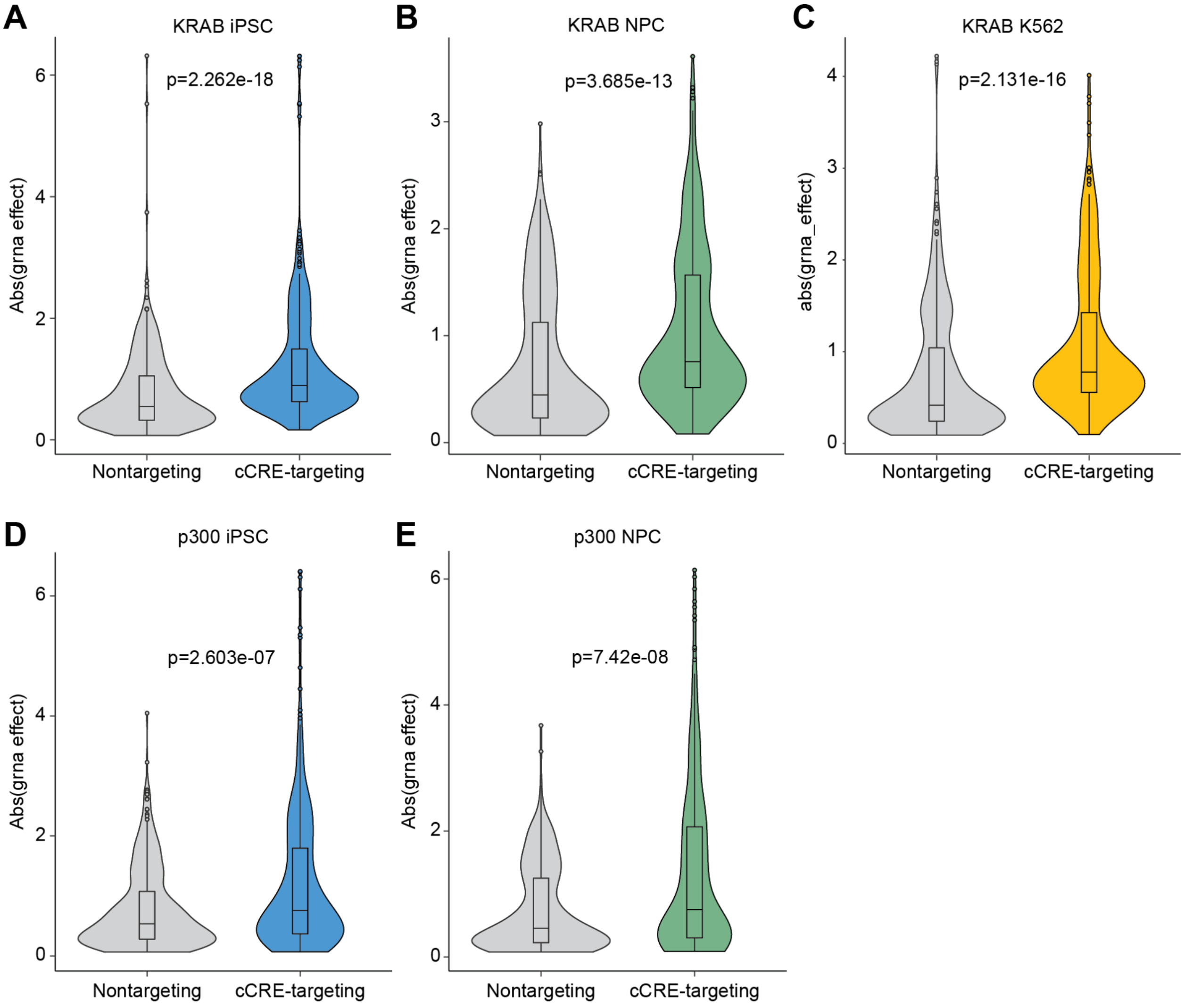


**Supplementary Figure 10. Comparison of control and cCRE-targeting gRNA perturbation effects on gene expression**. Absolute change in gene expression of top 3 gRNAs per cCRE-gene test (ordered by p-value) for nontargeting versus cCRE-targeting gRNAs in the **(A)** dCas9$^{KRAB}$ iPSC, **(B)** dCas9$^{KRAB}$ NPC, **(C)** dCas9$^{KRAB}$ K562, **(D)** dCas9$^{p300}$ iPSC, and **(E)** dCas9$^{p300}$ NPC experiments. P-values from Wilcoxon test are noted in each plot.

**Supplementary Figure 11. Correlation between change in gene expression and distance between CREs and paired genes.**

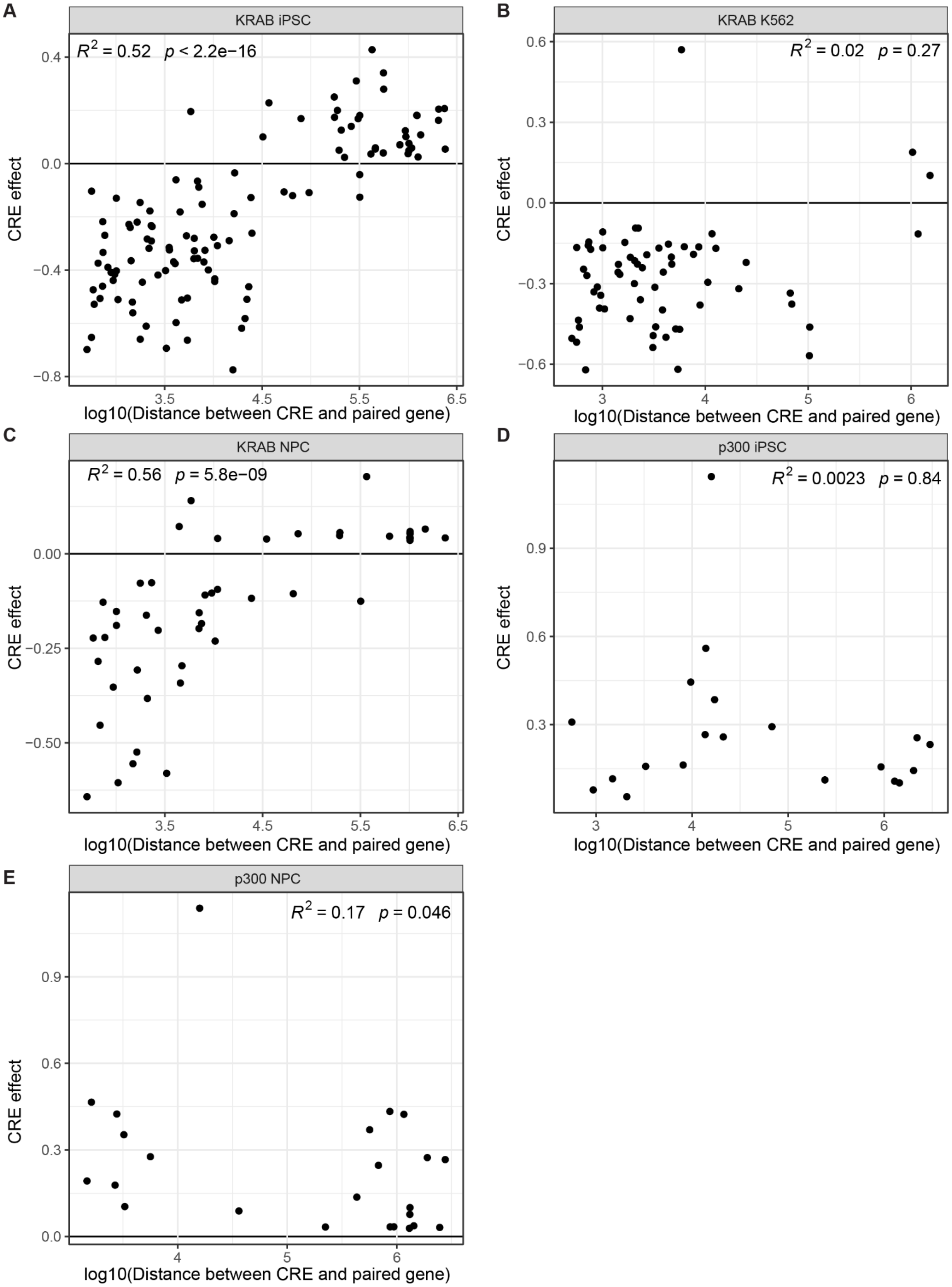


**Supplementary Figure 11. Correlation between change in gene expression and distance between CREs and paired genes.** Scatterplot comparing the change in gene expression for a given CRE-gene pair observed in the screen (‘CRE effect’) and the log10-transformed distance between the CRE and paired gene for all CRE-gene pairs in the **(A)** KRAB iPSC, **(B)** KRAB K562, **(C)** KRAB NPC, **(D)** p300 iPSC, and **(E)** p300 NPC experiments. Spearman correlation and related p-values are noted in each plot.

**Supplementary Figure 12. Example of a promoter that functions as a CRE of a nearby gene.**

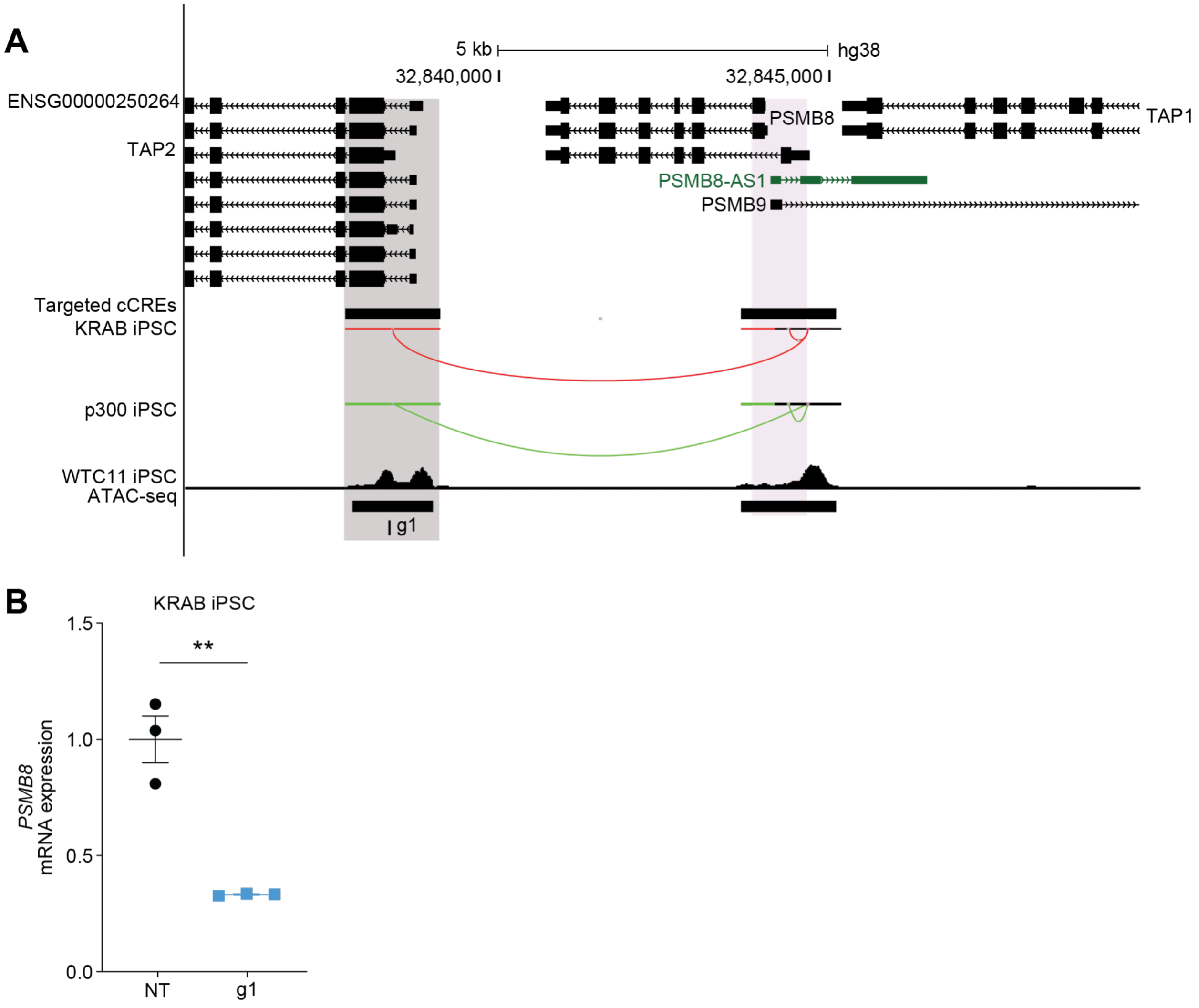


**Supplementary Figure 12. Example of a promoter that functions as a CRE of a nearby gene.**
**(A)** Browser track of the region containing the TAP2 promoter which functions as a CRE of the nearby gene, *PSMB8*. Gene annotation and targeted cCREs are shown with CRE-gene links in the KRAB and p300 iPSC experiments denoted by arcs drawn from the CRE (grey shading) to the gene (purple shading) and colored by red and green for decrease or increase in gene expression, respectively. ATAC-seq signal and called peaks in WTC11 iPSCs are shown. The gRNA used in **(B)** is shown in the final track. **(B)** Change in PSMB8 expression measured via RT-qPCR after delivering a NT gRNA or a gRNA targeting the *PSMB8* CRE to KRAB iPSCs (N=3). P-value from t-test denoted by ** ($p<0.01$).

**Supplementary Figure 13. Comparison of basal gene expression and perturbation effect between CRISPRi and CRISPRa.**

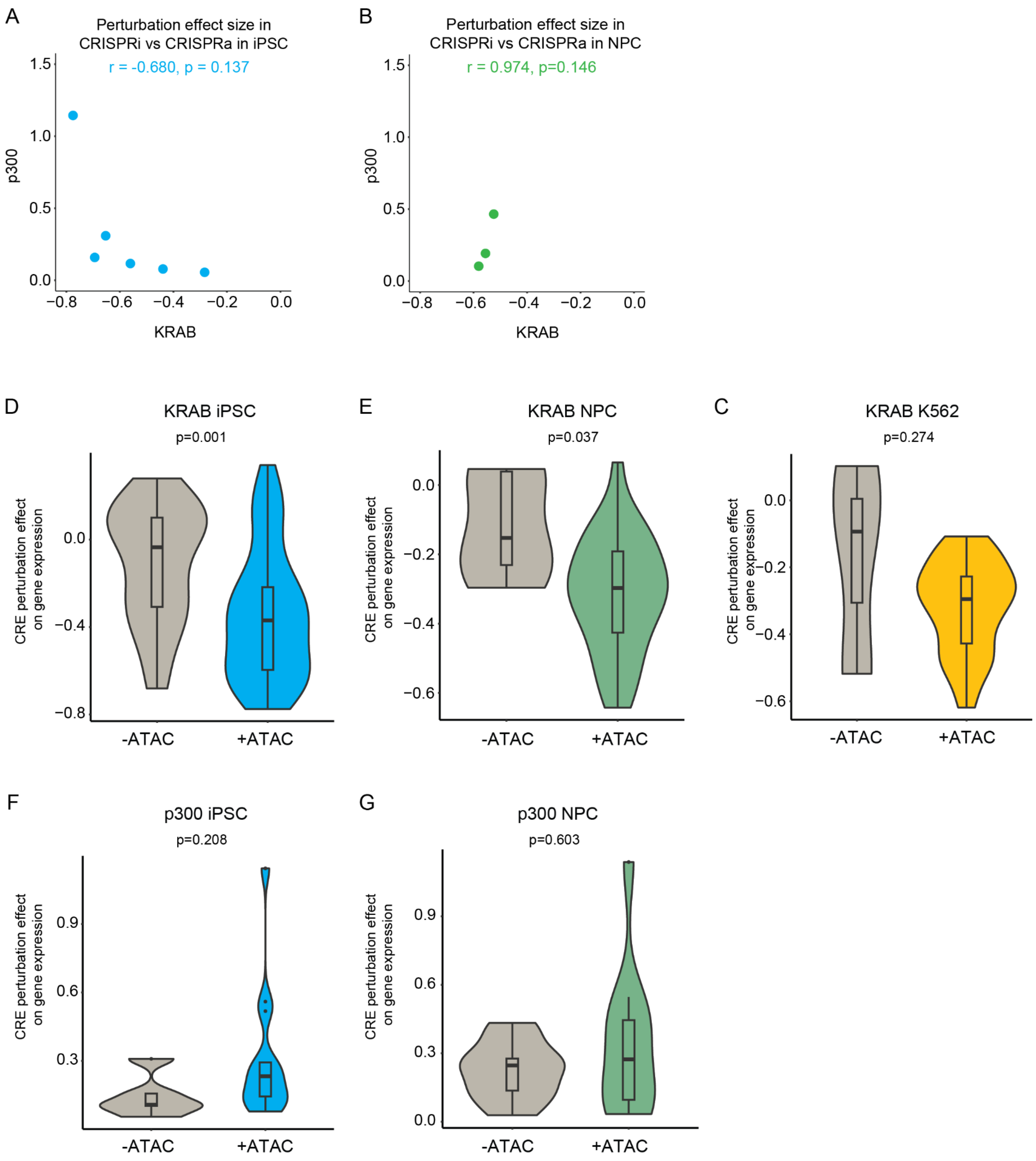


**Supplementary Figure 13. Comparison of perturbation effect on gene expression between CRISPRi and CRISPRa. (A-B)** Scatterplot comparing the change in gene expression for all CRE-gene pairs (FDR < 0.01) identified with both perturbation modalities between KRAB and p300 screens in **(A)** iPSCs and **(B)** NPCs. Pearson correlation coefficient and related p-value are noted in each plot. All shared CRE-gene pairs caused opposite changes in gene expression. **(C-G)**

Comparison of perturbation effect on gene expression between CREs that do not ('-ATAC') or do ('+ATAC') overlap ATAC-seq peaks in **(C)** KRAB iPSCs, **(D)** KRAB NPCs, **(E)** p300 iPSCs, **(F)** p300 NPCs, and **(G)** KRAB K562. The p-value from the Wilcoxon test comparing -ATAC versus +ATAC is noted in each plot.

**Supplementary Figure 14. CREs display diverse combinations of genomic and epigenomic features.**

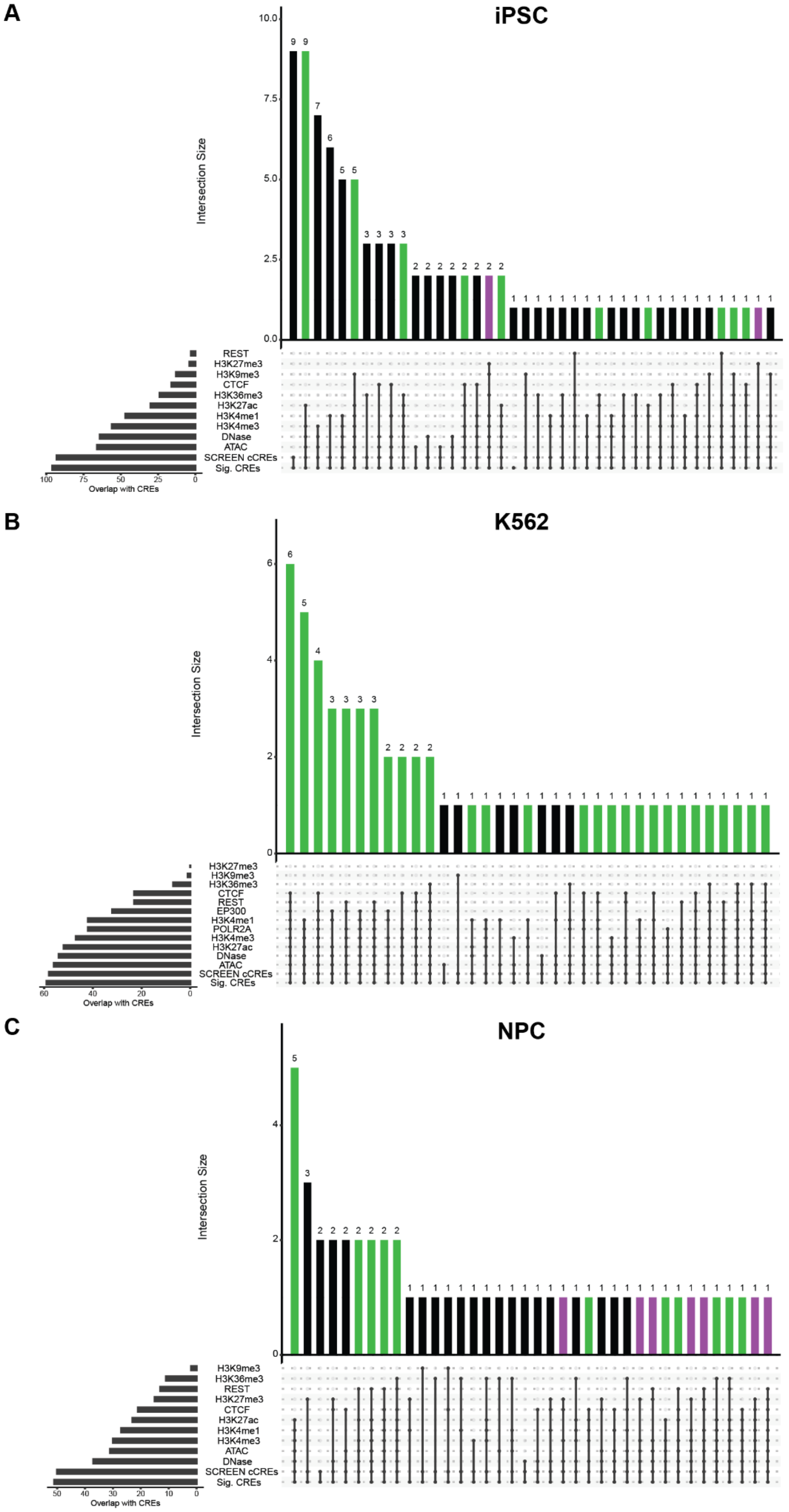

**Supplementary Figure 14. CREs display diverse combinations of genomic and epigenomic features.** Upset plot of intersection of CREs in **(A)** iPSCs, **(B)** K562s, and **(C)** NPCs, with various genomic and epigenomic annotations in the corresponding cell types. Purple shading of bars in the main plot indicates CREs marked by accessible chromatin, H3K27me3, and H3K4me1, in that cell type. Green shading of bars in the main plot indicates CREs marked by accessible chromatin and H3K27ac in that cell type.

**Supplementary Figure 15. Overlap of CREs with genomic and epigenomic annotations.**

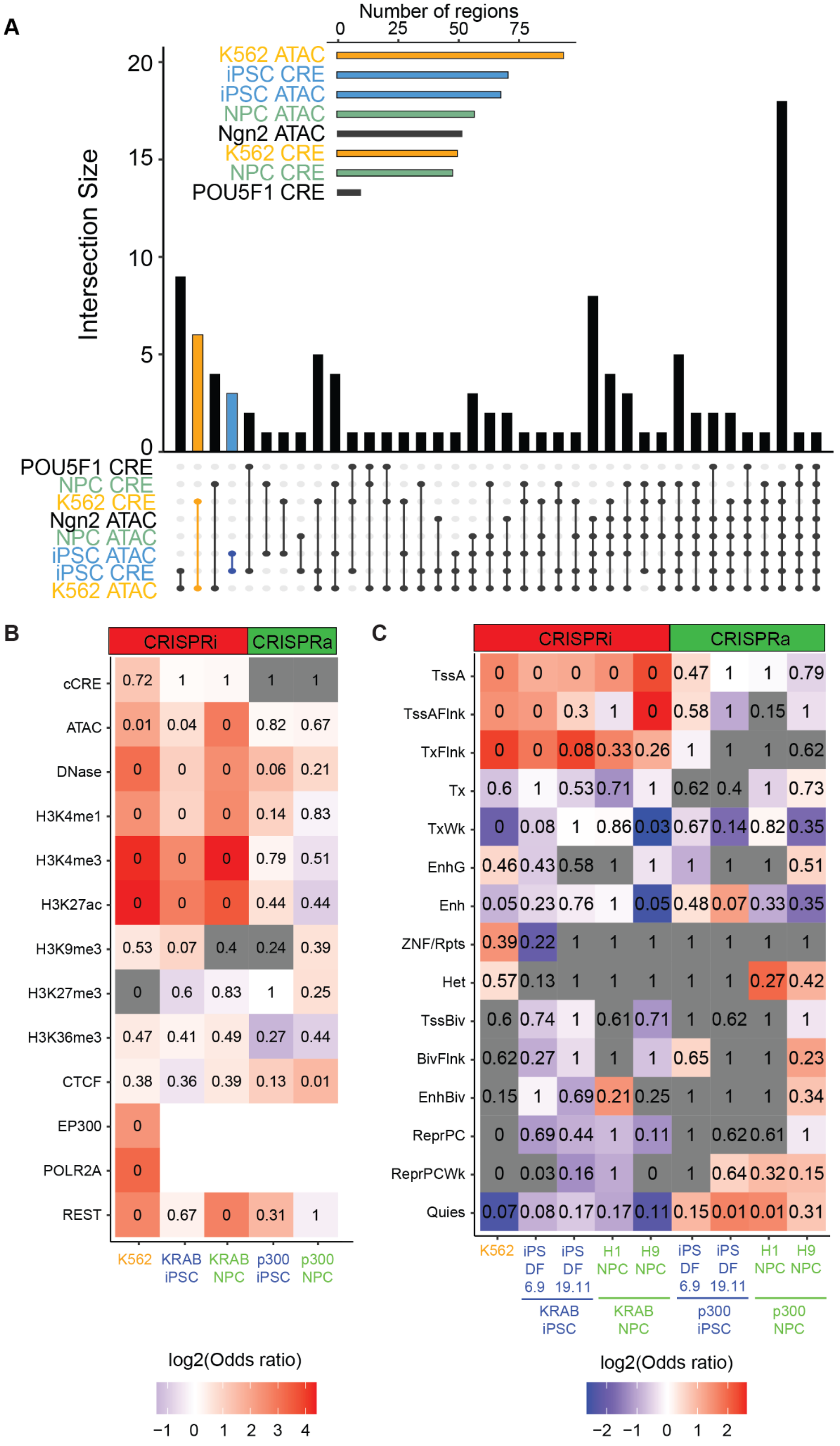

**Supplementary Figure 15. Majority of CREs are marked by accessible chromatin and active histone marks. (A)** Upset plot of CREs and ATAC-seq peaks in all three cell types and targeted cCREs in the library design. The total set sizes are shown in the upper right and the intersection size is shown in the main plot. Bars colored by orange and blue denote the intersections of cell type-specific CREs and cell type-specific ATAC-seq peaks. **(B)** Heatmap of Fisher's exact test results for genomic and epigenomic features of CREs in each cell type. Color indicates log2(Odds ratio) and p-value of Fisher's exact test is shown in each cell. **(C)** Heatmap of Fisher's exact test results for chromHMM annotations of CREs in each cell type. Color indicates log2(Odds ratio) and p-value of Fisher's exact test is shown in each cell. The x-axis labels indicate the cell type source for the chromHMM annotations. Y-axis labels are abbreviated as follows: TssA = Active TSS, TssBiv = Bivalent/poised TSS, BivFlnk = Flanking bivalent TSS/enhancer, EnhBiv = Bivalent enhancer, ReprPC = Repressed polycomb, ReprPCWk = Weak repressed polycomb, Quies = Quiescent/low, TssAFlnk = Flanking TSS, TxFlnk = Strong flanking transcription, Tx = Strong transcription, TxWk = Weak transcription, EnhG = Genic enhancer, Enh = Enhancer, ZNF/Rpts = ZNF genes and repeats, Het = Heterochromatin.

**Supplementary Figure 16. CRISPRi and CRISPRa perturbations reveal putative silencer CREs.**

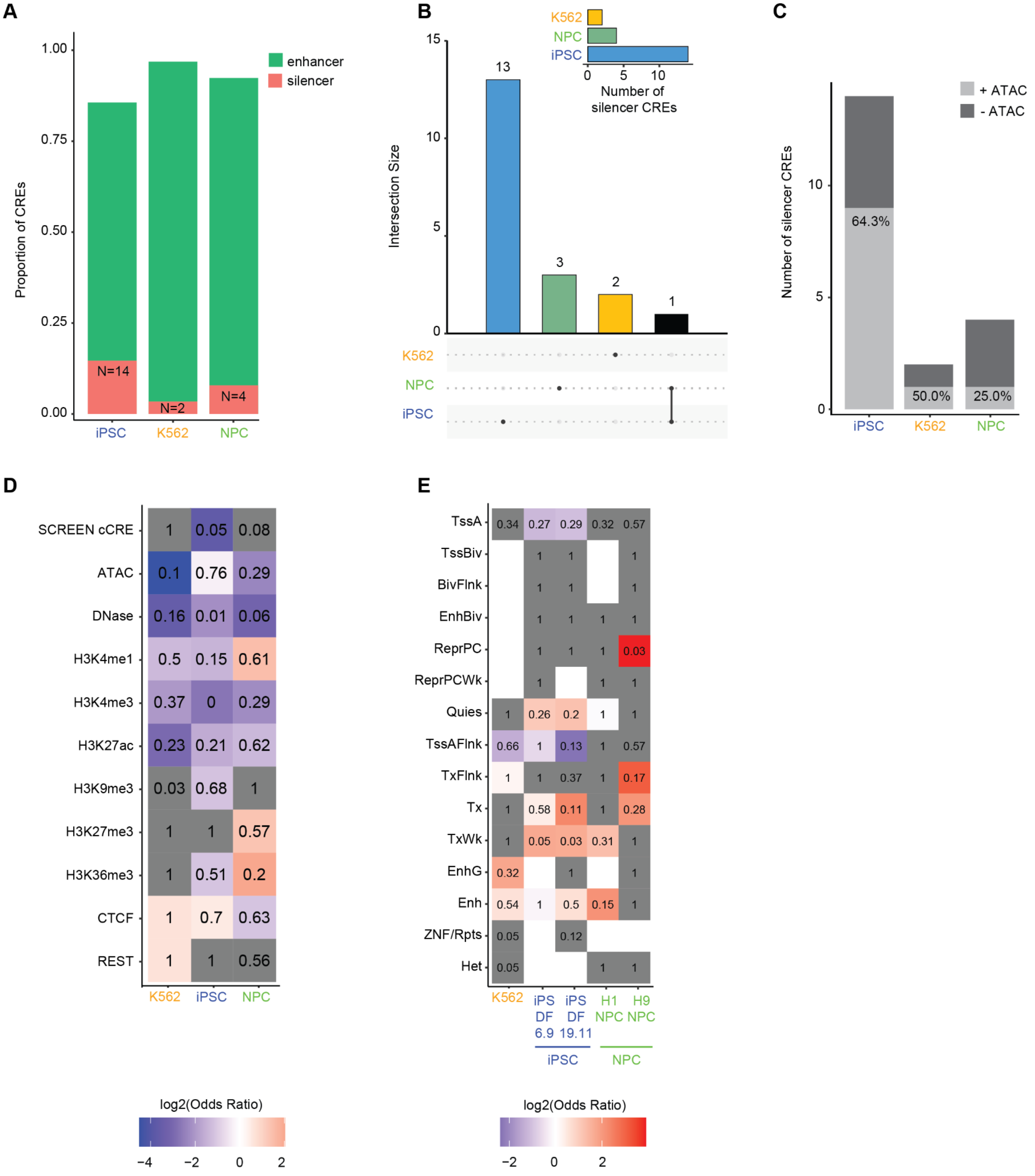


**Supplementary Figure 16. CRISPRi and CRISPRa perturbations reveal putative silencer CREs. (A)** Proportion of CREs defined as enhancers (green) or silencer (red). Count of silencer CREs noted in each bar for each cell type. **(B)** Upset plot of silencer CREs. Red highlight indicates count of silencer CREs identified in all three cell types. **(C)** Number of silencer CREs in iPSCs and NPCs that do (+ATAC) or do not (-ATAC) overlap ATAC-seq peaks in that cell type. The percentage of all silencer CREs that overlap an ATAC-seq peak is noted in each bar. **(D)** Heatmap of Fisher's exact test results for genomic and epigenomic features of silencer CREs in each cell type. Color indicates log2(Odds ratio) and p-value of Fisher's exact test is shown in each cell. **(E)** Heatmap of Fisher's exact test results for chromHMM annotations of silencer CREs in each cell type. Color indicates log2(Odds ratio) and p-value of Fisher's exact test is shown in each cell. The x-axis labels indicate the cell type source for the chromHMM annotations. Y-axis labels are abbreviated as follows: TssA = Active TSS, TssBiv = Bivalent/poised TSS, BivFlnk = Flanking bivalent TSS/enhancer, EnhBiv = Bivalent enhancer, ReprPC = Repressed polycomb, ReprPCWk = Weak repressed polycomb, Quies = Quiescent/low, TssAFlnk = Flanking TSS, TxFlnk = Strong flanking transcription, Tx = Strong transcription, TxWk = Weak transcription, EnhG = Genic enhancer, Enh = Enhancer, ZNF/Rpts = ZNF genes and repeats, Het = Heterochromatin.

**Supplementary Figure 17. E2G model performs better for CRISPRi than CRISPRa experiments.**

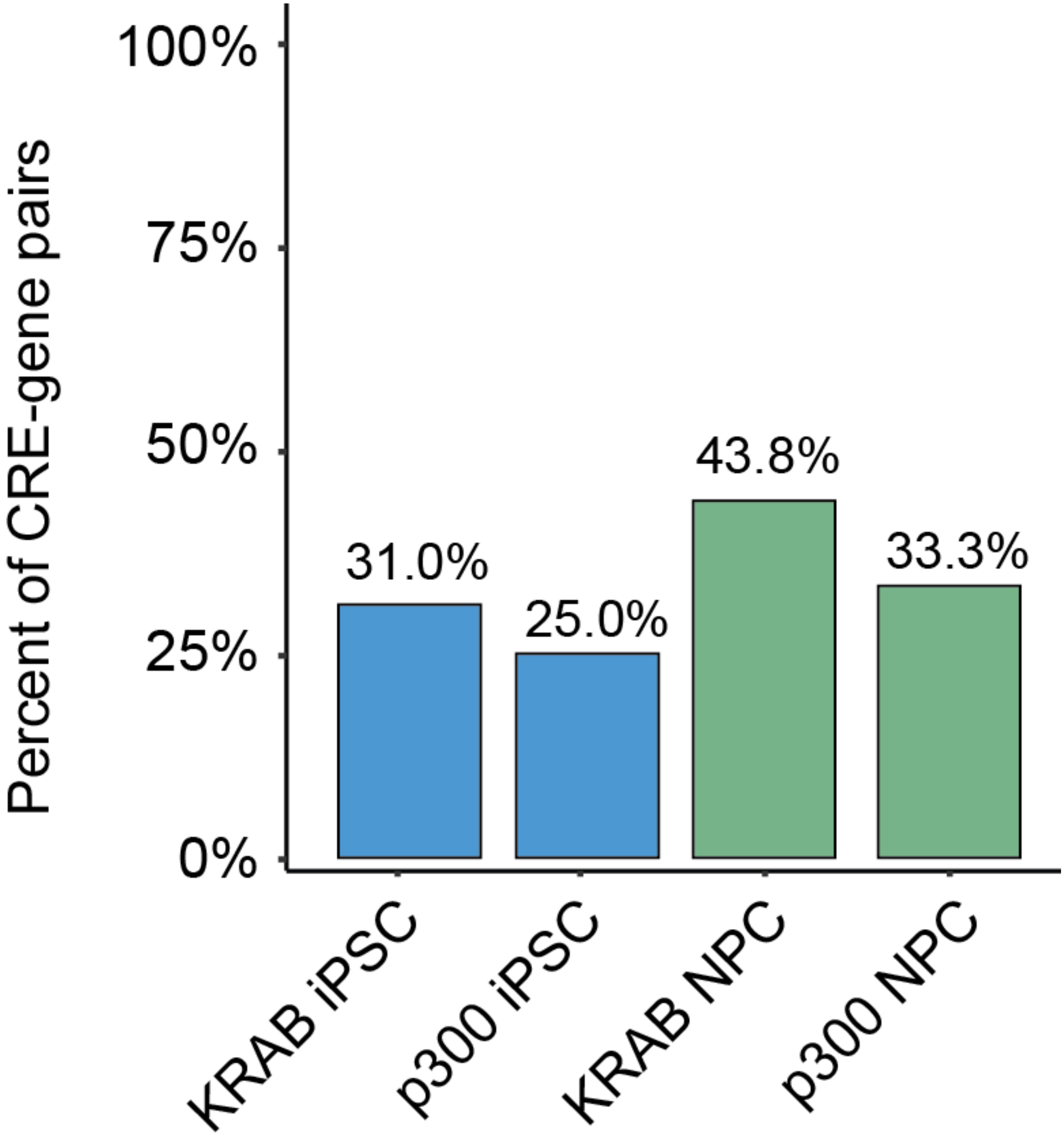


**Supplementary Figure 17. E2G model performs better for CRISPRi than CRISPRa experiments.** Percent of CRE-gene pairs predicted by the E2G model for CRISPRi and CRISPRa experiments in iPSCs and NPCs.

**Supplementary Figure 18. CRE-gene pair significance is not correlated with ABC Score.**

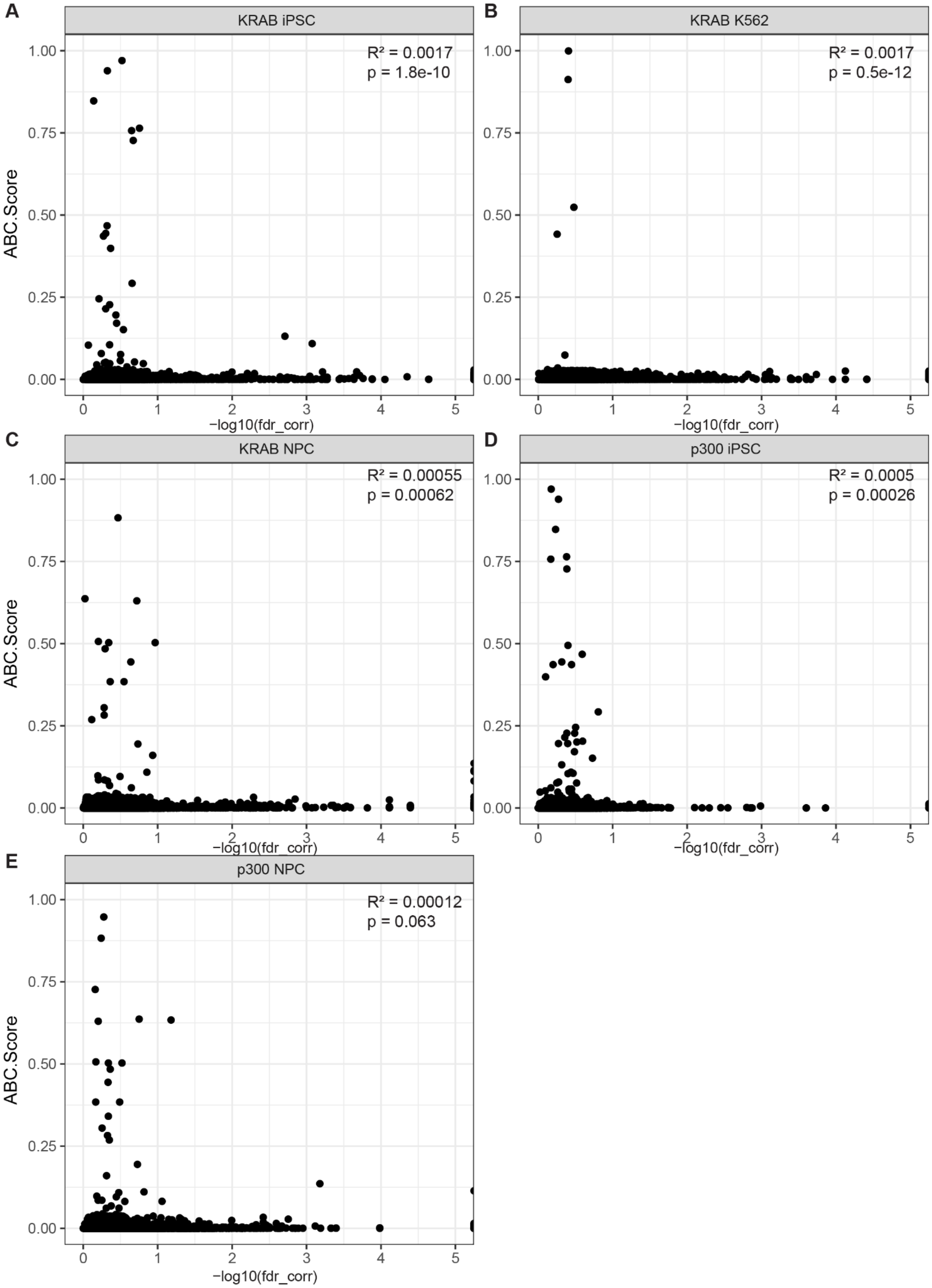


**Supplementary Figure 18. CRE-gene pair significance is not correlated with ABC Score.** Scatterplot comparing the ABC Score and the significance of all detected cCRE-gene tests in each screen for the **(A)** KRAB iPSC, **(B)** KAB K562, **(C)** KRAB NPC, **(D)** p300 iPSC, and **(E)** p300 NPC experiments. Spearman correlation and related p-value are noted in each plot.

**Supplementary Figure 19. Correlation of effect sizes for all cCRE-gene tests and cell-type specific cCRE-gene pairs.**

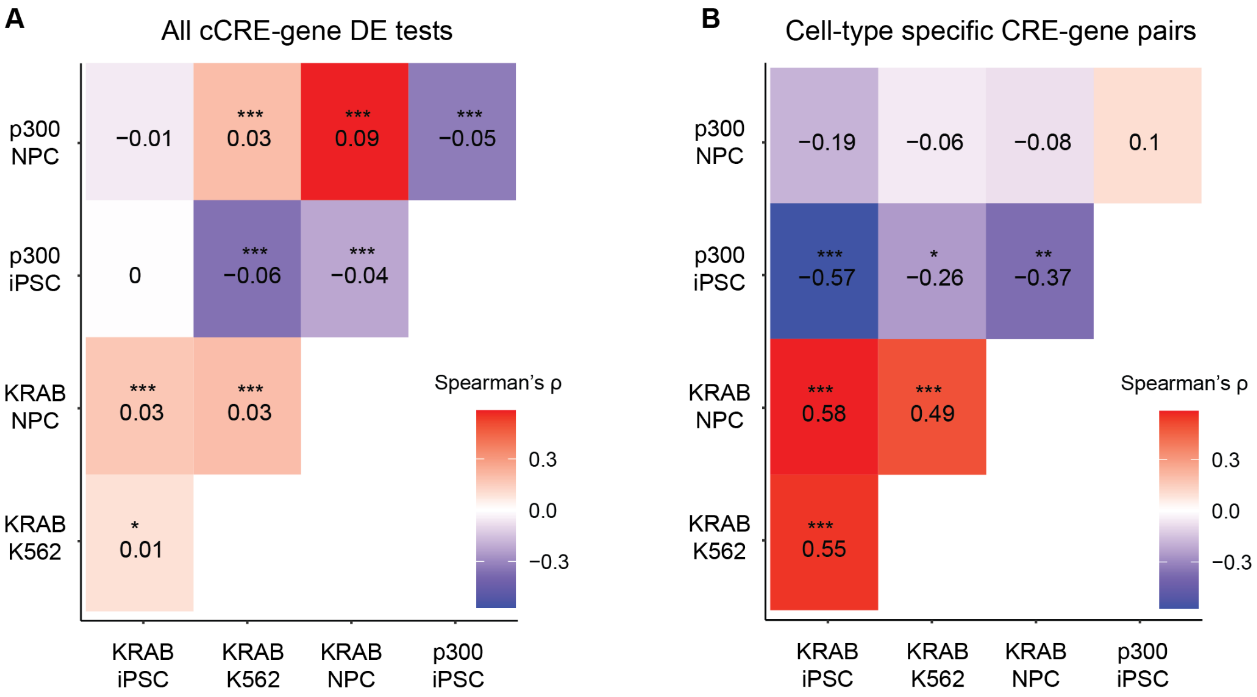


**Supplementary Figure 19. Correlation of effect sizes for all cCRE-gene tests and cell type-specific cCRE-gene pairs.** Spearman correlation of effect sizes for **(A)** all cCRE-gene tests and **(B)** all cell-type specific CRE-gene pairs across the five experiments. The shading and value noted in each cell represents Spearman correlation value. The significance is denoted as: *$p<0.05$, **$p<0.01$, ***$p<0.001$.

**Supplementary Figure 20. Basal gene expression is highly correlated between cell types.**

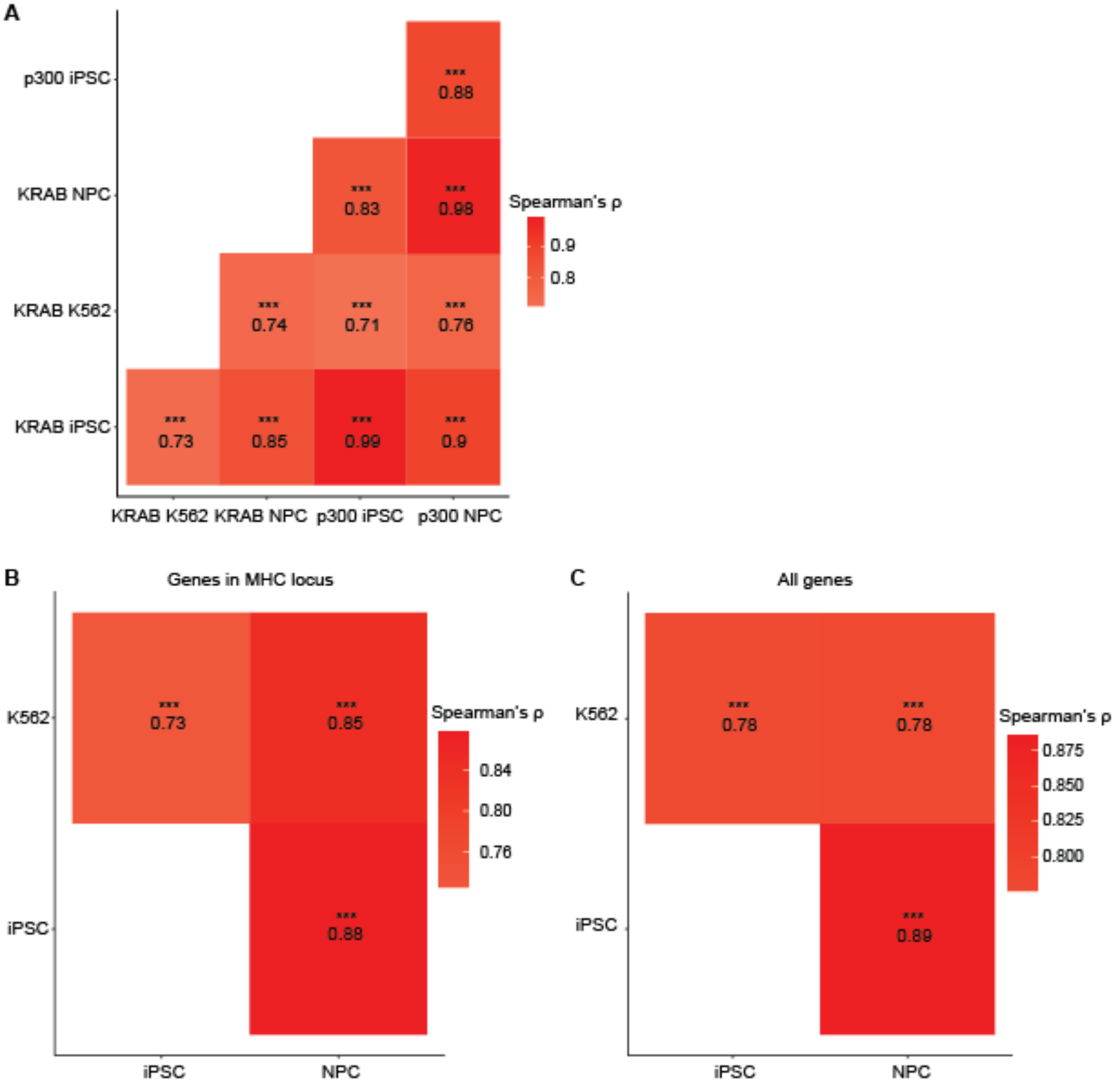


**Supplementary Figure 20. Basal gene expression is highly correlated between cell types.** Spearman correlation calculated between cell types for all genes in the MHC locus detected in the single cell screens using **(A)** normalized single cell UMI counts and **(B)** TPM values from bulk RNA-seq datasets, and **(C)** using TPM values for all genes detected in bulk RNA-seq datasets. The shading and value noted in each cell represents Spearman correlation value. The significance is denoted as: *p<0.05, **p<0.01, ***p<0.001.

**Supplementary Figure 21. Biological processes over-represented in CRE-gene pairs identified with CRISPRi.**

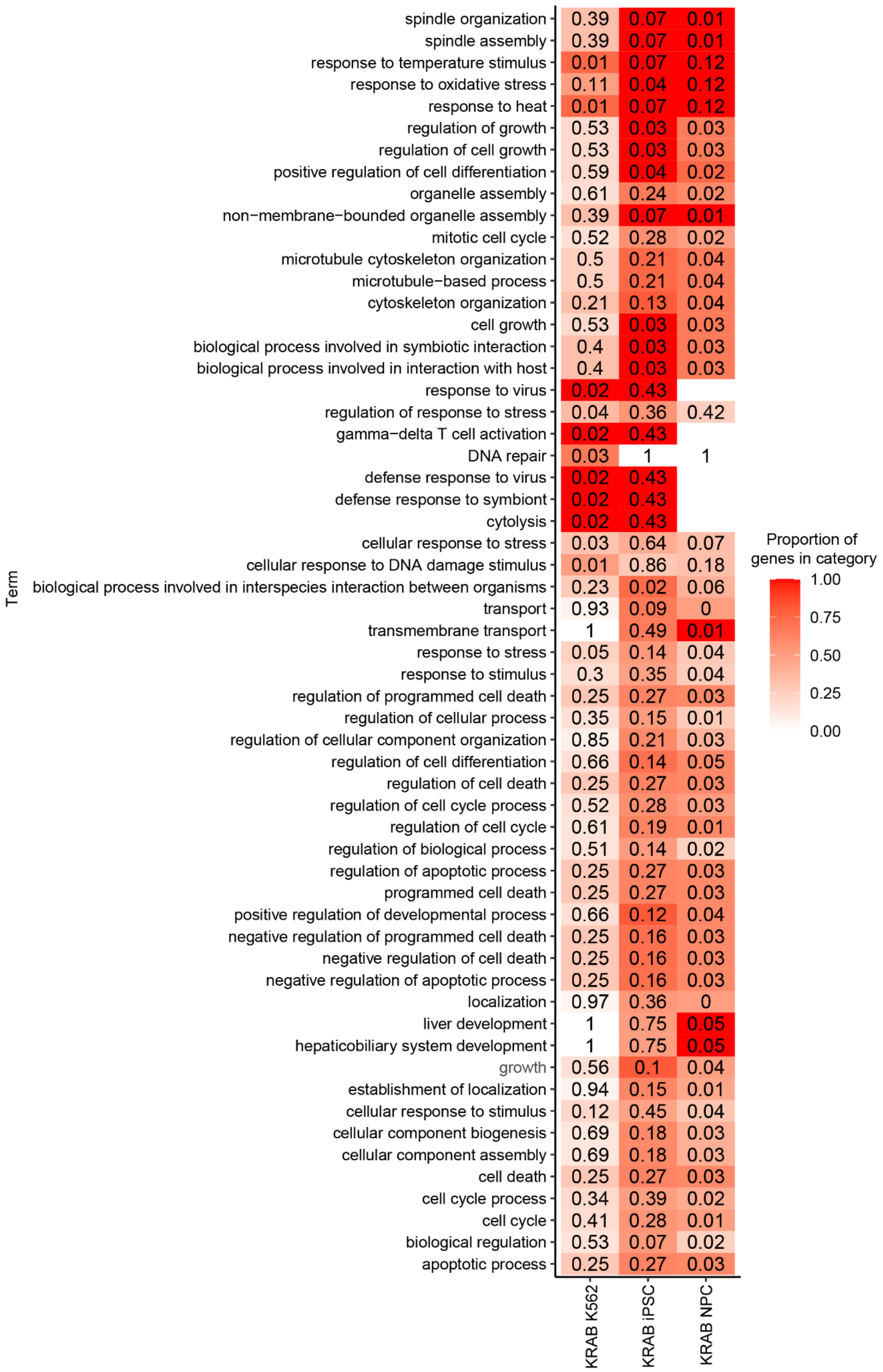

**Supplementary Figure 21. Biological processes over-represented in CRE-gene pairs identified with CRISPRi.** Heatmap of the over-represented biological processes (ranked by p-value) for genes connected to a CRE in the CRISPRi screens. The color indicates the proportion of genes connected to a CRE in the category out of the total number of genes in the category. The p-value for each term is noted in each cell.

**Supplementary Figure 22. Biological processes over-represented in CRE-gene pairs identified with CRISPRa.**

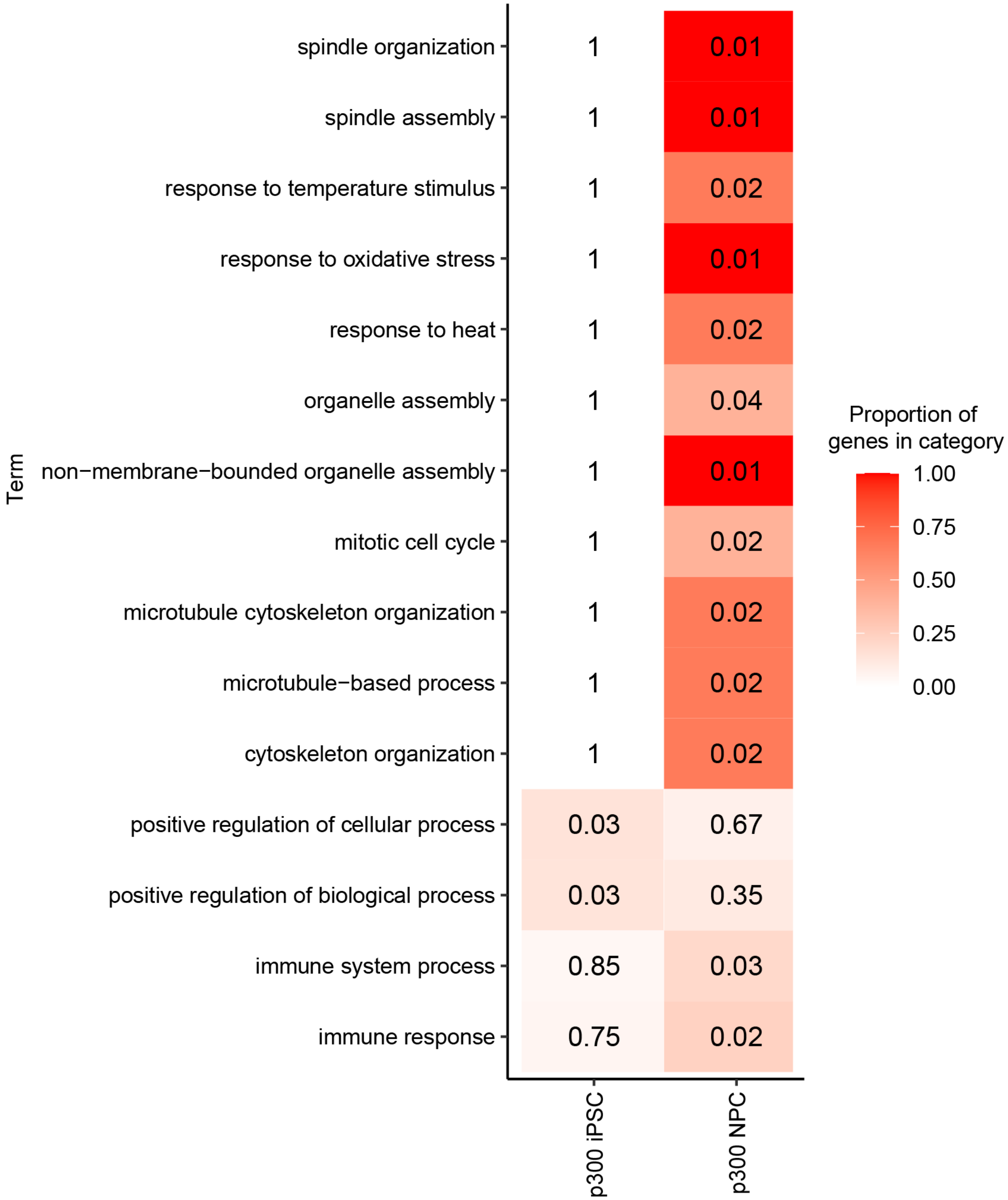


**Supplementary Figure 22. Biological processes over-represented in CRE-gene pairs identified with CRISPRa.** Heatmap of the over-represented biological processes (ranked by p-value) for genes connected to a CRE in the CRISPRa screens. The color indicates the proportion of genes connected to a CRE in the category out of the total number of genes in the category. The p-value for each term is noted in each cell.

### Supplementary Figure 23. GWAS traits without enrichment of SNPs in CREs.

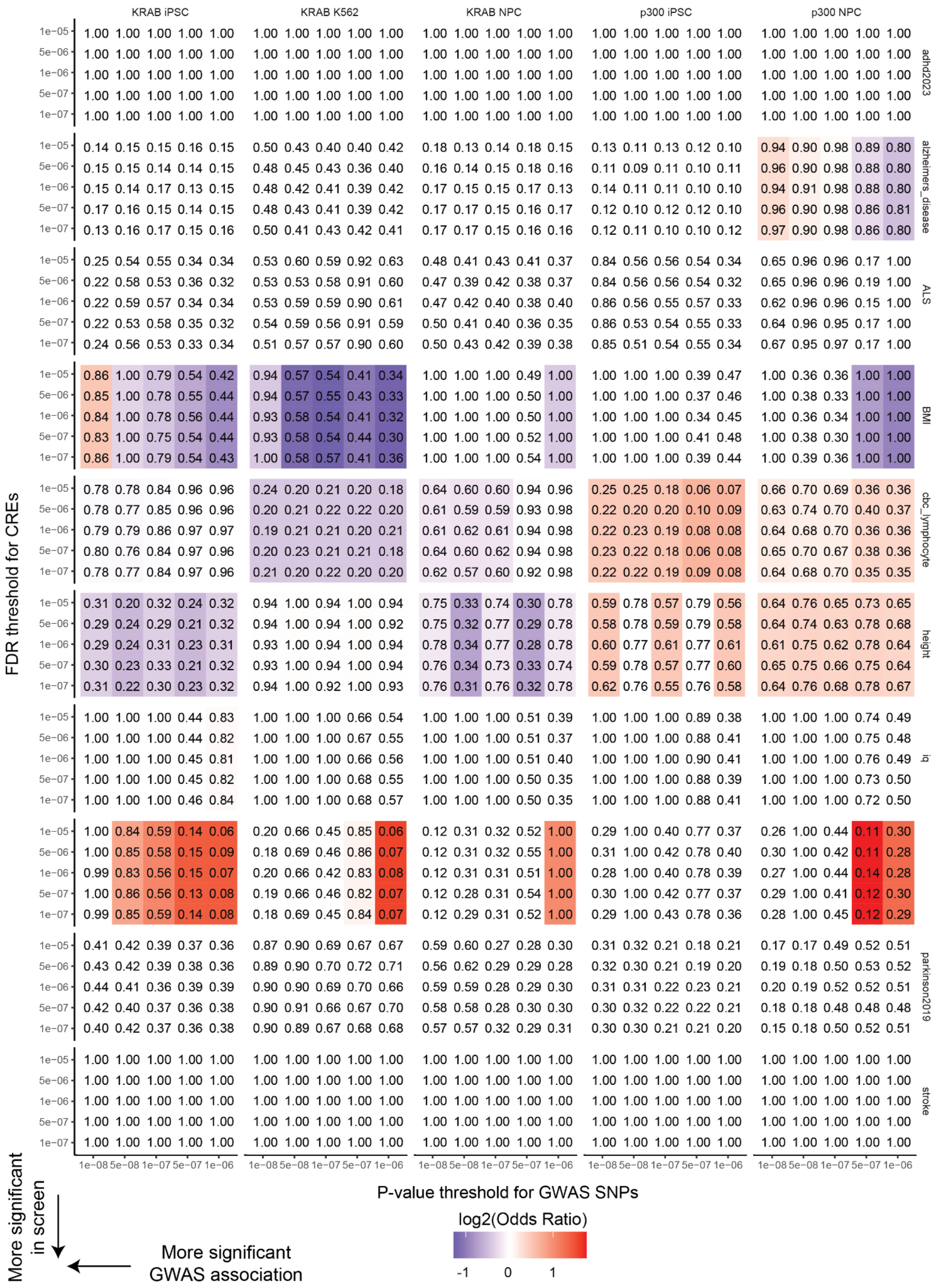

**Supplementary Figure 23. GWAS traits without enrichment of SNPs in CREs.** Heatmap of overlap of CREs with GWAS SNPs with shading indicating Odds ratio and p-values from Chi-squared test noted in each cell (**Methods**) for traits without significant enrichment (p>= 0.1; Attention-deficit/hyperactivity disorder (adhd2023), Alzheimer's disease, Amyotrophic Lateral Sclerosis (ALS), body mass index (BMI), complete blood count (CBC) lymphocyte, height, intelligence quotient (IQ), Major Depressive Disorder (mdd2019), Parkinson's disease (parkinson2019), and stroke, respectively). The x-axis indicates the significance threshold for GWAS SNPs and the y-axis indicates the significance threshold for the CREs.
